# Opposing signaling molecules exert cortical control over dissociable behaviors that tile an aggressive state

**DOI:** 10.1101/2025.03.13.643119

**Authors:** Rachel E. Gatlin, Yanfeng Du, Jordyn Gagon, Damhyeon Kwak, Sewon Park, Hailee Walker, Kathleen E. Rondem, Yang Liu, Lo Kronheim, Thomas Everett, Ashley Covington, Michaela M. Fluck, Tobias Zickmund, Larry S. Zweifel, Nicholas A. Frost, Moriel Zelikowsky

## Abstract

Neuropeptides exert broad effects across the brain to influence behavior. However, the mechanism by which neuropeptide signaling exerts top-down, cortical control over complex behavior remains poorly understood. Prolonged social isolation induces sweeping changes to behavior, including increasing aggression. Here, we find that isolation-induced aggression engages *Tachykinin-2* expressing (Tac2+) neurons in the mouse medial prefrontal cortex (mPFC). Behavioral analyses and *in vivo* recordings reveal that mPFC Tac2+ neurons are tuned to two distinct phases of an aggressive encounter: an initial phase marked by increased investigatory behaviors and a second phase comprised of increased aggression. RNA sequencing and subsequent molecule-specific loss-of-function perturbations targeting the ability of Tac2+ neurons to release either the stimulatory neuropeptide Neurokinin B or the inhibitory neurotransmitter GABA reveal that these signaling molecules exert dissociable control over increased investigation and aggression, respectively. Surprisingly, these findings reveal that the co-transmission of signaling molecules with opposing actions from the same population of cells is nevertheless able to exert coordinated control over distinct behaviors that tile an aggressive state, revealing an efficient, multiplexed neurochemical mechanism by which neuropeptidergic populations in the mPFC influence behavior.

---

Neuropeptides are thought to play a privileged role in the control of internal states. Their extended time course for signaling through G-protein coupled receptors (GPCRs), combined with the scalability of their release magnitude and diffusion over long distances leaves them well poised to modulate long-lasting states^1–6^. In *C*. *elegans,* distinct circuits comprised of antagonistic neuromodulators allow for rapid behavioral state transitions^4^. In mice, subcortical neurons utilize distinct signaling molecules to similarly guide behavioral state transitions^7^ or to drive a single specific behavior^8^. However, much of our understanding of neuropeptidergic control over internal states and behavior has been driven by studies in subcortical regions^8–11^. In contrast, little is known about the contributions of neuropeptide signaling in the cortex, where the functional role of neuropeptidergic cells remains largely understudied^12^.

The medial prefrontal cortex (mPFC), a social executive center in the brain, is thought to coordinate behavior via “top-down” projections to downstream structures^13–19^. Output of the mPFC is controlled through a careful balance of excitation and inhibition, which is highly sensitive to changes in experience, such as stress^13, 15, 20, 21^. One key source of stress known to impact the mPFC is social isolation (SI). Indeed, extended periods of social isolation have been shown to generate deleterious effects on social behavior, including increasing aggression, via changes to mPFC activity profiles^9, 22–27^. Despite this, the influence of genetically defined populations of mPFC neurons over isolation-altered behaviors remains unknown.

The Tachykinin-2 neuropeptide system has emerged as an example of distributed, neuromodulatory control over the state induced by prolonged social isolation^9^. Previous work has further implicated subcortical Tac2^+^ neurons in emotional and social behaviors^9, 28–36^. While *Tac2* and *Tacr3*, which encode the stimulatory neuropeptide Neurokinin B (NkB) and the Tac2-specific, Gq-coupled GPCR Neurokinin 3 receptor (Nk3R), respectively, are both known to be expressed in the mPFC, the prefrontal function of the Tachykinin-2 system remains unexplored in any context^35, 37^.

Here, we target the Tachykinin-2 system to interrogate the mechanism by which neuropeptides in the mPFC exert influence over complex behavior. Using behavioral analyses, single-nucleus RNA-sequencing (snRNAseq), cell-type-specific recordings and activity mapping, we identify mPFC Tac2^+^ cells as a population of unexplored GABAergic neurons whose activity is tuned to isolation-induced aggression. Molecule-specific loss-of-function perturbations targeting the ability of mPFC Tac2^+^ neurons to synthesize NkB^33, 38, 39^ or release the inhibitory neurotransmitter GABA, reveal that these signaling molecules exert dissociable control over investigatory behaviors that are increased prior to aggression and to aggressive behavior itself, respectively. Moreover, we find that local, CRISPR-mediated mutagenesis of *Tacr3* phenocopies the loss of NkB synthesis, suggesting that Nk3R-mediated signaling in the mPFC may serve to amplify the impact of Tac2^+^ neurons on behavior. Collectively, these results highlight a crucial role for neuropeptidergic signaling in the mPFC in the modulation of internal states and reveal a surprising neurochemical mechanism by which the mPFC coordinates top-down, cortical control over dissociable, state-induced behaviors via the action of opposing signaling molecules.

## RESULTS

### Prolonged social isolation induces a behavioral state characterized by increased investigation and aggression

To determine the contribution of cortical neuropeptide signaling to internal state control, we leveraged our model of chronic social isolation, which induces a state characterized by altered social behavior and increased aggression in male mice^9, 22–25, 40, 41^. Recent work suggests that prolonged^42,43,44,45^ and acute periods of social isolation in females are also sufficient to reshape social behavior^46–48^, however, isolation-induced aggression in females remains largely unstudied.

To examine the impact of prolonged social isolation across both sexes, male and female mice were isolated for two weeks and tested for aggression using the Resident Intruder (RI) assay^49^ (Fig. 1a). Compared to group housed controls (GH), socially isolated (SI) mice displayed an increase in the time spent engaged in social behaviors directed towards a novel conspecific mouse (“intruder”), including face interaction, body investigation, and anogenital investigation (Fig. 1b-d, Extended Data Fig. 1a-c, j-l)^50^. These increases occurred across investigation categories, demonstrating that isolation produces a global increase in social investigatory behaviors, rather than a re-shuffling of how these behaviors are allocated (Fig. 1e; Extended Data Fig. 1d, m). In addition, we found that SI mice showed a significant increase in aggressive behavior (Fig. 1f-g; Extended Data Fig. 1e-f, n-o), consistent with prior studies^9, 23, 34, 51^. These data demonstrate that prolonged social isolation shifts mice into a state characterized by increased investigatory behaviors and aggression, regardless of sex (Extended Data Fig. 1).

**Figure 1.**
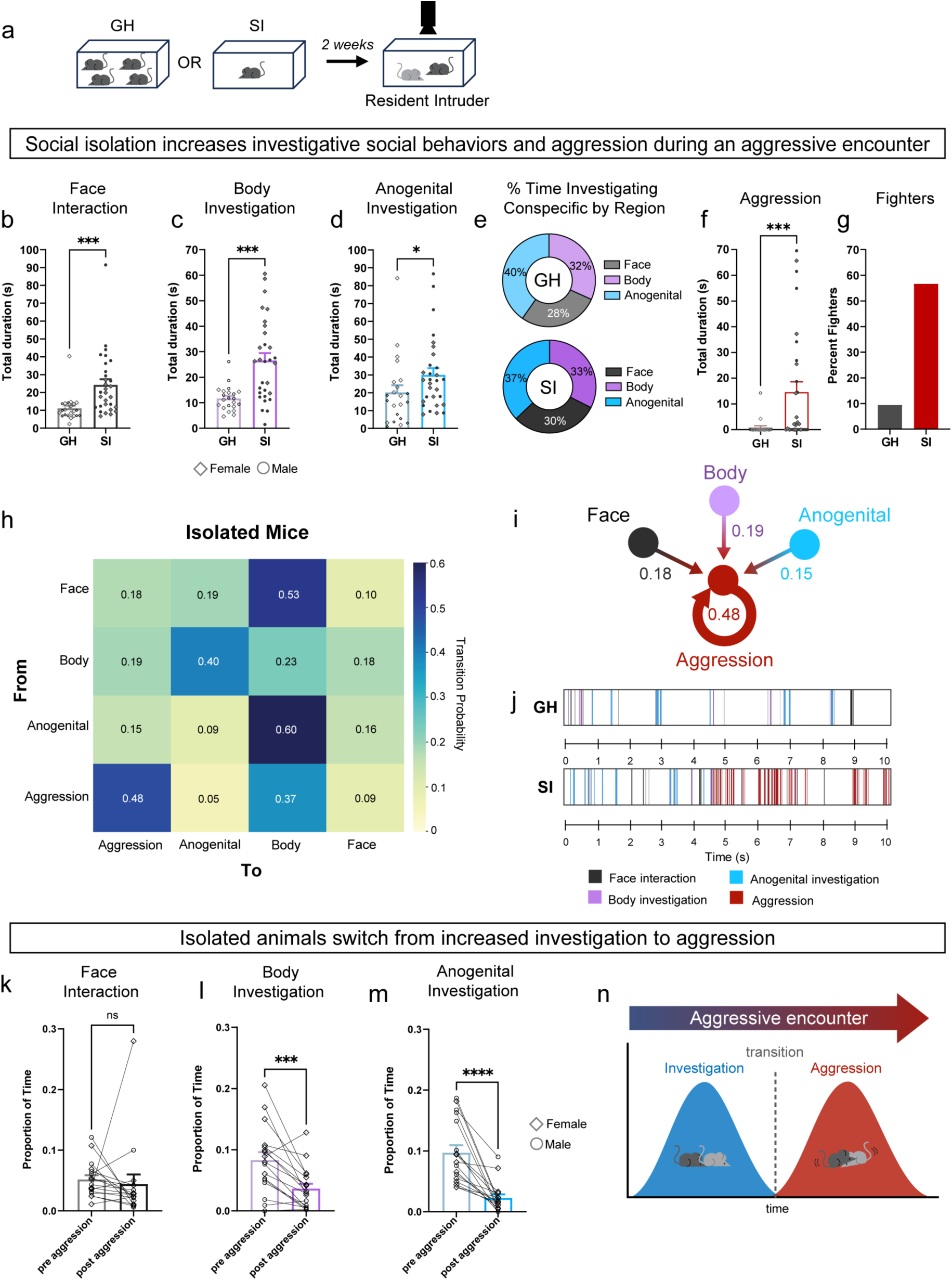
Social isolation induces a state of aggression. **a**, Experimental design. Male and female mice were group housed (GH) or socially isolated (SI) and tested on the Resident Intruder assay. **b**-**g**, n=22 GH, n=30 SI. **b**, Total duration (seconds) of face interaction in GH and SI mice, Mann-Whitney test, p = 0.0001. **c**, Total duration (seconds) of body investigation in GH and SI mice, unpaired t-test p = 0.0001, t = 4.224, df = 50. **d**, Total duration (seconds) of anogenital investigation in GH and SI mice, Mann-Whitney test p = 0.0301. **e**, Percent of total investigation time broken down by body region (face, body, anogenital) in group housed and isolated mice (total time spent investigating region/total investigation time x 100). **f**, Total duration (seconds) of aggression in GH and SI mice, Mann-Whitney test p=0.0004. **g**, Percent of animals exhibiting at least one bout of aggressive behavior (fighters). **h**, Transition matrix showing probability of isolated animal transitioning from one annotated behavior to another. **i**, Diagram summarizing behavior transition probabilities leading to aggression. **j**, Representative raster plots from a group housed and an isolated animal during the Resident Intruder assay. **k**-**m**, n=17 SI mice. **k**, Proportion of time aggressive isolated mice spent engaged in face interaction before (“pre”) and after (“post”) the first instance of aggression, Wilcoxon test, p=0.1202. **l**, Proportion of time aggressive isolated mice spent engaged in body investigation before (“pre”) and after (“post”) the first instance of aggression, paired t-test, p=0.0009. **m**, Proportion of time aggressive isolated mice spent engaged in anogenital investigation before (“pre”) and after (“post”) the first instance of aggression, Wilcoxon test, p<0.0001. **n**, Schematic representing the two phases of isolation-induced aggression: a first phase in which investigatory behaviors are increased (blue) and a second phase in which aggression is increased (red). *All bars are mean* ± *SEM, dots within bars represent mean for individual animals (diamonds=females, circles=males). ns p>0.05 * p < 0.05, ** p < 0.01, *** p < 0.001, **** p < 0.0001*.

To expand our understanding of how social isolation reshapes behavior, we generated behavioral transition matrices to determine the probability of transitioning from one behavior to the next following isolation^52–54^. SI mice were most likely to transition from one aggressive bout to another (Fig. 1h-i), indicating that once aggression begins it begets more aggression. Interestingly, raster plots visualizing behavior bouts revealed that once aggression begins, investigatory behaviors are largely shutdown (Fig. 1j), suggesting that the increase in investigation and aggression observed following social isolation may occupy dissociable temporal niches in an aggressive encounter.

To further explore whether the onset of aggression inhibits investigation behaviors in SI mice, we examined the proportion of time SI mice engaged in investigation before and after the first bout of aggression. We found that following the onset of aggression, both body and anogenital investigation – but not face interaction – were significantly reduced (Fig. 1k-m) in both sexes (Extended Data Fig. 1g-i, p-r) to levels similar to that of GH controls (Extended Data Fig. 1t-v). These data demonstrate that while some effects of prolonged social isolation may be sex specific^42, 55^, the ability of isolation to generate an internal state characterized by enhanced investigation and aggression is conserved across males and females alike. Collectively, our findings suggest that social isolation induces a state comprised of two distinct, mutually exclusive behavioral categories that tile an aggressive encounter: an initial phase in which investigatory behaviors are elevated, followed by a second phase in which aggressive behaviors are increased (Fig. 1n, Extended Data Fig. 1s).

### Prefrontal Tac2^+^ neurons are tuned to isolation-induced aggression

Recent work has implicated the mPFC in the control of aggression^22, 42, 56, 23, 57, 58^ but the neurochemical mechanism within the cortex that enables social isolation to induce an aggressive state is unknown. To understand whether isolation-induced aggressive encounters engage mPFC Tac2^+^ neurons, we assessed whether Tac2^+^ cells were activated following the RI assay. Using fluorescent in situ hybridization (*FISH*, RNAscope™) to probe for *Tac2* and the activity-dependent immediate early gene *cFos,* we found a significant increase in the percent of Tac2^+^ cells expressing *cFos* in SI compared to GH mice (Fig. 2a-c). This increase was specific to Tac2^+^ cells and could not be accounted for by an overall increase in *cFos* expression due to isolation (Extended Data Fig. 2a-b).

**Figure 2.**
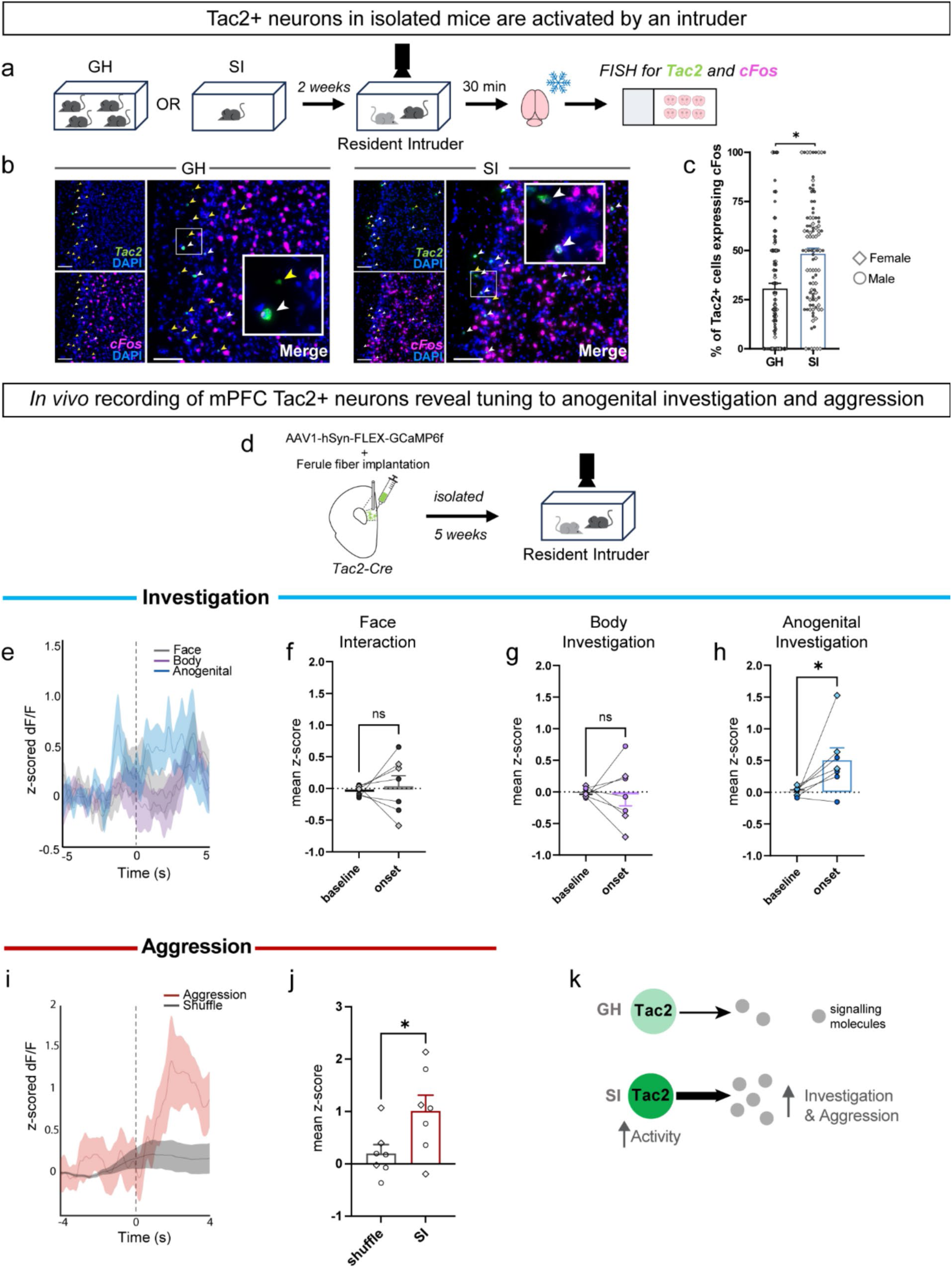
Prefrontal Tac2^+^ neuron activity is tuned to isolation-induced aggression. **a,** Experimental design. Mice were isolated (SI) or group housed (GH) and tested on the Resident Intruder assay. Brains were then extracted and processed for fluorescent in situ hybridization (FISH) to visualize *cFos* and *Tac2* mRNA. **b**, Representative images depicting *Tac2* (green) and *cFos* (magenta) expression in a group housed animal (left) and an isolated animal (right), scale bar represents 100 µm. White arrows indicate cells with *Tac2* and *cFos* expression, yellow arrows indicate cells with only *Tac2* expression. **c**, Percent of Tac2^+^ cells co-expressing *cFos*, N=16 mice (n=8 GH mice, n=8 SI mice), ∼4 sections/animal, two-tailed nested t-test p = 0.0267, t=2.475, df=14. **d**, Experimental design for fiber photometry experiments. **e**, Behavior-triggered neural activity trace of mPFC Tac2^+^ neurons (z-scored dF/F) for face interaction, body investigation and anogenital investigation bouts prior to first bout of aggression. **f**, Mean z-scored dF/F for face interaction comparing baseline period (-4 to -2s before behavior onset) to post behavior onset period (1 to 3s post onset), paired t-test, p=0.6367, t=0.4973, df=6. **g**, Mean z-scored dF/F for body investigation comparing baseline period to post behavior onset period, paired t-test, p=0.8765, t=0.1622, df=6. **h**, Mean z-scored dF/F for anogenital investigation comparing baseline period to post behavior onset period, paired t-test, p=0.0455, t=2.516, df=6. **i**, Behavior-triggered neural activity trace of mPFC Tac2^+^ neurons (z-scored dF/F) for aggression. **j**, Mean z-scored dF/F for aggression in isolated (SI) mice compared to circularly permutated data from the same animals (shuffle) post behavior onset, paired t-test, p=0.0424, t=2.570, df=6. N=7 mice. **k**, Model depicting that social isolation increases the activity of mPFC Tac2^+^ neurons and leads to increased investigatory behaviors and aggression. *All bars are mean* ± *SEM, dots within bars represent mean for individual animals (diamonds=females, circles=males). ns p>0.05 * p < 0.05, ** p < 0.01, *** p < 0.001, **** p < 0.0001*.

As *cFos* expression cannot disentangle which behaviors Tac2^+^ cells were activated during, we performed fiber photometry recordings to understand the *in vivo* activity of mPFC Tac2^+^ neurons in SI mice during the RI assay (Fig. 2d). Tac2-IRES-Cre mice^9, 59^ were injected with a virus encoding a Cre-dependent calcium indicator (GCaMP6f) and implanted with a ferule fiber targeting the mPFC to enable *in vivo* bulk recordings of Tac2^+^ population activity following isolation (Extended Data Fig. 2c). Calcium activity during the RI assay was recorded and aligned to all four social behaviors: face interaction, body investigation, anogenital investigation and aggression. Interestingly, when we examined calcium activity during investigatory behaviors (Fig. 2e-h), we found that Tac2^+^ neural activity (dF/F) was significantly increased during anogenital investigation occurring prior to the first bout of aggression (Fig. 2h), but not after aggression (Extended Data Fig. 2f). In contrast, we did not see significant tuning of Tac2^+^ cells to face interaction or body investigation (Figure 2f-g, Extended Data Fig. 2d-e), suggesting that these behaviors may be encoded by an alternate population of isolation-sensitive cells.

To assess the response profile of these cells during aggression, we examined the activity of mPFC Tac2^+^ neurons in isolated animals during bouts of aggressive behavior (Fig. 2i-j; Extended Data Fig. 2g). Tac2^+^ neurons showed an increase in their activity triggered by the onset of aggression, demonstrating that these cells are tuned to isolation-induced aggression. Consistent with prior work conducting fiber photometry recordings during aggression^60, 61^, this signal ramps up with time, suggesting that as an aggression bout progresses, the activity of mPFC Tac2^+^ neurons increases accordingly. Collectively, these data suggest that social isolation induces an increase in the activity of mPFC Tac2^+^ cells during anogenital investigation and aggression (Fig. 2k).

### Prefrontal Tac2^+^ cells represent a unique subclass of GABAergic neurons

Previous research has found that social isolation decreases the spiking activity of pyramidal neurons in the PFC during aggression^42^ and increases mPFC GABAergic neuron activity^23, 62^. We found that isolation increases the activity of mPFC Tac2^+^ neurons (Fig. 2). To characterize the genetic profile of *Tac2*-expressing cells in the mPFC, we used single-nucleus RNA sequencing (snRNAseq) as well as fluorescent in situ hybridization (*FISH*, RNAscope)^63^. Our snRNAseq analyses revealed that *Tac2* is predominantly expressed in inhibitory neuronal populations, (Fig. 3a-d), consistent with findings in other cortical regions^39, 64, 65, 66^. In contrast, *Tac2* is minimally expressed in glia (Fig. 3a-d). Interestingly, cells which contain low levels of *Tac2* represent a more heterogenous population, including expression of glutamatergic cell cluster types, while those expressing medium or high levels of *Tac2* are almost exclusively GABAergic (Extended Data Fig. 3a-c). *FISH* experiments revealed that Tac2^+^ cells represent a largely GABAergic population (*Gad1/Gad2^+^*), with virtually no overlap with glutamatergic (*Slc17a7^+^*) cells (Extended Data Fig. 3d-f).

**Figure 3.**
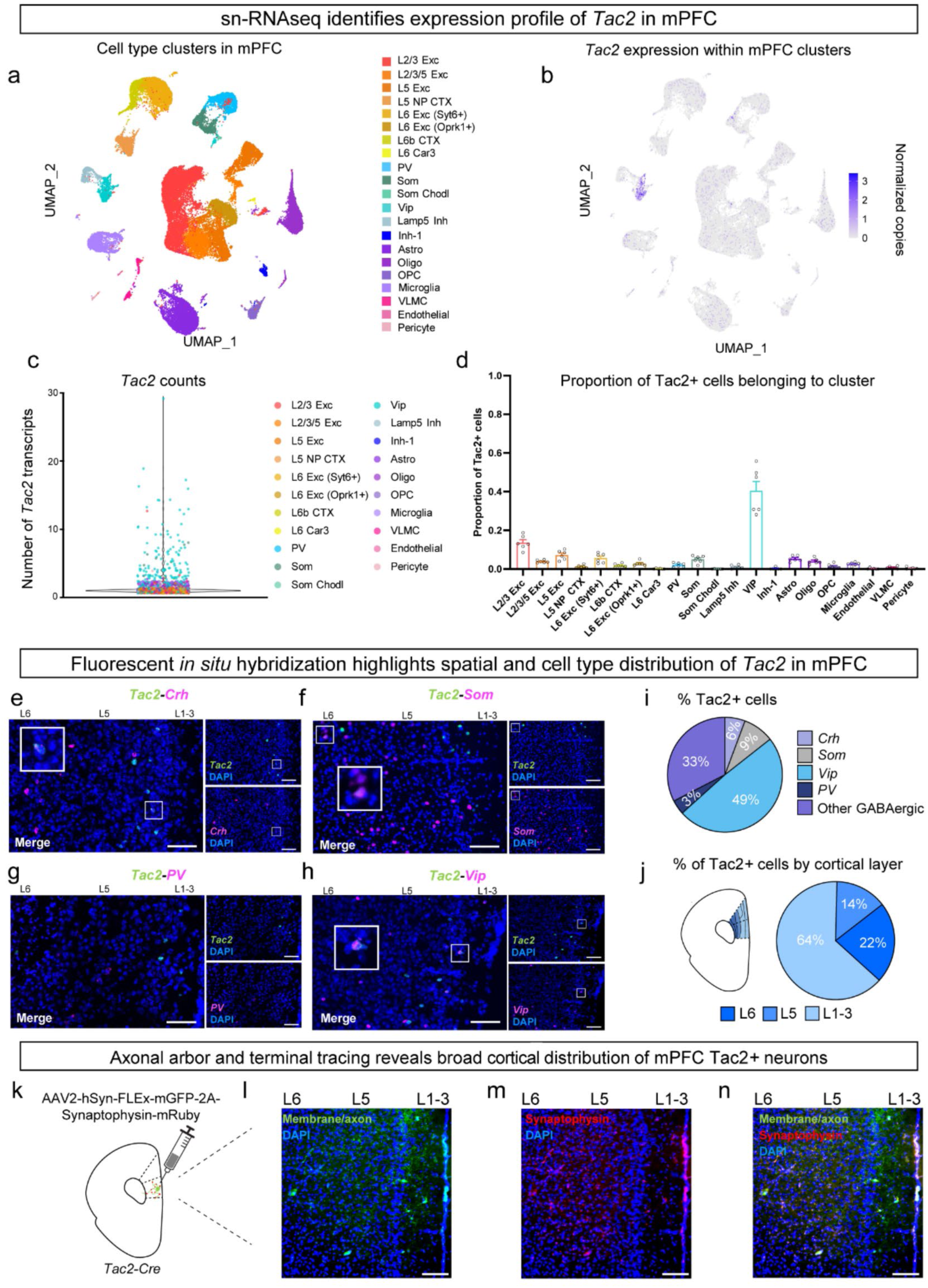
Prefrontal Tac2^+^ neurons represent an unexplored population of GABAergic neurons. **a-d,** snRNAseq analyses, N=6 mice**. a**, UMAP plot showing clusters of defined cortical cell types (50,020 cells). **b**, UMAP plot showing *Tac2* expression within each defined cortical cell type cluster (223.67 ± 26.49 Tac2^+^ cells/sample). **c,** Volcano plot depicting the number of *Tac2* transcripts (normalized) in each cell. **d**, Proportion of total *Tac2*-expressing cells in mPFC belonging to each cell cluster. **e-j**, Fluorescent in situ hybridization (RNAscope™), N=8 mice. **e-h**, Representative images of FISH, scale bar represents 50µm. *Tac2*: Tachykinin-2, *Crh*: corticotropin releasing factor, *Som*: somatostatin, *PV*: parvalbumin, *Vip*: vasoactive intestinal peptide. Inset boxes (white) highlight co-labelled cells. **i**, Pie chart showing percent of Tac2^+^ cells co-expressing *Crh*, *Som, PV*, or *Vip*. **j**, Diagram showing prelimbic and infralimbic regions of medial prefrontal cortex with cortical layers. L1-3: layers 1-3, L5: layer 5, L6: layer 6. Pie chart showing the percent of Tac2^+^ cells in cortical layers. **k**, Depiction of viral infusion into mPFC. **l**, Representative image showing Tac2^+^ neuronal axons labelled with GFP across cortical layers. **m**, Representative image of Tac2^+^ neuronal terminals (synaptophysin) in red (mRuby) across cortical layers. **n**, Merged image showing axonal arbor (green) and terminals (red) of Tac2^+^ neurons. **l-n**, scale bars represent 100µm. *All bars are mean* ± *SEM, dots within bars represent mean for individual animals*.

Our *FISH* results identified Tac2^+^ neurons as a mix of distinct, genetically defined GABAergic cell types, including those that co-express vasoactive intestinal peptide (*Vip)*, corticotrophin releasing hormone (*Crh*), somatostatin (*Som*), and parvalbumin (*PV*) (Fig. 3e-h). The gene most likely to be co-expressed with *Tac2* was vasoactive intestinal peptide (*Vip*), which accounted for roughly half of Tac2^+^ neurons, as visualized using *FISH* (Fig. 3i). *FISH* also allowed us to assess the spatial profile of *Tac2* expression, where we found that 64% of Tac2^+^ neurons reside in layers 1-3 of the cortex, where they overlap with *Vip* by 60%, while the remaining 36% of Tac2^+^ neurons reside in deeper layers (5 and 6), where they overlap with *Vip* by 40% (Fig. 3j; Extended Data Fig. 3g-h). Interestingly, we were not able to identify a single subtype of mPFC Tac2^+^ neuron that was uniquely activated by an aggressive encounter (Extended Data Fig. 4a-t), suggesting that isolation-induced increases in Tac2^+^ cell activity cannot be explained by the increased activity of a particular cell-type subclass. Collectively, these data suggest that mPFC Tac2^+^ neurons represent a genetically unique class of neurons, predominantly localized to early cortical layers, that are activated by isolation-induced aggressive encounters.

To characterize the anatomical features of Tac2^+^ neurons in the mPFC, we visualized the axonal arbor of these cells using anterograde tracing. We injected a Cre-dependent adeno-associated virus (AAV) expressing membrane-bound GFP and synaptophysin fused to mRuby^67^ into the mPFC of Tac2-IRES-Cre mice (Fig. 3k). This tracing revealed that Tac2^+^ cell bodies which primarily reside in layers 1-3 have axonal arbors extending into layer 5 and terminals spread throughout all cortical layers, with the highest density in layer 5 (Fig. 3l-n). These results illustrate that, though sparse, this class of neurons has the potential to reach broadly across the mPFC to alter social behavior, consistent with the idea that a small handful of neurons is sufficient to maintain a representation of a continuous variable important for guiding behavior^68^.

### Prefrontal Tac2^+^ neurons are necessary for isolation-induced aggression

Given that Tac2^+^ neurons in the mPFC are tuned to isolation-induced aggression (Fig. 2), we next tested whether mPFC Tac2^+^ neurons are necessary for isolation-induced aggression by performing a series of loss-of-function perturbations and behavior testing. Because we discovered that mPFC Tac2^+^ neurons are GABAergic (Fig. 3) and are therefore capable of releasing both NkB, the stimulatory neuropeptide encoded by *Tac2*^39, 69–71^ and GABA, the inhibitory neurotransmitter, we selectively targeted Tac2^+^ neurons or their ability to synthesize and/or release either signaling molecule, and examined behavior (Fig. 4a-b). To first determine whether Tac2^+^ neurons are required for SI-aggression, mPFC Tac2^+^ neurons were ablated in isolated mice using a virally encoded, Cre-dependent caspase-3 (executioner caspase) infused into the mPFC of Tac2-IRES-Cre mice (Fig. 4c, Extended Data Fig. 5i-k). Ablation of mPFC Tac2 neurons significantly reduced anogenital investigation and aggression, while leaving face interaction and body investigation unchanged (Fig. 4d-h), Extended Data Fig. 5a-h), consistent with our fiber photometry data demonstrating that Tac2^+^ cells are tuned to anogenital investigation and aggression. Importantly, silencing of stress-sensitive Crh^+^ mPFC cells – a population of sparsely expressed GABAergic neurons that show little overlap with *Tac2* (Fig. 3e, i) – had no effect on anogenital investigation or aggression (Extended Data Fig. 6), suggesting that our Tac2^+^ cell ablation results are not simply due to the inhibition of any small population of neurons in the mPFC. Collectively, these data reveal that Tac2^+^ neurons are specifically necessary for isolation-induced investigation and aggression (Fig. 4h).

**Figure 4.**
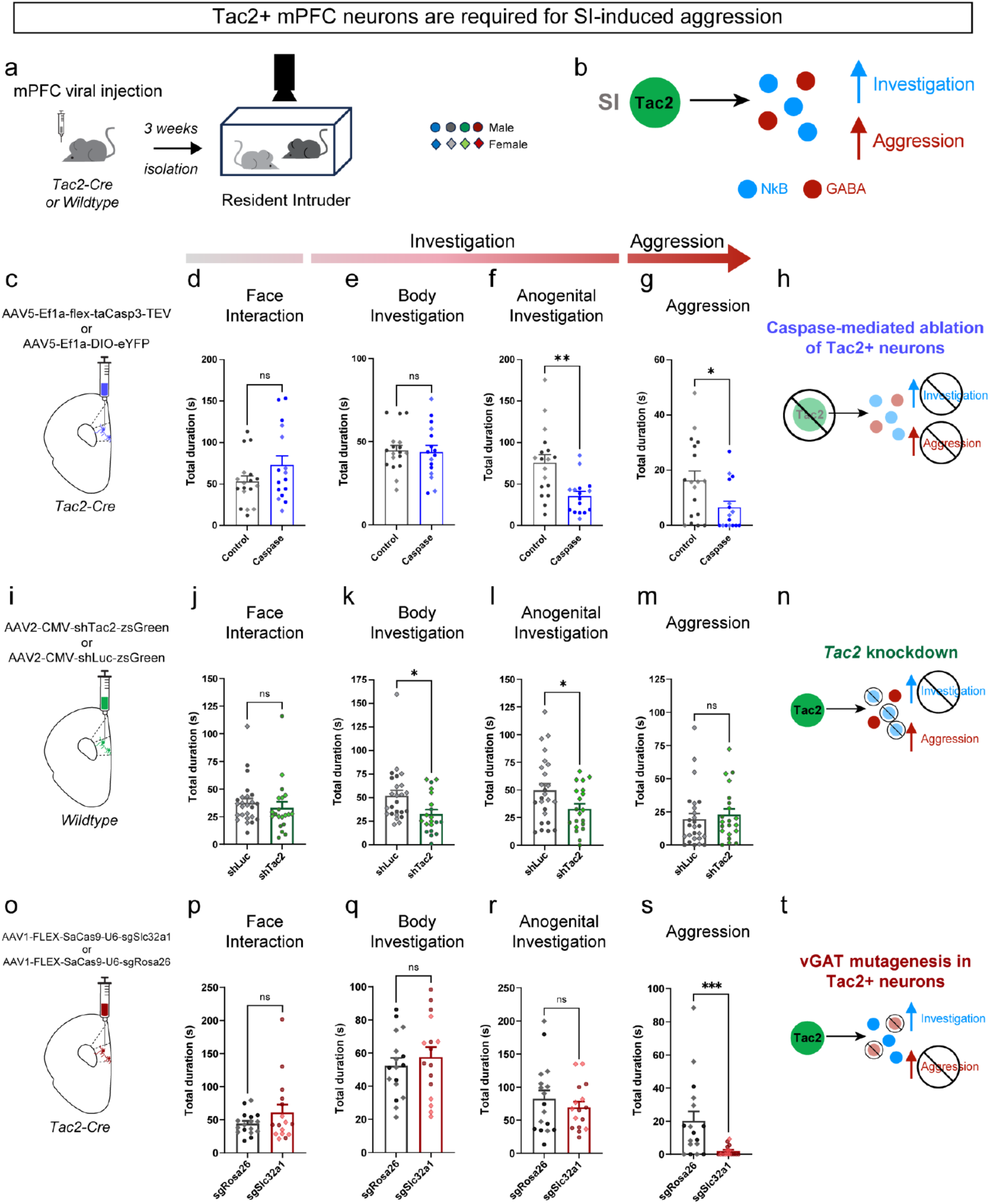
Opposing signalling molecules control increased investigation and aggression. **a**, Experimental design depicting approach for viral manipulations of mPFC Tac2^+^ neurons followed by behavioral testing on the Resident Intruder assay. **b**, Model of isolation-induced aggressive state and the role of mPFC Tac2^+^ neurons. **c**, Experimental approach for virally-mediated ablation of mPFC Tac2^+^ neurons. **d-g**, n=18 eYFP control, n=16 caspase. **d**, Total duration (seconds) of face interaction in control and caspase mice, Mann-Whitney test p=0.2514. **e**, Total duration (seconds) of body investigation in control and caspase mice, unpaired t-test p=0.8567, t=0.1820, df=32. **f** Total duration (seconds) of anogenital investigation in control and caspase mice, unpaired t-test p=0.0013, t= 3.528, df=32. **g**, Total duration (seconds) of aggressive behavior in control and caspase mice, Mann-Whitney test, p=0.0455. **h**, Diagram summarizing the impact of mPFC Tac2^+^ neural ablation in reducing SI-investigation and aggression. **i**, Experimental approach for virally-mediated knockdown of *Tac2* in mPFC. **j-m,** n=25 luciferase, n=20 shTac2. **j**, Total duration of face interaction (seconds) in Luciferase shRNA mice (shLuc) and *Tac2* shRNA mice (shTac2), Mann-Whitney test: p=0.2810. **k**, Total duration (seconds) of body investigation in Luciferase shRNA mice (shLuc) and *Tac2* shRNA mice (shTac2), unpaired t-test, p = 0.0146, t = 2.549, df = 43. **l,** Total duration (seconds) of anogenital investigation in Luciferase shRNA mice (shLuc) and *Tac2* shRNA mice (shTac2), unpaired t test: p = 0.0359, t = 2.166, df = 43. **m**, Total duration (seconds) of aggression in Luciferase shRNA mice (shLuc) and *Tac2* shRNA mice (shTac2), Mann-Whitney test, p = 0.3808. **n**, Diagram summarizing the impact of *Tac2* knockdown/Neurokinin B (NkB) signalling disruption in reducing SI-investigation but not aggression. **o**, Experimental approach for virally-mediated, Cre-dependent CRISPR/SaCas9 mutagenesis of *Slc32a1* (vGAT) in Tac2-IRES-Cre mice to prevent GABA transmission specifically from mPFC Tac2^+^ neurons. **p**-**s**, n= 17 *sgRosa26*, n=16 sg*Slc32a1.* **p**, Total duration (seconds) of face interaction in mice with guide RNA directed to the Rosa26 locus (sg*Rosa26*) or the vGAT locus (sg*Slc32a1*), Mann-Whitney test: p = 0.8173. **q**, Total duration (seconds) of body investigation in mice with guide RNA directed to the Rosa26 locus (sg*Rosa26*) or the vGAT locus (sg*Slc32a1*), unpaired t test: p = 0.4968, t = 0.6876, df = 31. **r**, Total duration (seconds) of anogenital investigation in mice with guide RNA directed to the Rosa26 locus (sg*Rosa26*) or the vGAT locus (sg*Slc32a1*), unpaired t test: p = 0.4085, t = 0.8379, df = 31. **s**, Total duration (seconds) of aggression in mice with guide RNA directed to the *Rosa26* locus (sg*Rosa26*) or the vGAT locus (sg*Slc32a1*), Mann-Whitney test: p = 0.0009. **t,** Model summarizing the impact of *Slc32a1* (vGAT) mutagenesis in Tac2^+^ neurons in attenuating SI-aggression but not investigation. *Bars represent mean ± SEM, dots within bars represent mean behavior duration for each animal (diamonds=females, circles=males). ns p > 0.05, * p < 0.05, ** p < 0.01, *** p < 0.001*.

### *Tac2* knockdown in mPFC attenuates isolation-enhanced investigation

While we identified a necessary role for mPFC Tac2^+^ neurons in controlling isolation-induced investigation and aggression (Fig. 4c-h), we next sought to determine the neurochemical mechanism underlying this effect. To examine the contributions of NkB signaling to isolation-induced behaviors, we took an shRNAi approach to knock down *Tac2* in the mPFC of isolated mice (Fig. 4i, Extended Data Fig. 7)^9^. Wildtype mice were infused with a virally mediated shRNA targeting *Tac2* (shTac2) or a control shRNA targeting luciferase (shLuc) into the mPFC and tested on the RI assay^9^ (Fig. 4a, i). Knockdown of *Tac2* in the mPFC of isolated mice resulted in a significant decrease in anogenital investigation (Fig. 4j-l), consistent with our ablation experiment (Fig. 4f). However, in contrast, *Tac2* knockdown had no impact on aggression (Fig. 4m). These data reveal that NkB signaling contributes to the effect of isolation to increase investigation but not aggression (Fig. 4n, Extended Data Fig. 7a-h).

### Disruption of GABA release from mPFC Tac2^+^ neurons attenuates isolation-induced aggression

Our loss-of-function results demonstrate that mPFC Tac2^+^ neurons are required for isolation-induced aggression (Fig. 4h) but that this cannot be explained by release of NkB from these cells (Fig. 4n). To determine whether GABA transmission from mPFC Tac2^+^ neurons mediates SI-aggression, we used a CRISPR-based approach to generate loss-of-function mutations in *Slc32a1*, the gene encoding the vesicular GABA transporter (vGAT), in mPFC Tac2^+^ neurons. An AAV encoding a Cre-dependent Staphylococcus aureus Cas9 (SaCas9) and guide RNA (sgRNA) directed to *Slc32a1*^72^ was injected into the mPFC of Tac2-IRES-Cre mice alongside a Cre-dependent KASH-eGFP virus^73^ (Fig. 4a, o) to induce nonsense mutations of *Slc32a1* in Tac2^+^ mPFC neurons and visualize the viral infusion site (Extended Data Fig. 8). A similar AAV cocktail encoding the same Cre-dependent Staphylococcus aureus Cas9 (SaCas9) but with a guide RNA targeting the Rosa26 locus^74^ was injected in control animals (Fig. 4o). Animals with the CRISPR-mediated mutation of *Slc32a1* showed no changes in investigative behaviors (Fig. 4p-r, Extended Data Fig. 8a-c, e-g). However, isolation-induced aggression was significantly reduced in these mice (Fig. 4s, Extended Data Fig. 8d, h). This reduction in aggressive behavior, with no impact on investigation, reveals that GABA transmission from mPFC Tac2^+^ neurons controls SI-aggression (Fig. 4t), whereas NkB signaling from the same population of neurons controls SI induced increases in investigatory behavior (Fig. 4n). Collectively, these findings point to dissociable roles for NkB and GABA in the control of isolation induced investigation and aggression, respectively.

Interestingly, in contrast to our caspase-mediated cell ablation effects (Fig. 4h), DREADD-mediated inhibition of mPFC Tac2^+^ neurons via administration of the DREADDs activating ligand deschloroclozapine (DCZ) (0.1 mg/kg, i.p., Extended Data Fig. 9a-p) delivered at test reduced aggression only (Extended Data Fig. 9q-w). This may reflect differences in approach, as cell ablation eliminates neuropeptide release while DREADDs-mediated inhibition may not^75, 76^. Alternatively, this discrepancy may reflect differences regarding the time scale of inhibition: cell ablation eliminates Tac2^+^ neuron activity – and the release of both NkB and GABA – across the entire window of isolation as well as at test, while acute DREADDs inhibition only limits the function of Tac2^+^ neurons at test. Indeed, consistent with this idea, we discovered that NkB levels in the mPFC increased following prolonged isolation (Extended Data Fig. 10), in line with what we previously found in subcortical structures^9^. These data suggest that the neuropeptide NkB is being chronically released across the window of isolation, which may be critical for the effects of isolation-induced changes in behavior that we see at test.

### Prefrontal Tacr3^+^ cells represent a genetically heterogenous population

Our data reveal that Tac2/NkB signaling is critical for isolation-induced investigatory behavior (Fig. 4n). To determine whether NkB exerts its effects via action at local Nk3Rs in mPFC, we interrogated the function of mPFC Tacr3^+^ cells. We first sought to establish the genetic identity of Tacr3^+^ cells in the mPFC, as they have never been previously explored. Using our snRNAseq dataset (Fig. 3), we found that *Tacr3,* the gene encoding Nk3R, is sparsely expressed across a number of distinct cell types, including excitatory and inhibitory neuron populations as well as in glia (Fig. 5a-d). Indeed, while *Tacr3*-expressing cells are predominantly excitatory neurons – consistent with their broader identity across the cortex^39, 65, 66, 77^ – we nevertheless found that they also represent a sparse, mixed group of GABAergic cells labelled by the interneuron markers *PV*, *Som* or *Vip*. Importantly, *Tac2* and *Tacr3* were not co-expressed (Extended Data Fig. 11a), suggesting that Tac2^+^ neurons exert their influence on a separable population of *Tacr3*-expressing cells.

**Figure 5.**
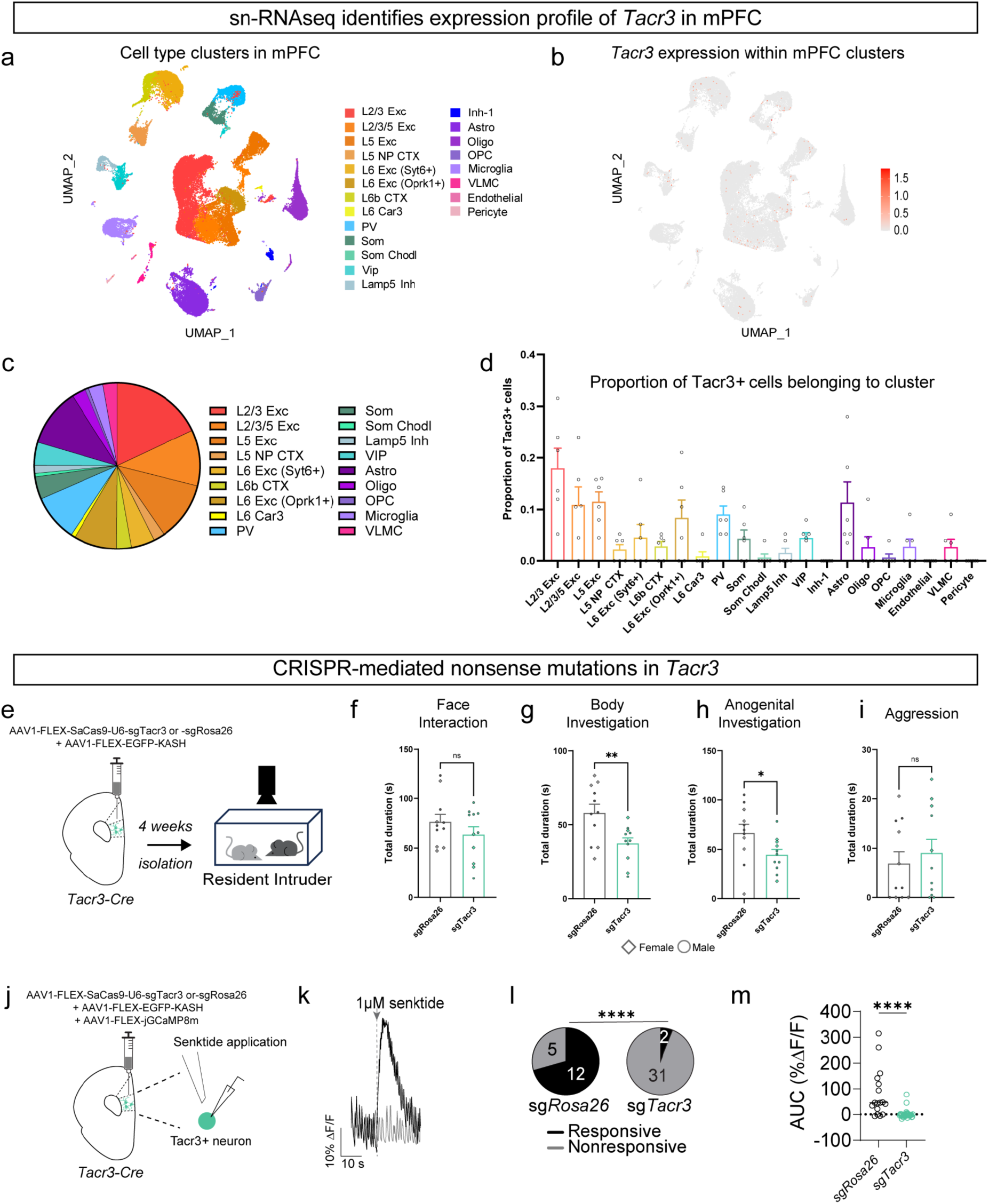
Prefrontal Tacr3^+^ cells represent a genetically heterogenous population which amplify the effects of Tac2/NkB signalling on isolation-induced investigation. **a-d,** snRNAseq analyses, N=6 mice. **a**, UMAP plot of cell type clusters in mPFC. **b**, *Tacr3* expression within mPFC clusters (23 ± 1.48 cells/sample). **c**, Pie chart summarizing the proportion of total *Tacr3* expression in mPFC by cell cluster type. **d**, Proportion of total *Tacr3*-expressing cells in mPFC belonging to each cell cluster. **e**, Experimental design. Tacr3-Cre mice were injected with sg*Rosa26* or sg*Tacr3*, isolated, and tested for behavior on the Resident Intruder assay. **f-i**, n=11 sg*Rosa26*, n=11 sg*Tacr3*. **f**, Total duration (seconds) of face interaction in sg*Rosa26* and sg*Tacr3* mice, unpaired t-test, p = 0.2525, t = 1.178, df = 20. **g**, Total duration (seconds) of body investigation in sg*Rosa26* and sg*Tacr3* mice, unpaired t-test, p = 0.0439, t = 2.121, df = 20. **h**, Total duration (seconds) of anogenital investigation in sg*Rosa26* and sg*Tacr3* mice, unpaired t-test, p = 0.0080, t = 2.944, df = 20. **i**, Total duration (seconds) of aggression in sg*Rosa26* and sg*Tacr3* mice, Mann-Whitney test, p = 0.4634. **j-m**, Validation of CRISPR/SaCas9-mediated mutagenesis of mPFC *Tacr3* cells. **j**, Tacr3-Cre mice were injected with a Cre-dependent GCaMP and either CRISPR/SaCas9 viruses directed towards *Rosa26* or *Tacr3.* Slices were taken and prepared for calcium imaging. Changes in fluorescence (dF/F) were measured in response to senktide (Nk3R agonist) application. **k**, Representative dF/F GCaMP trace from a cell responsive to senktide (black) and a non-responsive cell (grey). **l**, Proportion of cells in sg*Rosa26* (left) and sg*Tacr3* (right) mice responsive (black) or not responsive (grey) to senktide, Fisher’s exact test, p<0.0001. **m**, Area under the curve (percent change in fluorescence) of GCaMP signal between sg*Rosa26* and sg*Tacr3* mice in response to senktide, Mann-Whitney test, p<0.0001, lines represent mean ± SEM, **j-m**, n=17 cells from 3 sg*Rosa26* mice, and n=33 from 4 sg*Tacr3* mice. *Bars represent mean ± SEM, dots within bars represent mean behavior duration for each animal (diamonds=females, circles=males). ns p > 0.05, * p < 0.05, ** p < 0.01, *** p < 0.001, **** p<0.0001*.

### Prefrontal Neurokinin-3 Receptors amplify NkB’s influence over behavior

To understand the contribution of mPFC Nk3R’s to the internal state induced by social-isolation, we employed a CRISPR-mediated approach to generate nonsense mutations in *Tacr3*. To achieve this, an AAV encoding a Cre-dependent Staphylococcus aureus Cas9 (SaCas9) and guide RNA (sgRNA) directed to *Tacr3* were injected into the mPFC of Tacr3-Cre^78^ mice alongside a Cre-dependent KASH-eGFP virus^73^ to induce nonsense mutations in *Tacr3* in the mPFC (Fig. 5e) and visualize the infusion site (Extended Data Fig. 11b). Control mice were injected with an AAV encoding a Cre-dependent SaCas9 and sgRNA directed to Rosa26^72^ alongside the Cre-dependent KASH-eGFP virus as above. Mice were then isolated and tested on the Resident Intruder assay (Fig. 5e). We found that *Tacr3* mutagenesis was sufficient to reduce investigatory behaviors but had no effect on aggression (Fig. 5f-i), consistent with the results we obtained for *Tac2* manipulations (Fig. 4n). Thus, our loss-of-function manipulation targeting mPFC Nk3Rs phenocopied our shRNAi-mediated knockdown of *Tac2* suggesting that Nk3Rs serve to amplify the influence of NkB on behavior.

To verify that CRISPR/SaCas9 mutagenesis of *Tacr3* sufficiently reduced Nk3R-evoked depolarization, Tacr3-Cre mice were infused with an AAV encoding a Cre-dependent GCaMP as well as the SaCas9-sgTacr3 or SaCas9-sgRosa26 (control) AAVs. Senktide, the Nk3R agonist, was applied and calcium signals (dF/F) were recorded (Fig 5j-m). SaCas9-sgTac3 significantly reduced the number of cells responsive to senktide application (Fig. 5l) and significantly reduced the change in fluorescence of GCaMP in response to senktide application (Fig. 5m), confirming the efficacy of our SaCas9-sgTacr3 approach.

Importantly, we found that neither chemogenetic inhibition nor excitation of Tacr3^+^ cells at test was sufficient to produce any effects on isolation-induced behavior (Extended Data Fig. 12), further supporting the idea that NkB signaling across the window of isolation is critical for the effects of isolation to induce changes in behavior.

As CRISPR-mediated *Tacr3* disruption across the window of isolation was sufficient to attenuate isolation-induced enhancements in investigation (Fig. 5e-i), we next tested whether chronic activation of Tacr3^+^ cells across a “fictive” window of isolation could generate an isolation-like phenotype in group housed mice (Extended Data Fig. 13). Group housed, Tacr3-Cre mice were injected with a Cre-dependent virus encoding the excitatory DREADD hM3Dq into the mPFC (Extended Data Fig. 13a). Mice were administered CNO-spiked or standard (control) drinking water in their homecage for three weeks to induce chronic activation of Tacr3^+^ neurons across this time frame^9^. They were then tested on the Resident Intruder assay following an additional injection of CNO or saline (controls)^9^. This chronic gain-of-function manipulation was not sufficient to induce any changes in behavior for group housed mice (Extended Data Fig. 13b-e), suggesting that while Nk3R signaling is necessary for the effects of isolation to enhance investigatory behaviors, activation of the cells which express *Tacr3* is not sufficient to “convert” group housed mice into isolated mice, as multiple gain-of-function manipulations may be required, consistent with prior findings from our group and others^3, 9, 34, 36^.

Our *Tacr3* CRISPR findings suggest that because mutagenesis of *Tacr3* phenocopies the effects of disrupting Tac2/NkB signaling, Nk3R signaling mediates the increased investigation observed in isolated mice. Collectively, our data highlight the specific contributions of Tac2/NkB, GABA, and Nk3Rs to dissociable behaviors that span an aggressive encounter in isolated mice.

## DISCUSSION

These findings demonstrate that co-transmission of a stimulatory neuropeptide and GABA enable the coordinated control of sequential, mutually exclusive behaviors that tile an aggressive encounter. This work contributes a new mechanism by which a single neuronal population controls distinct but complementary behaviors through the release of both slow-acting, long-range neuropeptides and fast-acting, classical neurotransmitters. Importantly, our results suggest that the cumulative impact of neuropeptide release across a long period of time (e.g., the window of isolation) is sufficient to put an animal in a heightened state of arousal, characterized by an increase in investigatory behaviors. In contrast, the fast-acting and localized effect of GABA release is well-suited for the tight, time-locked control over aggression. Collectively, these results are the first to demonstrate that a single population of neurons controls mutually exclusive behaviors via the distinct signaling properties of the molecules it releases. In contrast, the handful of prior studies examining the functional impact of neuromodulator-neurotransmitter co-transmission either focus on signaling molecules with the same “sign” of action^79, 80^, opposing signaling molecules that converge to impact the same behavior^4, 8^, or opposing signaling molecules that control opposing states^7, 81^.

Prior work from our group and others has established that prolonged social isolation induces aggression^9, 42, 43, 82^. Here, we find that it also increases a number of social behaviors, including face interaction. Face-to-face contact, sometimes referred to as a face “threat”^83^ or “hard look”^84^, has been implicated in threatening encounters. Interestingly, we found that mPFC Tac2^+^ neuron activity showed no tuning to face interaction, nor did any of our *Tac2* loss-of-function perturbations impact this behavior. This suggests that face interaction may be controlled by a distinct signaling system operating in parallel to Tac2^+^ neurons that is similarly sensitive to isolation. Indeed, we found that inhibiting mPFC Crh^+^ neurons attenuated isolation-induced face interaction, indicating that multiple cell types and signaling systems work collectively to exert top-down control over behavior^57^.

Classically, the idea that neuromodulators play a specialized role in the control of internal states has largely focused on subcortical regions^1, 8, 85, 86^. In contrast, the role of neuropeptidergic signaling in the cortex is relatively unknown^12^. Recent work has revealed a role for neuromodulatory signaling in the mPFC in the control of fear^87^, suggesting that neuromodulatory action within the cortex may be critical for the top-down control over behavior. Here, we reveal that mPFC Tac2^+^ neurons, NkB signaling, and GABA release from these cells is critical for exerting control over distinct phases of an aggressive encounter in isolated animals. To our knowledge, these data represent the first discovery of a specific function for both Tac2^+^ and Tacr3^+^ cells in the mPFC. These results are consistent with a broader role for the mPFC in selecting behavioral responses based on internal and external information from converging inputs,^13^ and its control over behavioral transitions^87, 88^.

Our findings build upon prior work identifying a role for mPFC neurons in the control aggression^24, 26, 42, 57, 89–91^ and are consistent with a number of potential circuit motifs to explain the role of Tac2^+^ neurons in the top-down control of SI-aggression. Given that we identified mPFC Tac2^+^ neurons to be GABAergic, one parsimonious motif to explain our results is a “inhibition” model, where Tac2^+^ neurons would control aggression via their ability to locally inhibit pyramidal neurons (PNs), which as a whole are thought to exert top-down control (or apply the “brakes”) over aggression^42, 57, 92, 93^. This is consistent with our cFos and fiber photometry studies, where we found that isolation increased Tac2^+^ neuron activity, which, in turn, would inhibit PNs, thereby “releasing the brakes” on aggression. Moreover, given the expansive arbor of mPFC Tac2^+^ neurons, which reach into L5 (Fig 3 l-n), it is possible that Tac2^+^ neurons can directly inhibit L5 pyramidal neurons. Given that we found that roughly 50% of Tac2^+^ neurons are Vip^+^ (Tac2^+^VIP^+^ neurons), a second possible circuit motif to explain our results would be a “disinhibition” model. This is supported by the well characterized role of early layer cortical Vip^+^ neurons in the disinhibition of pyramidal neurons via their inhibition of intermediary Som^+^ interneurons that tonically inhibit PNs^94–97^. In this model, Tac2^+^Vip^+^ cells would work to disinhibit the specific class of PNs that have been shown to be important for driving aggression via their direct projections to hypothalamic structures^56^. This second disinhibition motif could be complemented by the first inhibition motif, which would operate in Tac2^+^ VIP^-^neurons. Indeed, given that we found no evidence for the preferential activation of any Tac2^+^ neuron subclass by isolation-induced aggression (Extended Data Fig. 3b-e), it may be more likely that a combination of motifs is at play. Lastly, in a third potential circuit motif, deeper Tac2^+^Vip^+^ neurons could directly inhibit the larger class of PNs thought to inhibit aggression ^97, 98^. Indeed, while Vip^+^ neurons have been heavily implicated in the disinhibition of PNs, recent work suggests that *Vip* labels a heterogenous population, with some subclasses of Vip^+^ cells working to disinhibit PNs and others operating to inhibit them^99–101^. This third model would be consistent with a “pull-push” motif, where changes in arousal state enables Vip^+^ cells in the frontal cortex to both directly inhibit PNs and excite them via disinhibition^102^.

In addition, our work suggests that cumulative neuropeptide signaling across an extended period may be a key mechanism for the control of internal states that build up across time, such as social isolation. Indeed, we found that while caspase-mediated ablation of Tac2^+^ neurons reduced the impact of isolation on both investigation and aggression, our DREADD-mediated inhibition of Tac2^+^ neurons reduced aggression but did not impact investigatory behaviors. The discrepancy in these results could be explained in two ways. The first is that hM4Di DREADDs not only hyperpolarize neurons via G-protein-gated inwardly rectifying potassium (GIRK) channels, but are also able to inhibit neurotransmitter release through the bg subunit’s interaction with the SNARE protein, SNAP-25^75, 76^, potentially resulting in the disruption of GABAergic signaling while maintaining neuropeptide release from this cell population. While there is evidence that SNAP-25 is part of neuropeptidergic dense core vesicle fusion and release^103, 104^, it remains understudied, making it difficult to assess whether hM4Di-mediated inhibition would impact neuropeptide release. The second is that caspase-mediated ablation resulted in the loss of mPFC Tac2^+^ neurons across the entire window of isolation as well as at test. In contrast, hM4Di inhibition only rendered Tac2^+^ neurons acutely silent at test. This suggests that one critical mechanism by which Tac2^+^ neurons exert control over the state of social isolation is by chronic release of NkB. Indeed, we found that NkB expression in the mPFC increases across the window of isolation, supporting the idea that prolonged social isolation may induce the repeated release of NkB from Tac2^+^ cells^3, 9^, which, in turn, may be critical for mediating the effects of isolation on behavior.

Importantly, our work expands our understanding of how prolonged social isolation impacts behavior across males and females alike. While both acute and prolonged isolation have been shown to have sex-dependent effects on social behavior^42, 47, 48, 55^, the work presented here demonstrates that both sexes respond to prolonged periods of social isolation with an increase in same-sex aggression. Interestingly, we found that in naïve animals, females tended to show an overall increase in isolation-induced investigatory behaviors in contrast to males. This suggests potentially distinct strategies in how males and females approach an aggressive encounter and is consistent with prior work highlighting sex-differences in aggression^45, 105–107^. Nevertheless, key effects were found across both sexes following isolation including the rapid shutdown of investigatory behaviors following the initiation of aggression (Fig. 1k-o), the increase in aggressive behavior (Fig. 1f), and the attenuation of investigation and aggression following Tac2^+^ neuron related loss-of-function perturbations (Fig. 4). While our results here and previously^9^ suggest a similar function for Tac2/NkB signaling in both sexes, recent work has identified sex-differences in Tac2-mediated control of fear behavior^108–110^. This suggests that either Tac2/NkB exerts dissociable, sex-specific effects in cortical vs. subcortical structures, or that Tac2/NkB may function to control distinct states (e.g., isolation and fear) in a separable fashion. Evidence that sex can influence neuropeptide expression or that neuropeptide receptor knockout can induce sex-dependent changes in behavior have been demonstrated in other systems^111, 112^, highlighting the need for future work to interrogate the role of Tac2/NkB across cortical and subcortical regions in a variety of behavior states and across sexes.

Aggression is a social behavior critical for survival. However, when elicited in excess or as a result of stress, aggression can become pathological^91^. As with other forms of pathological behavior, the mPFC is thought to offer promise as a region capable of exerting top-down control over aberrant social behavior^57, 91, 113, 114^. Here, we identify the neuropeptide NkB as a critical player in the circuitry of isolation-induced aggression, not just subcortically, but in the cortex as well. We find that mPFC Tac2^+^ neurons enable a delicate balance between the behaviors displayed during an aggressive encounter in response to social isolation. These findings suggest both a target region and a target cell type for the control of isolation-induced aggression and reveal a novel neurochemical mechanism by which a single population of neurons is able to exert coordinated control over distinct behaviors that comprise an internal brain state via the release of opposing signaling molecules.

## MATERIALS AND METHODS

### Animals

Wild-type (WT) C57BL/6N male and female mice (Charles River, strain: 027) and BALB/C intruders (strain: 028) were obtained from Charles River (7-10 weeks of age). Male and female heterozygous Tac2-IRES-Cre mice^115^ (Jackson Labs, Strain #:021878), “Tac2-Cre”, Crh-IRES-Cre (Jackson Labs, strain #: 012704, “Crh-Cre”) and Tacr3^iCre:GFP^ (“Tacr3-Cre”, Jackson labs, Strain #:035576) were bred and maintained in house at the University of Utah animal facility. Cre mice were backcrossed to wildtype C57BL/6N’s to reduce genetic drift. Mice were housed and maintained on a reverse 12-hr light-dark cycle with food and *water ad libitum*. All behavior testing was performed during the dark cycle. Experimental manipulations and animal care were performed following the National Institutes of Health Guide for Care and Use of Laboratory Animals and approved by the University of Utah Institutional Animal Care and Use Committee. Slice electrophysiological experiments were performed at the University of Washington in accordance with the University of Washington Institutional Animal Care and Use Committee.

### Resident Intruder assay

8-10-week-old wildtype C57Bl6/N, Tac2-IRES-Cre, Crh-Cre or Tacr3-Cre mice were single housed or remained group housed for a minimum of 2 weeks. For the Resident Intruder (RI) assay, mice were transferred to the behavioral testing room and allowed to acclimate in the dark for approximately 30 minutes prior to the start of testing. Mice remained in their home cage for the duration of testing. Homecages were placed into a clear Plexiglas chamber to enable video recording of the “Resident” mouse via an infrared Basler camera mounted overhead.^9^ Behavior was recorded for three minutes (baseline), then an age- and sex-matched Balb/c “Intruder” mouse was placed into the resident’s homecage for 10 minutes (test phase). For group housed animals, only one mouse was tested in the homecage at a time, while the other cage mates were placed in a clean holding cage until testing. Once tested, mice were placed in a separate clean holding cage to avoid behavioral contamination or communication between tested and untested mice. Following the completion of testing for all mice in each cage, the mice were returned to their home cage. The apparatus was cleaned with 70% ethanol between all tests to reduce olfactory cues.

#### Behavior scoring

All behavior (aggressive behaviors, face interaction, body investigation, anogenital investigation) was manually scored by a trained, blinded observer using Noldus Observer. Data was then exported to Excel (Microsoft) and further analyzed using Prism (GraphPad), Matlab or Python. *Face Interaction* was classified as frames in which the resident sniffed the facial area of the intruder. *Body Investigation* was classified as frames in which the resident sniffed the torso of the intruder. *Anogenital Investigation* was classified as frames in which the resident sniffed the anogenital region of the intruder. *Aggression* was classified as frames in which the resident engaged in biting the front, flank, or back of the intruder, biting or gnawing the back or neck of the intruder from a dominance mounted position, tussling, and barbing or biting the face of the intruder.

#### Transition matrices

Behavioral transition matrices were generated by calculating the probability of a mouse transitioning from one of the four scored behaviors (face interaction, body investigation, anogenital investigation, aggressive behavior) to the next behavior within a one-second window of time^53^. Code used to generate transition matrices can be found on Zelikowsky Lab GitHub page (see data availability).

#### Temporal analyses of behavior transitions

Socially isolated mice with aggressive bouts were used in these analyses. To characterize the temporal distribution of investigatory social behaviors before and after aggression, pre-aggression was defined as the period prior to the onset of the first aggressive bout, and post-aggression as the period after the onset of the first aggressive bout. All analyses were performed in MATLAB.

### RNAscope (fluorescent in situ hybridization)

For cFos RNAscope™ experiments, mice were anesthetized using isoflurane (4-5%), rapidly decapitated, and brains extracted 30 minutes from the start of the test phase of the RI assay. Brains were rapidly flash frozen in 2-methylbutane (Sigma-Aldrich, catalog #: M32631-4L) and stored at -80 °C for up to 4 months. Cryosections were taken at 20µm thick and mounted directly onto slides (SuperFrost PLUS Adhesion slide, Electron Microscopy Sciences, catalog #: 71869-11) in the cryostat. Roughly 4-5 sections from two animals, one per housing condition, were placed on each slide to control for slide-to-slide variability. Slides were stored in airtight containers at -80°C for up to four months before the RNAscope™ assay was performed. The RNAscope™ assay was performed according to the manufacturer’s instructions ^63^ (Advanced Cell Diagnostics). The following probes (ACDBio) were used: *Tac2*, *PV, Som*, *cFos*, *Vip*, *Crh, Slc17a7, Slc32a1*. For experiments using *Gad1* and *Gad2* probes, a 50:50 mixture of each probe was used and is indicated by “*Gad1/2*”. Slides were then imaged on a Keyence BZ-X800 microscope and/or an Echo SDC-G spinning disk confocal within one week of the assay.

### cFos Immunohistochemistry

For cFos immunohistochemistry experiments, mice underwent transcardial perfusions with 0.9% saline (MWI Animal Health, catalog # 033500) followed by 4% paraformaldehyde (PFA; Paraformaldehyde powder, CAS 30525-89-4, Sigma Aldrich, catalog #: 158127-500G) 90 minutes after the middle of the test phase of the Resident Intruder assay and post-fixed in 4% (PFA) overnight at 4°C. Brains were then transferred to 15% sucrose solution (Sigma-Aldrich, S0389-5KG, CAS 57-50-1) at 4°C overnight. The brains were then transferred to 30% sucrose solution overnight at 4°C. Brains were then sectioned at 40 μm into 1X PBS (Gibco, ref 10010-023) and stored at 4°C for up to a week before immunohistochemistry was performed. Sections were incubated in 10% Normal Donkey Serum (Sigma-Aldrich, catalog #: D9663-10mL) + 0.3% Triton X-100 (Sigma, X-100) in 1X PBS (Gibco, ref 10010-023) for one hour at room temperature. The sections were incubated in the primary antibody (rabbit anti-cFos, Abcam 190289) at 4°C for 72 hours. Following this incubation, sections were rinsed in 1X PBS (Gibco, ref 10010-023) three times, then incubated in secondary antibody (Donkey anti-rabbit Alexa Fluor-488, 1:250 dilution in blocking buffer, catalog #: A-21206, ThermoFisher) for 12 hours. After this period, sections were rinsed three times in 1X PBS (Gibco, ref 10010-023), then mounted onto slides (SuperFrost PLUS Adhesion slide, Electron Microscopy Sciences, catalog #: 71869-11) and counterstained with DAPI mounting media (DAPI Fluoromount-G Mounting Medium, Liquid, Southern Biotech, catalog # 0100-20). Slides were imaged with a Keyence BX-Z800 or Echo spinning disk confocal by a blinded experimenter within seven days of mounting. The total number of mCherry-positive cells and the number of mCherry-positive cells co-labelled with cFos were quantified on sections with viral expression in the mPFC (FIJI). GraphPad PRISM was used to perform nested t-tests for the number of virus-positive, cFos^+^ cells out of the total number of mCherry-positive cells.

### Neurokinin B Immunohistochemistry

After 1 day or 3 weeks of social isolation or group housing conditions, C57Bl6/N mice (8-10 weeks at start of experiment) were transcardially perfused with 1X PBS (Gibco, Cat. No. 10010-031), followed by 4% paraformaldehyde (PFA; Sigma-Aldrich, Cat. No. 158127). Brains were post-fixed overnight in 4% PFA at 4°C, transferred to 15% sucrose for 24 h followed by 30% sucrose for an additional 24 hours for cryoprotection. They were then embedded in Tissue-Tek O.C.T. Compound (Sakura Finetek, Cat. No. 4583), frozen on dry ice, and stored at -80°C until sectioning. Coronal brain sections (30μm) were cut using a cryostat (Leica Biosystems), collected in PBS, mounted onto Superfrost Plus slides (Fisher Scientific, Cat. No. 12-550-15) and stored at -20°C until immunostaining. Sections were washed three times for 5 min each in 1X PBST (1X PBS containing 0.1% Tween 20) and blocked in blocking buffer (10% normal goat serum + 0.3% Triton X-100 in 1X PBS) for 1 h at room temperature and incubated overnight at 4°C with rabbit anti-Neurokinin B primary antibody (1:200; Abcam, Cat. No. AB203050). The following day, sections were washed three times in PBST and incubated for 2 h at room temperature with Alexa Fluor 594 goat anti-rabbit IgG secondary antibody (1:300; Invitrogen, Cat. No. A11012). Sections were then washed three times for 10 min each in PBST and cover slipped using DAPI Fluoromount-G mounting medium (SouthernBiotech, Cat. No. 0100-20). Images were scanned using a Zeiss Axio Scan 7 slide scanner. Regions of interest were defined and exported in ZEN 3.4 and mean fluorescence intensity was quantified using ImageJ.

### Single-nucleus RNA sequencing

The mPFC (prelimbic and infralimbic cortices) was dissected from group housed and isolated C57Bl/6N male mice in ice-cold RNase-free 1X PBS containing 15μM actinomycin-D (Sigma-Aldrich, catalog #A1410-10MG). Nuclei were isolated using manual homogenization and the Nuclei PURE Prep kit (Millipore Sigma, catalog # NUC201-1KT) with 15 µM actinomycin-D added to the lysis buffer. Nuclei were resuspended in a resuspension buffer containing 0.5% Ultrapure BSA (Fisher Scientific #AM2616) to prevent clumping of nuclei and 0.2 U/µl RNase inhibitor (New England Biolabs #M0314S) to prevent RNA degradation^116–118^. Single-nuclei barcoded cDNA libraries were generated by the University of Utah High Throughput Sequencing using the 10x Genomics user guide “Chromium Next GEM Single Cell 3’ Reagent Kits v3.1”. Libraries were sequenced on the Illumina NovaSeq 6000. Fastq files were aligned to the GRCm39 reference genome using 10X Cell Ranger (v4.0.8). Filtered gene expression matrices were then imported into R (v4.3.0), and nuclei with fewer than 800 features, greater than 6000 features, or greater than 5% mitochondrial reads were removed to eliminate low-quality nuclei and multiplets. Seurat (v4.2.1)^119^ was used to calculate nearest neighbors at 0.8 resolution from the first 30 principal components and cluster nuclei using UMAP dimensionality reduction. Then, clusters were assigned to known cell types based on expression of known cell-type-specific marker genes^77, 120^. UMAP plots describing expression of genes of interest were generated using the Seurat FeaturePlot function. Code used in analysis of single nucleus RNAseq can be found on the Zelikowsky Lab GitHub page (see data availability) and in our prior work^118^.

#### Tac2 and Tacr3 Proportion Analysis

To determine the proportion of Tac2^+^ or Tacr3^+^ cells that belong to each cell-type cluster a custom R script was used to find the number of cells expressing the gene of interest in each cluster for each sample. The proportion of total cells expressing the gene of interest belonging to each cell-type was then calculated for every sample, exported to a CSV file, and plotted in GraphPad Prism. Similarly, the proportion of cells with the gene of interest in broader categories, such as inhibitory clusters, was calculated by summing the number of cells expressing the gene of interest in clusters of that type and dividing by the total number of cells expressing the gene of interest. Further analyses were performed to examine the proportion of Tac2^+^ cells that belong to each cell-type cluster for cells that had low expression levels of *Tac2* (“low expressing”; <1 copy) or high levels of Tac2 (“high expressing”; >2 copies). The dataset was then filtered by these parameters and the same analysis described above was then performed to calculate proportions within the subsets of cells.

### Surgical procedures

Mice were anesthetized in an induction chamber using 4-5% isoflurane, and anesthesia was maintained at 1-2% for the duration of the procedure while mice were mounted on a stereotaxic surgery rig (Kopf Instruments, California, USA). Once mounted, ophthalmic ointment (Soothe lubricant eye ointment, Bausch and Lomb) was applied to prevent the eyes from drying. Topical hair removal cream (Nair) was applied to the scalp to remove hair. Following removal of cream and wiping with distilled water, the surgical area was cleaned 3x with povidone-iodine solution (Dynarex, catalog 1415) and isopropyl alcohol (Solimo). A subcutaneous injection of sustained-release buprenorphine at a dose of 3.25 mg/kg animal weight (Ethiqa XR 1.3mg/mL, Fidelis Animal Health) was given before incisions to provide long-lasting analgesia. A small incision was made along the scalp to expose the skull, the skin was retracted, and the skull was levelled in all directions using a Model 1905 Stereotaxic Alignment Indicator (KOPF). Burr holes were drilled over mPFC coordinates for subsequent viral infusion.

All infusion coordinates into the mPFC were made in relation to bregma (mm). For all experiments, bilateral infusions were performed at AP: +2.14, ML: +/- 0.25, DV: -2.79. Virus was backfilled into pulled fine glass capillaries (∼ 50 µm diameter tip; World Precision Instruments, item no. 504949) filled with mineral oil (sigma Aldrich, M5904, CAS 8042-47-5) and 140nL virus/side was infused at a rate of 80nL/min (Nanoliter 2000, World Precision Instruments) controlled by an ultra-micro-syringe pump (Micro4, World Precision Instruments) unless otherwise specified. Capillaries remained in place for 5 minutes following infusion to allow for full diffusion of virus and to reduce backflow up the infusion tract. For most experiments the scalp was drawn together and sealed with GLUture (World Precision Instruments, catalog # 503763). For fiber photometry experiments isolated male and female Tac2-IRES-Cre mice were infused with 200µL of a Cre-dependent GCaMP6f (AAV1-Syn-Flex-GCaMP6f-WPRE-SV40; Addgene #10083; coordinates: AP +2.13, ML: +0.3, DV-2.78) into the mPFC and a fiber optic cannula (Doric, MFC_400/430-0.66_3.0mm_ZF2.5_FL) was implanted along the viral tract at 0.1mm above the DV coordinate of the viral infusion site. After implantation, the skull was covered in Metabond (Parkell C&B-Metabond Quick Luting Cement base and Radiopaque L-Powder S398) and PEMA temporary resin acrylic (Bosworth Trim II) to secure the implant. Following completion of an experiment, mice were perfused and brains were extracted, sectioned, mounted, stained with DAPI and coverslipped for histological verification. Infusion sites were then confirmed using epifluorescence microscopy (Leica DM2000 LED).

### Histology

For all experiments using AAV vectors (Dendritic arbor characterization, DREADDs, shRNA, CRISPR, caspase), brains were extracted from animals within one week of the completion of experiments. Animals were anesthetized with isoflurane (4-5%) and rapidly decapitated, and brains were then extracted and post-fixed in 4% paraformaldehyde (CAS 30525-89-4, Sigma Aldrich, catalog #: 158127-500G) overnight at 4°C. Brains were then transferred to 15% sucrose overnight at 4°C, followed by 30% sucrose overnight at 4°C. After the final sucrose transfer, brains were embedded in OCT (Tissue-Tek O.C.T Compound, Sakura Finetek USA Inc., catalog #: 4583) and stored at -80 °C until cryo-sectioning. Brains were then sectioned, mounted, stained with DAPI and cover slipped for histological verification. Viral expression and/or fiber placement was verified for all animals using epifluorescence microscopy (Leica DM2000 LED).

### Characterization of the axonal arbor of mPFC Tac2^+^ neurons

Male and female Tac2-IRES-Cre mice were single-housed (3 weeks) and underwent surgery in which a Cre-dependent adeno-associated virus (AAV) expressing membrane-GFP and synaptophysin fused to mRuby (AAV2-hSyn-FLEx-mGFP-2A-Synaptophysin-mRuby; Addgene #: 71760-AAV2, titer: 2.7 x 10^12^ GC/mL, lot: v138755) was infused into the mPFC. Following single housing and surgical recovery, histology and imaging was performed as described above.

### Fiber photometry

#### Behavior

Tac2-Cre mice were infused with a Cre-dependent GCaMP and implanted with a ferule fiber targeting the mPFC (see surgical details above). They were subsequently isolated and allowed to recover for 5 weeks. Two days before behavior, mice were habituated to the fiber photometry patchcord (Doric Lenses, MFP_200/230/900-0.57_3m_FC-ZF2.5). Optic fibers were attached to the cable, and mice were placed in the Resident Intruder chamber for 10 min. On test day, behavior was recorded while the resident freely roamed their home cage for 5 minutes (baseline), then an age and sex-matched Balb/c intruder was introduced, and behavior was recorded for 15 minutes (as described above). Behavior was manually scored by a blinded experimenter using Noldus Observer (as described above).

#### Data acquisition

Fiber photometry signals were acquired using a FP3002 (Neurophotometrics, MPF Bioscience) photometry system at a sampling rate of 60Hz, with an LED excitation power at the fiber tip of 50µW. Power was tested prior to the start of behavior testing using a Thor labs laser meter (Thor labs, PM160 Si sensor power meter) for both 415nm and 470nm channels. Acquisition was controlled via Bonsai software, which simultaneously triggered video recording in Noldus Ethovision at the onset of photometry data acquisition. Photometry and behavioral video streams were synchronized via TTL pulse. Two signals were collected, a 415nm isosbestic, Ca²⁺-independent signal, and a 470nm Ca²⁺-dependent signal.

#### Fiber photometry data analysis

For each session, raw fluorescent traces were normalized by dividing the 470nm signal by the frame-matched 415nm signal. The resulting normalized trace was z-scored relative to a moving baseline window (3s duration, -5 to -2s before each behavior onset): for each behavioral event dF/F in the 5s before and after onset was z-scored by subtracting the mean baseline dF/F and dividing by its standard deviation. This baseline window was applied uniformly to all behavior onsets. To ensure independence of events, only behavior bouts separated by at least 5s from the preceding bout of the same category were included in analyses. For investigatory behaviors, statistical differences were assessed by comparing mean dF/F during the baseline window (-4s to -2s) to the onset window (1-3s). For aggression, statistical significance was assessed using a circular permutation test (1000 iterations). N=7 mice, 409 total behavior bouts (face: 108 bouts, body: 114 bouts, anogenital: 96 bouts, aggression: 91 bouts).

Sessions were excluded if there were indications of a poor fiber connection, as indicated by a normalized Ca²⁺ signal that failed to exceed 2 standard deviations above the baseline mean (baseline window: 30 s, −35 s to −5 s before intruder introduction) for at least 0.5 s (30 consecutive frames) following intruder introduction. 0.5 s Gaussian smoothing was applied only for visualization in figures and was not applied to the underlying data used for statistical analysis. Code used in fiber photometry analyses can be found on the Zelikowsky Lab GitHub page (see data availability).

### DREADD-mediated manipulation of neural activity

#### Acute DREADDs

For acute DREADDs manipulations (DCZ delivery at test only), Tac2-IRES-Cre, Crh-Cre or Tacr3-Cre mice were screened for aggression by single housing for two weeks before test to a baseline level of aggressiveness so it could be determined whether silencing/activating neurons had an impact on aggression as these experiments were within subjects. Animals that exhibited aggression were then infused with a virally-mediated Cre-dependent DREADDs (AAV2-hSyn-DIO-hM4Di-mCherry, Addgene # 44362, titer 3.3×10^13^ GC/mL, lot v138755, 140nL per side) bilaterally into the mPFC. Following surgeries, mice remained single-housed for 3 weeks before being tested on the Resident Intruder assay twice (tests separated by 48 hours). Prior to behavioral testing, mice received 0.1 mg/kg body weight Deschloroclozapine (DCZ, CAS: 9177-07-7, MedChemExpress, catalog #: HY-42110) or vehicle (1.5% DMSO/saline) via intraperitoneal injection 10 minutes before the start of the baseline phase of the RI assay. A within-subjects design was used, such that each animal received DCZ on one test day and vehicle on the other (test order counterbalanced). Mice were randomly assigned to receive either DCZ or vehicle first, with roughly equal numbers/group/day. Subsequent behavioral testing and analyses proceeded as described above. DREADDs efficacy was confirmed by assessing cFos immunoreactivity as an indirect marker of cell activity (see cFos immunohistochemistry).

To determine whether DCZ had off-target effects on mouse behavior in the resident intruder assay, male and female WT littermates of Tac2-IRES-Cre mice were injected with either 0.1mg/kg, 0.25mg/kg, 0.5 mg/kg or 1.0 mg/kg body weight of DCZ on one test day and vehicle on the other (order counterbalanced). The behavioral testing on RI occurred as described for DREADDs experiments. One week later, mice were tested on a different dose of DCZ to control for variability across mice.

#### Chronic DREADDs

To chronically activate mPFC *Tacr3* neurons, group-housed male and female Tacr3-Cre mice were infused with AAV2-hSyn-DIO-hM3Dq-mCherry (Addgene # 44361, titer 2.4×10^13^ GC/mL, lot v1193467, 140nL per side). Clozapine N oxide (CNO, Enzo LifeSciences, catalog #: BML-NLS105_00056, lot #: 44HV22) was administered at 0.5mg per 100 mL water to mice through their drinking water for three weeks to constitutively activate (hM3D) mPFC Tacr3 cells^9^. Control animals received standard water. Each cage was provided with enough water such that each mouse in the cage could consume 10mL of water per day. On test day, a 2mg/kg dose of CNO (for CNO water group) or vehicle (for standard water control group) was administered via intraperitoneal injection 20 minutes before the RI test began.

### Tac2 shRNA experiments

Wild-type (WT) C57BL/6N male and female mice received infusions of a Tac2 shRNA (AAV2-CMV-shTac2-zsGreen, titre: 5.03×10^12^ GC/mL) virus or a luciferase control virus (AAV2-CMV-shLuciferase, titre: 2.37×10^12^ GC/mL) into the mPFC. After 4 weeks of expression and social isolation, animals were tested on the Resident Intruder assay.

To validate the efficacy of shRNA-mediated *Tac2* knockdown, mPFC sections from all experimental and control mice were processed for *Tac2* mRNA expression. Following the completion of behavioral testing, brains were flash frozen within three days and sectioned in preparation for RNAscope™ (see methods above). For RNAscope™, each slide contained two mice, one from each viral condition, and the *Tac2* probe was used to visualize *Tac2*. Experimenters were blinded to both the sex and viral condition of animals during RNAscope™, imaging, behavior scoring, and cell counting. Knockdown of *Tac2* was verified by taking automated counts of DAPI and manual counts *of Tac2* in FIJI (quantified by expression in *Tac2* channel that co-localized with DAPI signal). GraphPad PRISM was used for a nested t-test statistical analysis of cell count data. Viral placement and expression were verified for all animals as described in histology above.

### Tac2 caspase experiments

Tac2-IRES-Cre mice received bilateral infusions of a viral cocktail containing either 1 part AAV1-hSyn-mCherry (Addgene 114472-AAV1, lot V177823, titre: 1.9 x 10^13^ GC/mL): 5 parts AAV5-Ef1a-FLEX-taCasp3-TEVp (Addgene 45580-AAV5, lot V219498, titre: 2 x 10^13^ GC/mL) or 1 part AAV1-hSyn-mCherry: 5 parts AAV5-Ef1a-DIO-eYFP (Addgene 27056-AAV5, lot V213843, titre 1.9 x 10^13^ GC/mL). A total of 140nL was injected. Following surgery, mice were single housed for four weeks to allow for viral expression, after this isolation period, animals were tested on the Resident intruder assay as described above. Behavior was scored and analyzed as described above.

To validate the efficacy of caspase-mediated ablation of mPFC Tac2^+^ neurons, brains were extracted and flash frozen within four days and sectioned in preparation for RNAscope™ as described above. Following the RNAscope™ assay for *Tac2* and imaging as described above, the total fluorescence intensity (mean gray value) of *Tac2* was quantified by a blinded experimenter using FIJI and a nested t-test was used to compare between groups.

### CRISPR mutagenesis experiments

#### sgSlc32a1 (vGAT) mutagenesis in mPFC Tac2 neurons

Tac2-IRES-Cre mice received bilateral infusions of a cocktail containing either 1 part EGFP-KASH: 7 parts saCas9 (AAV1-CMV-FLEX-saCas9-U6-sgSlc32a1) or 1 part AAV1-EGFP-KASH: 7 parts AAV1-CMV-FLEX-saCas9-U6-Rosa26. Following surgery, mice were single-housed for four weeks to allow for viral expression. After this period, mice were tested on the Resident Intruder assay as described above. Behavior was scored and analyzed as described above.

To confirm CRISPR-mediated nonsense decay of RNA transcripts for *Slc32a1*, brains from sgSlc32a1 and sgRosa26 groups were flash frozen and prepared for RNAscope as described above. Before performing RNAscope, virus expression was confirmed by epifluorescent microscopy on fixed slides. In animals, that had virus expression, probes for *Slc32a1* and *Tac2* were used in the RNAscope assay. Sections were then imaged, and the fluorescence intensity (mean grey value) of the *Slc32a1* signal within a mask (outline of the *Tac2* signal) was quantified by a blinded experimenter using FIJI.

#### Slice Electrophysiology

Tac2-IRES-Cre mice were infused with AAV1-CMV-FLEX-saCas9-U6-sgSlc32a1 or AAV1-CMV-FLEX-saCas9-U6-Rosa26 and AAV1-Ef1a-FLEX-hChR2(H134R)-mCherry (1×10^13 GC/ml, lot v62777, Addgene 20297-AAV1) into the mPFC. Mice were euthanized and brains were rapidly extracted and coronal brain slices (200 μm) were prepared in an ice-cold slush solution containing (in mM): 92 N-methyl-D-glucamine, 2.5 KCl, 1.25 NaH₂PO₄, 30 NaHCO₃, 20 HEPES, 25 glucose, 2 thiourea, 5 sodium ascorbate, 3 sodium pyruvate, 0.5 CaCl₂, and 10 MgSO₄ (pH 7.3–7.4). Slices were allowed to recover for ≤12 min in the same solution at 32 °C and were then transferred to a room-temperature solution containing (in mM): 92 NaCl, 2.5 KCl, 1.25 NaH₂PO₄, 30 NaHCO₃, 20 HEPES, 25 glucose, 2 thiourea, 5 sodium ascorbate, 3 sodium pyruvate, 2 CaCl₂, and 2 MgSO₄. Slices were incubated for an additional 45 min before recordings were performed in artificial cerebrospinal fluid (ACSF) at 32 °C, continuously perfused at ∼2 mL min⁻¹. The ACSF contained (in mM): 126 NaCl, 2.5 KCl, 1.2 NaH₂PO₄, 1.2 MgCl₂, 11 D-glucose, 18 NaHCO₃, and 2.4 CaCl₂. All solutions were continuously bubbled with 95% O₂ and 5% CO₂. Whole-cell recordings were performed using an Axopatch 700B amplifier (Molecular Devices), with signals filtered at 1 kHz. Recording electrodes (3–5 MΩ) were filled with an internal solution containing (in mM): 130 potassium gluconate, 10 HEPES, 5 NaCl, 1 EGTA, 5 Mg-ATP, and 0.5 Na-GTP (pH 7.3; 280 mOsm).

For optostimulation, blue light pulses (470 nm) were delivered using an LED (Lumencor) through a 60× water-immersion objective (5 ms, 20 mW). Optogenetically evoked IPSCs (oIPSCs) were recorded at a holding potential of −10 mV. Amplitudes were calculated from the average of at least five events. Cells exhibiting detectable events ≥10 pA were classified as responsive. Picrotoxin (100μM) was bath-applied via perfusion. Data were analyzed using Clampfit software (Molecular Devices).

For proportion analysis comparing sg*Slc32a1* and sgRosa groups, Fisher’s exact test was used. The effects of picrotoxin on oIPSCs were quantified by comparing amplitudes before and 5–10 min after bath application. Statistical significance was assessed using a paired t-test. A one-tailed test was applied based on a priori directional hypothesis (decrease in amplitude).

### *Tacr3* mutagenesis in mPFC

Tacr3-Cre mice received bilateral infusion of a viral cocktail containing either 1 part EGFP-KASH: 7 parts saCas9 (AAV1-CMV-FLEX-saCas9-U6-sgTacr3) or 1 part AAV1-EGFP-KASH and 7 parts AAV1-CMV-FLEX-saCas9-U6-Rosa26. Following surgery, mice were single-housed for four weeks to allow for viral expression. After this period, mice were tested on the Resident Intruder assay as described above. Behavior was scored and analyzed as described above.

#### Slice Calcium Imaging

CRISPR-mediated mutagenesis of *Tacr3* in mPFC of isolated mice was performed analogous to the mutagenesis of mPFC sgSlc32a1 in Tac2^+^ neurons, except Tacr3^iCre^ mice (male and female, 8 -10 weeks of age) were infused with AAV1-CMV-FLEX-saCas9-U6-sgTacr3 and AAV1-Syn-FLEX-jGCaMP8m (titre 1.0 x 10^13^ GC/mL, Zweifel lab) or AAV1-CMV-FLEX-saCas9-U6-Rosa26 and AAV1-Syn-FLEX-jGCaMP8m (titre 1.0 x 10^13^ GC/mL, Zweifel lab). Behavior and surgeries were performed as described above. Coronal brain slices (200 μm) were prepared as described above. For puff application experiments, ∼10 MΩ pipettes were filled with senktide (1 μM) dissolved in ACSF containing (in mM): 126 NaCl, 2.5 KCl, 1.2 NaH₂PO₄, 1.2 MgCl₂, 11 D-glucose, 18 NaHCO₃, and 2.4 CaCl₂.

As this experiment was designed to validate sg*Tacr3* knockout efficiency, only cells exhibiting spontaneous firing were selected for analysis. Under this criterion, nonresponsive cells were interpreted as lacking functional *Tacr3* receptor expression. The puffing pipette was positioned adjacent to the cell of interest, and only one cell was analyzed per puff. Cells were included for analysis only if a visible mechanical deflection was observed during puff application. Cells showing an increase in calcium (GCaMP) signal were classified as responsive. Images were acquired using a high-speed camera (Hamamatsu OrcaFlash 4.0LT). ΔF/F was calculated using ImageJ and RStudio. For comparisons between sgTacr3 and control groups, Fisher’s exact test was used for proportion analysis. The area under the curve (0–10 s) was compared using the Mann– Whitney test.

### Statistics and Reproducibility

Data were processed and analyzed using MATLAB, Python and GraphPad Prism v9. Data points from individual animals (behavioral experiments, snRNAseq) and individual brain sections (in situ hybridization, immunohistochemistry) were plotted whenever possible. Error bars and bars on graphs represent mean ± SEM in graphs. All data were analyzed with two-tailed tests and normality was tested using the Kolmogorov-Smirnov test. If at least one group in a dataset failed the Kolmogorov-Smirnov test, then non-parametric tests were performed. Statistical significance was indicated by * p < 0.05, ** p < 0.01, *** p < 0.001, **** p < 0.0001, and n.s. no statistical significance. Details of all statistical analyses can be found in the respective figure captions. Experimenters were blinded to experimental conditions (sex, housing, viral condition, drug condition, etc) whenever possible. The testing order of animals in all experiments was randomized and test order for repeated testing was counterbalanced. Effect sizes (Cohen’s d, eta squared) were calculated using practices outlined in Psychometrica: Computation of effect sizes based on statistical test used, normality, and N of each group^121^.

## ACKNOWLEDGEMENTS

We thank Dr. Tomomi Karigo for the Tac2 shRNA and Luciferase shRNA viral plasmids; Bai Luo, Director of the Drug Discovery Core Facility at the University of Utah for their assistance in packaging shRNA plasmids into adeno-associated viral envelopes; the University of Utah Cell Imaging Core for use of the Zeiss Axioscan 7 Slide Scanner; the University of Utah Machine Shop for building the Resident Intruder assay boxes; Dr. Kristen Keefe for experimental feedback; Becky Dennis for technical assistance.

## AUTHOR CONTRIBUTIONS

M.Z. and R.E.G. conceived the project, designed experiments, and interpreted the data. R.E.G. performed stereotaxic surgeries, behavioral experiments, in situ hybridization experiments, immunohistochemistry, histology, microscopy, behavioral scoring, cell counting, statistical analyses and generated figures. Y.D. performed stereotaxic surgeries, behavioral experiments, fiber photometry experiments, behavioral scoring, cell counting, immunohistochemistry, histology, microscopy, data analyses, and generated figures. J.G. performed stereotaxic surgeries, behavioral tests, in situ hybridization experiments (FISH), behavioral scoring, histology, microscopy, and cell counting, and data analyses. D.K. performed analyses for behavioral transitions and visualization of behaviors across the Resident Intruder assay and contributed to fiber photometry experiments. S.P. performed fiber photometry experiments and analyzed the data. H.W. performed snRNAseq experiments and analyses. K.E.R. assisted with figure design, performed RNAscope™ experiments, imaging, cell counting, and histology. T.E. performed a subset of surgeries and behavioral tests. L.K. assisted with FISH experiments. A.C. and M.M.F. performed histology, cell counting, and scored behavior. Y.L. and L.S.Z. performed electrophysiology and calcium imaging validation experiments for CRISPR sgSlc32a1 and sgTacr3 experiments, analyzed data, and designed figures. T.Z. scored behavior. N.A.F. supervised snRNAseq experiments and behavioral transition analyses. M.Z. and R.E.G. wrote the manuscript and designed the figures with input from all of the authors.

## DATA AND CODE AVAILABILITY

All experimental data are available in the main text or extended data. Custom MATLAB and Python code is available on the Zelikowsky lab GitHub in the Gatlin-et-al-2026 repository (https://github.com/ZelikowskyLab/Gatlin-et-al-2026).

## REFERENCES

1. Flavell SW, Gogolla N, Lovett-Barron M, Zelikowsky M. The emergence and influence of internal states. Neuron. 2022;110(16):2545–70. Epub 20220527. doi: 10.1016/j.neuron.2022.04.030. PubMed PMID: 35643077; PMCID: PMC9391310.

2. van den Pol AN. Neuropeptide transmission in brain circuits. Neuron. 2012;76(1):98–115. doi: 10.1016/j.neuron.2012.09.014. PubMed PMID: 23040809; PMCID: PMC3918222.

3. Zelikowsky M, Ding K, Anderson DJ. Neuropeptidergic Control of an Internal Brain State Produced by Prolonged Social Isolation Stress. Cold Spring Harb Symp Quant Biol. 2018;83:97–103. Epub 20190404. doi: 10.1101/sqb.2018.83.038109. PubMed PMID: 30948452.

4. Flavell SW, Pokala N, Macosko EZ, Albrecht DR, Larsch J, Bargmann CI. Serotonin and the neuropeptide PDF initiate and extend opposing behavioral states in C. elegans. Cell. 2013;154(5):1023–35. Epub 20130822. doi: 10.1016/j.cell.2013.08.001. PubMed PMID: 23972393; PMCID: PMC3942133.

5. Banghart MR, Sabatini BL. Photoactivatable neuropeptides for spatiotemporally precise delivery of opioids in neural tissue. Neuron. 2012;73(2):249–59. doi: 10.1016/j.neuron.2011.11.016. PubMed PMID: 22284180; PMCID: PMC3282187.

6. Calhoun AJ, Pillow JW, Murthy M. Unsupervised identification of the internal states that shape natural behavior. Nature Neuroscience. 2019;22(12):2040–9. doi: 10.1038/s41593-019-0533-x.

7. Zhang SX, Kim A, Madara JC, Zhu PK, Christenson LF, Lutas A, Kalugin PN, Sunkavalli PS, Jin Y, Pal A, Tian L, Lowell BB, Andermann ML. Stochastic neuropeptide signals compete to calibrate the rate of satiation. Nature. 2025;637(8044):137–44. Epub 20241106. doi: 10.1038/s41586-024-08164-8. PubMed PMID: 39506113; PMCID: PMC11981016.

8. Soden ME, Yee JX, Zweifel LS. Circuit coordination of opposing neuropeptide and neurotransmitter signals. Nature. 2023;619(7969):332–7. Epub 20230628. doi: 10.1038/s41586-023-06246-7. PubMed PMID: 37380765; PMCID: PMC10947507.

9. Zelikowsky M, Hui M, Karigo T, Choe A, Yang B, Blanco MR, Beadle K, Gradinaru V, Deverman BE, Anderson DJ. The Neuropeptide Tac2 Controls a Distributed Brain State Induced by Chronic Social Isolation Stress. Cell. 2018;173(5):1265–79.e19. doi: 10.1016/j.cell.2018.03.037. PubMed PMID: 29775595; PMCID: PMC5967263.

10. Asede D, Doddapaneni D, Bolton MM. Amygdala Intercalated Cells: Gate Keepers and Conveyors of Internal State to the Circuits of Emotion. J Neurosci. 2022;42(49):9098–109. doi: 10.1523/jneurosci.1176-22.2022. PubMed PMID: 36639901; PMCID: PMC9761677.

11. Allen WE, Chen MZ, Pichamoorthy N, Tien RH, Pachitariu M, Luo L, Deisseroth K. Thirst regulates motivated behavior through modulation of brainwide neural population dynamics. Science. 2019;364(6437):253. Epub 20190404. doi: 10.1126/science.aav3932. PubMed PMID: 30948440; PMCID: PMC6711472.

12. Casello SM, Flores RJ, Yarur HE, Wang H, Awanyai M, Arenivar MA, Jaime-Lara RB, Bravo-Rivera H, Tejeda HA. Neuropeptide System Regulation of Prefrontal Cortex Circuitry: Implications for Neuropsychiatric Disorders. Frontiers in Neural Circuits. 2022;16. doi: 10.3389/fncir.2022.796443.

13. Yizhar O, Levy DR. The social dilemma: prefrontal control of mammalian sociability. Current Opinion in Neurobiology. 2021;68:67–75. Epub 20210204. doi: 10.1016/j.conb.2021.01.007. PubMed PMID: 33549950.

14. Chen P, Hong W. Neural Circuit Mechanisms of Social Behavior. Neuron. 2018;98(1):16–30. doi: 10.1016/j.neuron.2018.02.026. PubMed PMID: 29621486; PMCID: PMC6028944.

15. Arnsten AF. Stress signalling pathways that impair prefrontal cortex structure and function. Nat Rev Neurosci. 2009;10(6):410–22. doi: 10.1038/nrn2648. PubMed PMID: 19455173; PMCID: PMC2907136.

16. Padilla-Coreano N, Batra K, Patarino M, Chen Z, Rock RR, Zhang R, Hausmann SB, Weddington JC, Patel R, Zhang YE, Fang HS, Mishra S, LeDuke DO, Revanna J, Li H, Borio M, Pamintuan R, Bal A, Keyes LR, Libster A, Wichmann R, Mills F, Taschbach FH, Matthews GA, Curley JP, Fiete IR, Lu C, Tye KM. Cortical ensembles orchestrate social competition through hypothalamic outputs. Nature. 2022;603(7902):667–71. Epub 20220316. doi: 10.1038/s41586-022-04507-5. PubMed PMID: 35296862; PMCID: PMC9576144.

17. Cum M, Santiago Pérez JA, Wangia E, Lopez N, Wright ES, Iwata RL, Li A, Chambers AR, Padilla-Coreano N. A systematic review and meta-analysis of how social memory is studied. Sci Rep. 2024;14(1):2221. Epub 20240126. doi: 10.1038/s41598-024-52277-z. PubMed PMID: 38278973; PMCID: PMC10817899.

18. Ferrara NC, Che A, Briones B, Padilla-Coreano N, Lovett-Barron M, Opendak M. Neural Circuit Transitions Supporting Developmentally Specific Social Behavior. J Neurosci. 2023;43(45):7456–62. doi: 10.1523/jneurosci.1377-23.2023. PubMed PMID: 37940586; PMCID: PMC10634550.

19. George A, Padilla-Coreano N, Opendak M. For neuroscience, social history matters. Neuropsychopharmacology. 2023;48(7):979–80. Epub 20230315. doi: 10.1038/s41386-023-01566-8. PubMed PMID: 36922626; PMCID: PMC10209051.

20. Kuniishi H, Nakatake Y, Sekiguchi M, Yamada M. Adolescent social isolation induces distinct changes in the medial and lateral OFC-BLA synapse and social and emotional alterations in adult mice. Neuropsychopharmacology. 2022;47(9):1597–607. Epub 20220613. doi: 10.1038/s41386-022-01358-6. PubMed PMID: 35697823; PMCID: PMC9283446.

21. Yizhar O, Fenno LE, Prigge M, Schneider F, Davidson TJ, O’Shea DJ, Sohal VS, Goshen I, Finkelstein J, Paz JT, Stehfest K, Fudim R, Ramakrishnan C, Huguenard JR, Hegemann P, Deisseroth K. Neocortical excitation/inhibition balance in information processing and social dysfunction. Nature. 2011;477(7363):171–8. Epub 20110727. doi: 10.1038/nature10360. PubMed PMID: 21796121; PMCID: PMC4155501.

22. Biro L, Miskolczi C, Szebik H, Bruzsik B, Varga ZK, Szente L, Toth M, Halasz J, Mikics E. Post-weaning social isolation in male mice leads to abnormal aggression and disrupted network organization in the prefrontal cortex: Contribution of parvalbumin interneurons with or without perineuronal nets. Neurobiol Stress. 2023;25:100546. Epub 20230529. doi: 10.1016/j.ynstr.2023.100546. PubMed PMID: 37323648; PMCID: PMC10265620.

23. Li X, Sun H, Zhu Y, Wang F, Wang X, Han L, Cui D, Luo D, Zhai Y, Zhuo L, Xu X, Yang J, Li Y. Dysregulation of prefrontal parvalbumin interneurons leads to adult aggression induced by social isolation stress during adolescence. Front Mol Neurosci. 2022;15:1010152. Epub 20221004. doi: 10.3389/fnmol.2022.1010152. PubMed PMID: 36267698; PMCID: PMC9577330.

24. Chaibi I, Bennis M, Ba-M’Hamed S. GABA-A receptor signaling in the anterior cingulate cortex modulates aggression and anxiety-related behaviors in socially isolated mice. Brain Res. 2021;1762:147440. Epub 20210318. doi: 10.1016/j.brainres.2021.147440. PubMed PMID: 33745922.

25. Wang ZJ, Shwani T, Liu J, Zhong P, Yang F, Schatz K, Zhang F, Pralle A, Yan Z. Molecular and cellular mechanisms for differential effects of chronic social isolation stress in males and females. Mol Psychiatry. 2022;27(7):3056–68. Epub 20220421. doi: 10.1038/s41380-022-01574-y. PubMed PMID: 35449296; PMCID: PMC9615910.

26. Heukelum SV, Geers FE, Tulva K, van Dulm S, Beckmann CF, Buitelaar JK, Glennon JC, Vogt BA, Havenith MN, França ASC. Structural Degradation in Midcingulate Cortex Is Associated with Pathological Aggression in Mice. Brain Sci. 2021;11(7). Epub 20210629. doi: 10.3390/brainsci11070868. PubMed PMID: 34209993; PMCID: PMC8301779.

27. Bicks LK, Yamamuro K, Flanigan ME, Kim JM, Kato D, Lucas EK, Koike H, Peng MS, Brady DM, Chandrasekaran S, Norman KJ, Smith MR, Clem RL, Russo SJ, Akbarian S, Morishita H. Prefrontal parvalbumin interneurons require juvenile social experience to establish adult social behavior. Nat Commun. 2020;11(1):1003. Epub 20200221. doi: 10.1038/s41467-020-14740-z. PubMed PMID: 32081848; PMCID: PMC7035248.

28. Florido A, Velasco ER, Soto-Faguás CM, Gomez-Gomez A, Perez-Caballero L, Molina P, Nadal R, Pozo OJ, Saura CA, Andero R. Sex differences in fear memory consolidation via Tac2 signaling in mice. Nat Commun. 2021;12(1):2496. Epub 20210503. doi: 10.1038/s41467-021-22911-9. PubMed PMID: 33941789; PMCID: PMC8093426.

29. Zhao ZD, Han X, Chen R, Liu Y, Bhattacherjee A, Chen W, Zhang Y. A molecularly defined D1 medium spiny neuron subtype negatively regulates cocaine addiction. Sci Adv. 2022;8(32):eabn3552. Epub 20220812. doi: 10.1126/sciadv.abn3552. PubMed PMID: 35960793; PMCID: PMC9374336.

30. Hook M, Xu F, Terenina E, Zhao W, Starlard-Davenport A, Mormede P, Jones BC, Mulligan MK, Lu L. Exploring the involvement of Tac2 in the mouse hippocampal stress response through gene networking. Gene. 2019;696:176–85. Epub 20190212. doi: 10.1016/j.gene.2019.02.013. PubMed PMID: 30769143; PMCID: PMC6754182.

31. Andero R, Dias BG, Ressler KJ. A role for Tac2, NkB, and Nk3 receptor in normal and dysregulated fear memory consolidation. Neuron. 2014;83(2):444–54. Epub 20140626. doi: 10.1016/j.neuron.2014.05.028. PubMed PMID: 24976214; PMCID: PMC4103970.

32. Andero R, Daniel S, Guo JD, Bruner RC, Seth S, Marvar PJ, Rainnie D, Ressler KJ. Amygdala-Dependent Molecular Mechanisms of the Tac2 Pathway in Fear Learning. Neuropsychopharmacology. 2016;41(11):2714–22. Epub 20160526. doi: 10.1038/npp.2016.77. PubMed PMID: 27238620; PMCID: PMC5026739.

33. Boyle CA, Hu B, Quaintance KL, Mastrud MR, Lei S. Ionic signalling mechanisms involved in neurokinin-3 receptor-mediated augmentation of fear-potentiated startle response in the basolateral amygdala. J Physiol. 2022;600(19):4325–45. Epub 20220913. doi: 10.1113/jp283433. PubMed PMID: 36030507; PMCID: PMC9529888.

34. Asahina K, Watanabe K, Duistermars BJ, Hoopfer E, González CR, Eyjólfsdóttir EA, Perona P, Anderson DJ. Tachykinin-expressing neurons control male-specific aggressive arousal in Drosophila. Cell. 2014;156(1-2):221–35. doi: 10.1016/j.cell.2013.11.045. PubMed PMID: 24439378; PMCID: PMC3978814.

35. Al Abed AS, Reynolds NJ, Dehorter N. A Second Wave for the Neurokinin Tac2 Pathway in Brain Research. Biol Psychiatry. 2021;90(3):156–64. Epub 20210223. doi: 10.1016/j.biopsych.2021.02.016. PubMed PMID: 33867115.

36. Asahina K, Zelikowsky M. Comparative perspectives on neuropeptide function and social isolation. Biological Psychiatry. 2025. doi: 10.1016/j.biopsych.2025.01.019.

37. Ebner K, Sartori SB, Singewald N. Tachykinin receptors as therapeutic targets in stress-related disorders. Curr Pharm Des. 2009;15(14):1647–74. doi: 10.2174/138161209788168074. PubMed PMID: 19442179.

38. Stacey AE, Woodhall GL, Jones RS. Neurokinin-receptor-mediated depolarization of cortical neurons elicits an increase in glutamate release at excitatory synapses. Eur J Neurosci. 2002;16(10):1896–906. doi: 10.1046/j.1460-9568.2002.02266.x. PubMed PMID: 12453053.

39. Gallopin T, Geoffroy H, Rossier J, Lambolez B. Cortical sources of CRF, NKB, and CCK and their effects on pyramidal cells in the neocortex. Cereb Cortex. 2006;16(10):1440–52. Epub 20051207. doi: 10.1093/cercor/bhj081. PubMed PMID: 16339088.

40. Myers T, Birmingham EA, Rhoads BT, McGrath AG, Miles NA, Schuldt CB, Briand LA. Post-weaning social isolation alters sociability in a sex-specific manner. Front Behav Neurosci. 2024;18:1444596. Epub 20240829. doi: 10.3389/fnbeh.2024.1444596. PubMed PMID: 39267986; PMCID: PMC11390411.

41. Matthews GA, Nieh EH, Vander Weele CM, Halbert SA, Pradhan RV, Yosafat AS, Glober GF, Izadmehr EM, Thomas RE, Lacy GD. Dorsal raphe dopamine neurons represent the experience of social isolation. Cell. 2016;164(4):617–31.

42. Tan T, Wang W, Liu T, Zhong P, Conrow-Graham M, Tian X, Yan Z. Neural circuits and activity dynamics underlying sex-specific effects of chronic social isolation stress. Cell Rep. 2021;34(12):108874. doi: 10.1016/j.celrep.2021.108874. PubMed PMID: 33761364.

43. Pinna G, Dong E, Matsumoto K, Costa E, Guidotti A. In socially isolated mice, the reversal of brain allopregnanolone down-regulation mediates the anti-aggressive action of fluoxetine. Proceedings of the National Academy of Sciences. 2003;100(4):2035–40.

44. Morè L. Intra-female aggression in the mouse (Mus musculus domesticus) is linked to the estrous cycle regularity but not to ovulation. Aggress Behav. 2008;34(1):46–50. doi: 10.1002/ab.20217. PubMed PMID: 17647257.

45. Kuchiiwa T, Kuchiiwa S. Evaluation of aggressiveness of female mice using a semi-automated apparatus for measurement of aggressive biting behavior toward an inanimate object. Journal of Neuroscience Methods. 2016;257:179–84. doi: 10.1016/j.jneumeth.2015.10.005.

46. Liu D, Rahman M, Johnson A, Tsutsui-Kimura I, Pena N, Talay M, Logeman BL, Finkbeiner S, Choi S, Capo-Battaglia A, Abdus-Saboor I, Ginty DD, Uchida N, Watabe-Uchida M, Dulac C. A Hypothalamic Circuit Underlying the Dynamic Control of Social Homeostasis. bioRxiv. 2023. Epub 20230519. doi: 10.1101/2023.05.19.540391. PubMed PMID: 37293031; PMCID: PMC10245688.

47. Grammer J, Dryer A, Tweed C, Conoscenti M, Zelikowsky M. Extended amygdala orchestrates social motivation in socially isolated mice. 2026.

48. Liu D, Rahman M, Johnson A, Amo R, Tsutsui-Kimura I, Sullivan ZA, Pena N, Talay M, Logeman BL, Finkbeiner S, Qian L, Choi S, Capo-Battaglia A, Abdus-Saboor I, Ginty DD, Uchida N, Watabe-Uchida M, Dulac C. A hypothalamic circuit underlying the dynamic control of social homeostasis. Nature. 2025;640(8060):1000–10. doi: 10.1038/s41586-025-08617-8.

49. Thurmond JB. Technique for producing and measuring territorial aggression using laboratory mice. Physiol Behav. 1975;14(6):879–81. doi: 10.1016/0031-9384(75)90086-4. PubMed PMID: 1237905.

50. Blanchard RJ, Wall PM, Blanchard DC. Problems in the study of rodent aggression. Hormones and Behavior. 2003;44(3):161–70. doi: 10.1016/S0018-506X(03)00127-2.

51. Oliveira VEM, Evrard F, Faure MC, Bakker J. Social isolation and aggression training lead to escalated aggression and hypothalamus-pituitary-gonad axis hyperfunction in mice. Neuropsychopharmacology. 2024;49(8):1266–75. Epub 20240209. doi: 10.1038/s41386-024-01808-3. PubMed PMID: 38337026; PMCID: PMC11224373.

52. Chen C, Altafi M, Corbu M-A, Trenk A, van den Munkhof H, Weineck K, Bender F, Carus-Cadavieco M, Bakhareva A, Korotkova T, Ponomarenko A. The dynamic state of a prefrontal–hypothalamic–midbrain circuit commands behavioral transitions. Nature Neuroscience. 2024;27(5):952–63. doi: 10.1038/s41593-024-01598-3.

53. Ahmadlou M, Houba JHW, van Vierbergen JFM, Giannouli M, Gimenez GA, van Weeghel C, Darbanfouladi M, Shirazi MY, Dziubek J, Kacem M, de Winter F, Heimel JA. A cell type-specific cortico-subcortical brain circuit for investigatory and novelty-seeking behavior. Science. 2021;372(6543). doi: 10.1126/science.abe9681. PubMed PMID: 33986154.

54. Minakuchi T, Guthman EM, Acharya P, Hinson J, Fleming W, Witten IB, Oline SN, Falkner AL. Independent inhibitory control mechanisms for aggressive motivation and action. Nat Neurosci. 2024;27(4):702–15. Epub 20240212. doi: 10.1038/s41593-023-01563-6. PubMed PMID: 38347201.

55. Grammer J, Valles R, Bowles A, Zelikowsky M. An integrative assay for measuring social aversion and motivation in freely behaving mice. Cell Rep Methods. 2025;5(8):101108. Epub 20250717. doi: 10.1016/j.crmeth.2025.101108. PubMed PMID: 40680732; PMCID: PMC12461583.

56. Biro L, Sipos E, Bruzsik B, Farkas I, Zelena D, Balazsfi D, Toth M, Haller J. Task Division within the Prefrontal Cortex: Distinct Neuron Populations Selectively Control Different Aspects of Aggressive Behavior via the Hypothalamus. J Neurosci. 2018;38(17):4065–75. Epub 2018/03/01. doi: 10.1523/JNEUROSCI.3234-17.2018. PubMed PMID: 29487128; PMCID: PMC6596023.

57. Li X, Xiong L, Li Y. The role of the prefrontal cortex in modulating aggression in humans and rodents. Behav Brain Res. 2025;476:115285. Epub 20241005. doi: 10.1016/j.bbr.2024.115285. PubMed PMID: 39369825.

58. Halász J, Tóth M, Kalló I, Liposits Z, Haller J. The activation of prefrontal cortical neurons in aggression— A double labeling study. Behavioural Brain Research. 2006;175(1):166–75. doi: 10.1016/j.bbr.2006.08.019.

59. Harris JA, Hirokawa KE, Sorensen SA, Gu H, Mills M, Ng LL, Bohn P, Mortrud M, Ouellette B, Kidney J, Smith KA, Dang C, Sunkin S, Bernard A, Oh SW, Madisen L, Zeng H. Anatomical characterization of Cre driver mice for neural circuit mapping and manipulation. Front Neural Circuits. 2014;8:76. Epub 20140710. doi: 10.3389/fncir.2014.00076. PubMed PMID: 25071457; PMCID: PMC4091307.

60. Guo Z, Yin L, Diaz V, Dai B, Osakada T, Lischinsky JE, Chien J, Yamaguchi T, Urtecho A, Tong X, Chen ZS, Lin D. Neural dynamics in the limbic system during male social behaviors. Neuron. 2023;111(20):3288–306.e4. Epub 20230815. doi: 10.1016/j.neuron.2023.07.011. PubMed PMID: 37586365; PMCID: PMC10592239.

61. Yang T, Bayless DW, Wei Y, Landayan D, Marcelo IM, Wang Y, DeNardo LA, Luo L, Druckmann S, Shah NM. Hypothalamic neurons that mirror aggression. Cell. 2023;186(6):1195–211.e19. doi: 10.1016/j.cell.2023.01.022.

62. Yamamuro K, Yoshino H, Ogawa Y, Okamura K, Nishihata Y, Makinodan M, Saito Y, Kishimoto T. Juvenile Social Isolation Enhances the Activity of Inhibitory Neuronal Circuits in the Medial Prefrontal Cortex. Front Cell Neurosci. 2020;14:105. Epub 20200512. doi: 10.3389/fncel.2020.00105. PubMed PMID: 32477068; PMCID: PMC7235301.

63. Wang F, Flanagan J, Su N, Wang LC, Bui S, Nielson A, Wu X, Vo HT, Ma XJ, Luo Y. RNAscope: a novel in situ RNA analysis platform for formalin-fixed, paraffin-embedded tissues. J Mol Diagn. 2012;14(1):22–9. doi: 10.1016/j.jmoldx.2011.08.002. PubMed PMID: 22166544; PMCID: PMC3338343.

64. Yao Z, van Velthoven CTJ, Nguyen TN, Goldy J, Sedeno-Cortes AE, Baftizadeh F, Bertagnolli D, Casper T, Chiang M, Crichton K, Ding S-L, Fong O, Garren E, Glandon A, Gouwens NW, Gray J, Graybuck LT, Hawrylycz MJ, Hirschstein D, Kroll M, Lathia K, Lee C, Levi B, McMillen D, Mok S, Pham T, Ren Q, Rimorin C, Shapovalova N, Sulc J, Sunkin SM, Tieu M, Torkelson A, Tung H, Ward K, Dee N, Smith KA, Tasic B, Zeng H. A taxonomy of transcriptomic cell types across the isocortex and hippocampal formation. Cell. 2021;184(12):3222–41.e26. doi: 10.1016/j.cell.2021.04.021.

65. Tasic B, Yao Z, Graybuck LT, Smith KA, Nguyen TN, Bertagnolli D, Goldy J, Garren E, Economo MN, Viswanathan S, Penn O, Bakken T, Menon V, Miller J, Fong O, Hirokawa KE, Lathia K, Rimorin C, Tieu M, Larsen R, Casper T, Barkan E, Kroll M, Parry S, Shapovalova NV, Hirschstein D, Pendergraft J, Sullivan HA, Kim TK, Szafer A, Dee N, Groblewski P, Wickersham I, Cetin A, Harris JA, Levi BP, Sunkin SM, Madisen L, Daigle TL, Looger L, Bernard A, Phillips J, Lein E, Hawrylycz M, Svoboda K, Jones AR, Koch C, Zeng H. Shared and distinct transcriptomic cell types across neocortical areas. Nature. 2018;563(7729):72–8. doi: 10.1038/s41586-018-0654-5.

66. Smith SJ, Sümbül U, Graybuck LT, Collman F, Seshamani S, Gala R, Gliko O, Elabbady L, Miller JA, Bakken TE, Rossier J, Yao Z, Lein E, Zeng H, Tasic B, Hawrylycz M. Single-cell transcriptomic evidence for dense intracortical neuropeptide networks. Elife. 2019;8. Epub 20191111. doi: 10.7554/eLife.47889. PubMed PMID: 31710287; PMCID: PMC6881117.

67. Beier KT, Steinberg EE, DeLoach KE, Xie S, Miyamichi K, Schwarz L, Gao XJ, Kremer EJ, Malenka RC, Luo L. Circuit Architecture of VTA Dopamine Neurons Revealed by Systematic Input-Output Mapping. Cell. 2015;162(3):622–34. doi: 10.1016/j.cell.2015.07.015. PubMed PMID: 26232228; PMCID: PMC4522312.

68. Noorman M, Hulse BK, Jayaraman V, Romani S, Hermundstad AM. Maintaining and updating accurate internal representations of continuous variables with a handful of neurons. Nat Neurosci. 2024;27(11):2207–17. Epub 20241003. doi: 10.1038/s41593-024-01766-5. PubMed PMID: 39363052; PMCID: PMC11537979.

69. Seabrook GR, Bowery BJ, Hill RG. Pharmacology of tachykinin receptors on neurones in the ventral tegmental area of rat brain slices. Eur J Pharmacol. 1995;273(1-2):113–9. doi: 10.1016/0014-2999(94)00681-v. PubMed PMID: 7537676.

70. Pieri M, Severini C, Amadoro G, Carunchio I, Barbato C, Ciotti MT, Zona C. AMPA receptors are modulated by tachykinins in rat cerebellum neurons. J Neurophysiol. 2005;94(4):2484–90. doi: 10.1152/jn.00436.2005. PubMed PMID: 16160091.

71. Mawe GM. Tachykinins as mediators of slow EPSPs in guinea-pig gall-bladder ganglia: involvement of neurokinin-3 receptors. J Physiol. 1995;485 (Pt 2)(Pt 2):513–24. doi: 10.1113/jphysiol.1995.sp020747. PubMed PMID: 7545233; PMCID: PMC1158010.

72. Hunker AC, Soden ME, Krayushkina D, Heymann G, Awatramani R, Zweifel LS. Conditional Single Vector CRISPR/SaCas9 Viruses for Efficient Mutagenesis in the Adult Mouse Nervous System. Cell Rep. 2020;30(12):4303–16.e6. doi: 10.1016/j.celrep.2020.02.092. PubMed PMID: 32209486; PMCID: PMC7212805.

73. Swiech L, Heidenreich M, Banerjee A, Habib N, Li Y, Trombetta J, Sur M, Zhang F. In vivo interrogation of gene function in the mammalian brain using CRISPR-Cas9. Nature biotechnology. 2015;33(1):102–6.

74. Zambrowicz BP, Imamoto A, Fiering S, Herzenberg LA, Kerr WG, Soriano P. Disruption of overlapping transcripts in the ROSA βgeo 26 gene trap strain leads to widespread expression of β-galactosidase in mouse embryos and hematopoietic cells. Proceedings of the National Academy of Sciences. 1997;94(8):3789–94.

75. Gerachshenko T, Blackmer T, Yoon EJ, Bartleson C, Hamm HE, Alford S. Gbetagamma acts at the C terminus of SNAP-25 to mediate presynaptic inhibition. Nat Neurosci. 2005;8(5):597–605. Epub 20050417. doi: 10.1038/nn1439. PubMed PMID: 15834421.

76. Roth BL. DREADDs for Neuroscientists. Neuron. 2016;89(4):683–94. doi: 10.1016/j.neuron.2016.01.040. PubMed PMID: 26889809; PMCID: PMC4759656.

77. Yao Z, van Velthoven CTJ, Nguyen TN, Goldy J, Sedeno-Cortes AE, Baftizadeh F, Bertagnolli D, Casper T, Chiang M, Crichton K, Ding SL, Fong O, Garren E, Glandon A, Gouwens NW, Gray J, Graybuck LT, Hawrylycz MJ, Hirschstein D, Kroll M, Lathia K, Lee C, Levi B, McMillen D, Mok S, Pham T, Ren Q, Rimorin C, Shapovalova N, Sulc J, Sunkin SM, Tieu M, Torkelson A, Tung H, Ward K, Dee N, Smith KA, Tasic B, Zeng H. A taxonomy of transcriptomic cell types across the isocortex and hippocampal formation. Cell. 2021;184(12):3222–41.e26. Epub 20210517. doi: 10.1016/j.cell.2021.04.021. PubMed PMID: 34004146; PMCID: PMC8195859.

78. Muppirala AN, Mitchell PE, Courtney E, Debnath S, D’Alessandro LR, Mani N, Dadabhoy M, Robinson A, Díaz-Muñoz C, Paraskevopoulou MD, D’Amato M, Gould TW, Rao M. Tachykinin signaling defines distinct populations of glia in the enteric nervous system. Neuron. 2026;114(6):1066–82.e8. doi: 10.1016/j.neuron.2025.11.030.

79. Musella ML, Sammari M, Ajal C, Le Cor A, Russier M, Debanne D, Incontro S. Somatostatin released by single O-LM interneurons slowly inhibits intrinsic neuronal excitability in a paracrine and autocrine manner. 2025.

80. Thirumalai V, Marder E. Colocalized Neuropeptides Activate a Central Pattern Generator by Acting on Different Circuit Targets. The Journal of Neuroscience. 2002;22(5):1874–82. doi: 10.1523/jneurosci.22-05-01874.2002.

81. Marquina-Solis J, Feng L, Vandewyer E, Beets I, Hawk J, Colón-Ramos DA, Yu J, Fox BW, Schroeder FC, Bargmann CI. Antagonism between neuropeptides and monoamines in a distributed circuit for pathogen avoidance. Cell Rep. 2024;43(4):114042. Epub 20240403. doi: 10.1016/j.celrep.2024.114042. PubMed PMID: 38573858; PMCID: PMC11063628.

82. Ross AP, McCann KE, Larkin TE, Song Z, Grieb ZA, Huhman KL, Albers HE. Sex-dependent effects of social isolation on the regulation of arginine-vasopressin (AVP) V1a, oxytocin (OT) and serotonin (5HT) 1a receptor binding and aggression. Horm Behav. 2019;116:104578. Epub 20190829. doi: 10.1016/j.yhbeh.2019.104578. PubMed PMID: 31449813; PMCID: PMC6885541.

83. Blanchard RJ, Blanchard DC. Aggressive behavior in the rat. Behavioral biology. 1977;21(2):197–224.

84. Hiller I. The White-tailed Deer: Texas A&M University Press; 1996.

85. Soden ME, Yee JX, Cuevas B, Rastani A, Elum J, Zweifel LS. Distinct Encoding of Reward and Aversion by Peptidergic BNST Inputs to the VTA. Front Neural Circuits. 2022;16:918839. Epub 20220704. doi: 10.3389/fncir.2022.918839. PubMed PMID: 35860212; PMCID: PMC9289195.

86. Kim DI, Kang SJ, Jhang J, Jo YS, Park S, Ye M, Pyeon GH, Im GH, Kim SG, Han S. Encoding opposing valences through frequency-dependent transmitter switching in single peptidergic neurons. bioRxiv. 2024. Epub 20241109. doi: 10.1101/2024.11.09.622790. PubMed PMID: 39574736; PMCID: PMC11581014.

87. Wang H, Flores RJ, Yarur HE, Limoges A, Bravo-Rivera H, Casello SM, Loomba N, Enriquez-Traba J, Arenivar M, Wang Q, Ganley R, Ramakrishnan C, Fenno LE, Kim Y, Deisseroth K, Or G, Dong C, Hoon MA, Tian L, Tejeda HA. Prefrontal cortical dynorphin peptidergic transmission constrains threat-driven behavioral and network states. Neuron. 2024;112(12):2062–78.e7. Epub 20240412. doi: 10.1016/j.neuron.2024.03.015. PubMed PMID: 38614102; PMCID: PMC11250624.

88. Chen C, Altafi M, Corbu MA, Trenk A, van den Munkhof H, Weineck K, Bender F, Carus-Cadavieco M, Bakhareva A, Korotkova T, Ponomarenko A. The dynamic state of a prefrontal-hypothalamic-midbrain circuit commands behavioral transitions. Nat Neurosci. 2024;27(5):952–63. Epub 20240318. doi: 10.1038/s41593-024-01598-3. PubMed PMID: 38499854; PMCID: PMC11089001.

89. Takahashi A, Nagayasu K, Nishitani N, Kaneko S, Koide T. Control of intermale aggression by medial prefrontal cortex activation in the mouse. PLoS One. 2014;9(4):e94657. Epub 2014/04/18. doi: 10.1371/journal.pone.0094657. PubMed PMID: 24740241; PMCID: PMC3989250.

90. van Heukelum S, Tulva K, Geers FE, van Dulm S, Ruisch IH, Mill J, Viana JF, Beckmann CF, Buitelaar JK, Poelmans G, Glennon JC, Vogt BA, Havenith MN, França ASC. A central role for anterior cingulate cortex in the control of pathological aggression. Current Biology. 2021;31(11):2321–33.e5. doi: 10.1016/j.cub.2021.03.062.

91. Siever LJ. Neurobiology of aggression and violence. Am J Psychiatry. 2008;165(4):429-42. Epub 20080317. doi: 10.1176/appi.ajp.2008.07111774. PubMed PMID: 18346997; PMCID: PMC4176893.

92. Li X, Xiong L, Li Y. The role of the prefrontal cortex in modulating aggression in humans and rodents. Behavioural Brain Research. 2024:115285.

93. Biro L, Toth M, Sipos E, Bruzsik B, Tulogdi A, Bendahan S, Sandi C, Haller J. Structural and functional alterations in the prefrontal cortex after post-weaning social isolation: relationship with species-typical and deviant aggression. Brain Struct Funct. 2017;222(4):1861–75. Epub 20160923. doi: 10.1007/s00429-016-1312-z. PubMed PMID: 27664119.

94. Lee S, Kruglikov I, Huang ZJ, Fishell G, Rudy B. A disinhibitory circuit mediates motor integration in the somatosensory cortex. Nature Neuroscience. 2013;16(11):1662–70. doi: 10.1038/nn.3544.

95. Pfeffer CK, Xue M, He M, Huang ZJ, Scanziani M. Inhibition of inhibition in visual cortex: the logic of connections between molecularly distinct interneurons. Nature Neuroscience. 2013;16(8):1068–76. doi: 10.1038/nn.3446.

96. Kullander K, Topolnik L. Cortical disinhibitory circuits: cell types, connectivity and function. Trends in Neurosciences. 2021;44(8):643–57. doi: 10.1016/j.tins.2021.04.009.

97. Pi H-J, Hangya B, Kvitsiani D, Sanders JI, Huang ZJ, Kepecs A. Cortical interneurons that specialize in disinhibitory control. Nature. 2013;503(7477):521–4. doi: 10.1038/nature12676.

98. Anastasiades PG, Carter AG. Circuit organization of the rodent medial prefrontal cortex. Trends Neurosci. 2021;44(7):550–63. Epub 20210507. doi: 10.1016/j.tins.2021.03.006. PubMed PMID: 33972100; PMCID: PMC8222144.

99. Dellal S, Zurita H, Kruglikov I, Valero M, Abad-Perez P, Geron E, Meng JH, Prönneke A, Hanson JL, Mir E, Ongaro M, Wang X-J, Buzsáki G, Machold R, Rudy B. Inhibitory and disinhibitory VIP IN-mediated circuits in neocortex. 2025.

100. Apicella AJ, Marchionni I. VIP-Expressing GABAergic Neurons: Disinhibitory vs. Inhibitory Motif and Its Role in Communication Across Neocortical Areas. Frontiers in Cellular Neuroscience. 2022;16. doi: 10.3389/fncel.2022.811484.

101. Topolnik L, Tamboli S. The role of inhibitory circuits in hippocampal memory processing. Nature Reviews Neuroscience. 2022;23(8):476–92. doi: 10.1038/s41583-022-00599-0.

102. Garcia-Junco-Clemente P, Ikrar T, Tring E, Xu X, Ringach DL, Trachtenberg JT. An inhibitory pull–push circuit in frontal cortex. Nature Neuroscience. 2017;20(3):389–92. doi: 10.1038/nn.4483.

103. Arora S, Saarloos I, Kooistra R, Van De Bospoort R, Verhage M, Toonen RF. SNAP-25 gene family members differentially support secretory vesicle fusion. Journal of Cell Science. 2017;130(11):1877–89.

104. Abramian A, Hoogstraaten RI, Murphy FH, McDaniel KF, Toonen RF, Verhage M. Rabphilin-3A negatively regulates neuropeptide release, through its SNAP25 interaction. Elife. 2024;13. Epub 20241016. doi: 10.7554/eLife.95371. PubMed PMID: 39412498; PMCID: PMC11483123.

105. Hashikawa K, Hashikawa Y, Tremblay R, Zhang J, Feng JE, Sabol A, Piper WT, Lee H, Rudy B, Lin D. Esr1+ cells in the ventromedial hypothalamus control female aggression. Nature Neuroscience. 2017;20(11):1580–90. doi: 10.1038/nn.4644.

106. Liu M, Kim D-W, Zeng H, Anderson DJ. Make war not love: The neural substrate underlying a state-dependent switch in female social behavior. Neuron. 2022;110(5):841–56.e6. doi: 10.1016/j.neuron.2021.12.002.

107. Hashikawa K, Hashikawa Y, Lischinsky J, Lin D. The Neural Mechanisms of Sexually Dimorphic Aggressive Behaviors. Trends in Genetics. 2018;34(10):755–76. doi: 10.1016/j.tig.2018.07.001.

108. Florido A, Moreno E, Canela EI, Andero R. Nk3R blockade has sex-divergent effects on memory in mice. Biol Sex Differ. 2022;13(1):28. Epub 20220611. doi: 10.1186/s13293-022-00437-z. PubMed PMID: 35690790; PMCID: PMC9188709.

109. Florido A, Velasco ER, Romero LR, Acharya N, Marin Blasco IJ, Nabás JF, Perez-Caballero L, Rivero G, Olabarrieta E, Nuñez-delMoral A, González-Parra JA, Porta-Casteràs D, Cano M, Steward T, Antony MS, Cardoner N, Torrubia R, Jackson AC, Fullana MA, Andero R. Sex differences in neural projections of fear memory processing in mice and humans. Sci Adv. 2024;10(28):eadk3365. Epub 20240710. doi: 10.1126/sciadv.adk3365. PubMed PMID: 38985873; PMCID: PMC11235172.

110. Acharya N, Nabás JF, Andero R. NK3R antagonism reduces fear expression in a PTSD-like model of female mice. Brain Medicine. 2025;1(4):73–4. doi: 10.61373/bm025l.0035.

111. Borrow AP, Bales NJ, Stover SA, Handa RJ. Chronic Variable Stress Induces Sex-Specific Alterations in Social Behavior and Neuropeptide Expression in the Mouse. Endocrinology. 2018;159(7):2803–14. doi: 10.1210/en.2018-00217.

112. McCall NM, Sprow GM, Delpire E, Thiele TE, Kash TL, Pleil KE. Effects of sex and deletion of neuropeptide Y2 receptors from GABAergic neurons on affective and alcohol drinking behaviors in mice. Frontiers in Integrative Neuroscience. 2013;7. doi: 10.3389/fnint.2013.00100.

113. Everitt BJ, Robbins TW. Drug Addiction: Updating Actions to Habits to Compulsions Ten Years On. Annu Rev Psychol. 2016;67:23–50. Epub 20150807. doi: 10.1146/annurev-psych-122414-033457. PubMed PMID: 26253543.

114. Davidson RJ, Putnam KM, Larson CL. Dysfunction in the Neural Circuitry of Emotion Regulation--A Possible Prelude to Violence. Science. 2000;289(5479):591–4. doi: 10.1126/science.289.5479.591.

115. Cai H, Haubensak W, Anthony TE, Anderson DJ. Central amygdala PKC-δ+ neurons mediate the influence of multiple anorexigenic signals. Nature neuroscience. 2014;17(9):1240–8.

116. Wu YE, Pan L, Zuo Y, Li X, Hong W. Detecting Activated Cell Populations Using Single-Cell RNA-Seq. Neuron. 2017;96(2):313–29.e6. doi: 10.1016/j.neuron.2017.09.026. PubMed PMID: 29024657.

117. Ayhan F, Douglas C, Lega BC, Konopka G. Nuclei isolation from surgically resected human hippocampus. STAR Protoc. 2021;2(4):100844. Epub 20210916. doi: 10.1016/j.xpro.2021.100844. PubMed PMID: 34585170; PMCID: PMC8455478.

118. Walker H, Frost NA. Protocol for the generation of single-nuclei RNA-seq libraries and quantification of heterogeneous cell types activated during social interaction. STAR Protocols. 2024;5(4):103395. doi: 10.1016/j.xpro.2024.103395.

119. Hao Y, Hao S, Andersen-Nissen E, Mauck WM, 3rd, Zheng S, Butler A, Lee MJ, Wilk AJ, Darby C, Zager M, Hoffman P, Stoeckius M, Papalexi E, Mimitou EP, Jain J, Srivastava A, Stuart T, Fleming LM, Yeung B, Rogers AJ, McElrath JM, Blish CA, Gottardo R, Smibert P, Satija R. Integrated analysis of multimodal single-cell data. Cell. 2021;184(13):3573–87.e29. Epub 20210531. doi: 10.1016/j.cell.2021.04.048. PubMed PMID: 34062119; PMCID: PMC8238499.

120. Bhattacherjee A, Djekidel MN, Chen R, Chen W, Tuesta LM, Zhang Y. Cell type-specific transcriptional programs in mouse prefrontal cortex during adolescence and addiction. Nat Commun. 2019;10(1):4169. Epub 20190913. doi: 10.1038/s41467-019-12054-3. PubMed PMID: 31519873; PMCID: PMC6744514.

121. Lenhard WL, A. Computation of effect sizes. Psychometrica. 2022. doi: 10.13140/RG.2.2.17823.92329.

