## Supplementary material for "Opposing signaling molecules exert cortical control over dissociable behaviors that tile an aggressive state": supplmental figures and legends

### EXTENDED DATA FIGURES

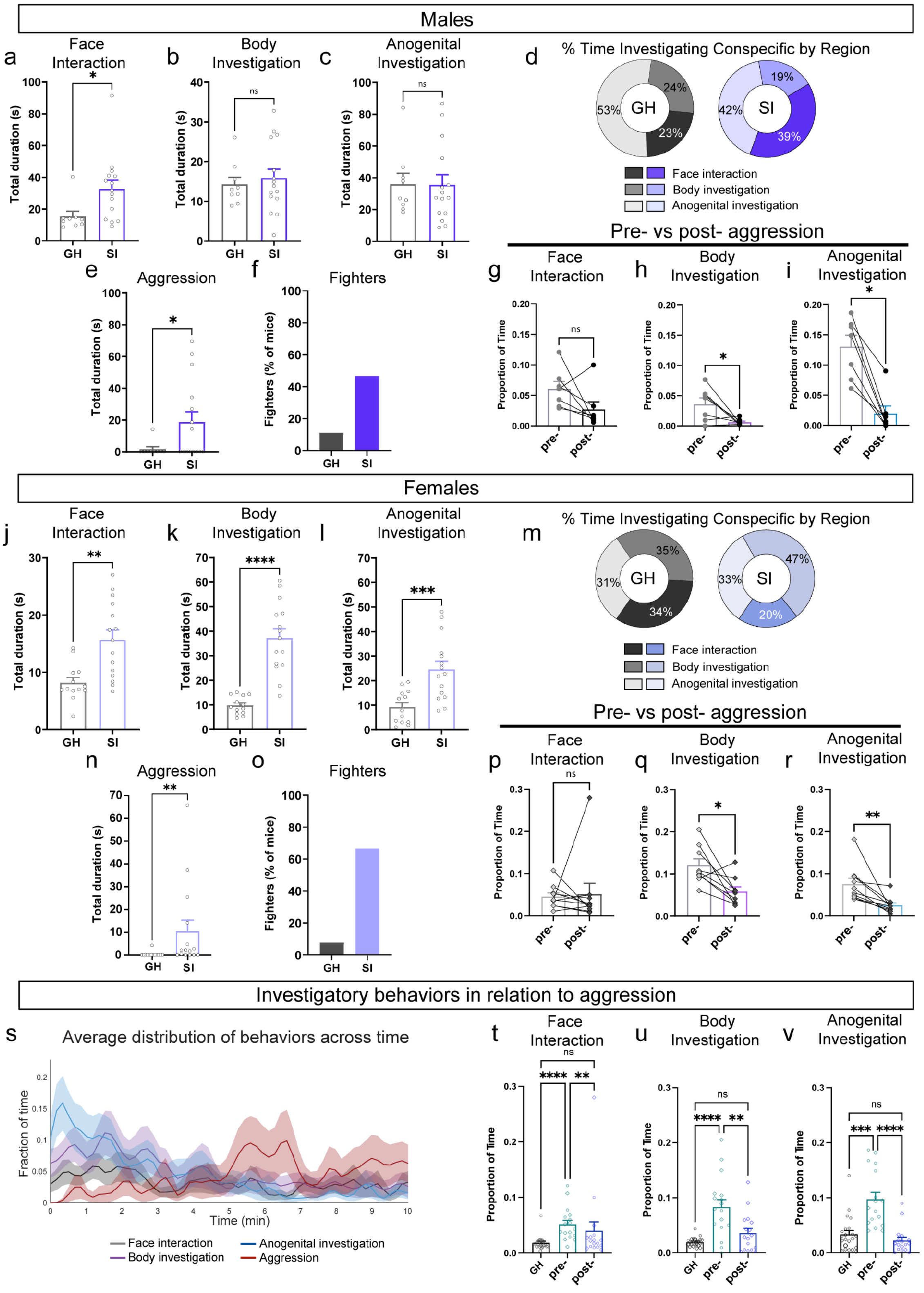


**Extended Data Fig 1. | Prolonged social isolation induces a state of aggression in males and females. a-e,** Males, N = 24 (9 GH, 15 SI). **a**, Total duration (seconds) of face interaction towards intruder for group housed (GH) and socially isolated (SI) males; Mann-Whitney test: p=0.0251. **b**, Total duration (seconds) of body investigation towards intruder for

GH vs SI, unpaired t test p=0.6432, t=0.4697, df=22. **c**, Total duration (seconds) of anogenital investigation towards intruder for GH vs SI, Mann-Whitney test p = 0.7702. **d**, Pie charts showing percent of total investigation time directed towards body region (face, anogenital, or body). **e**, Total duration (seconds) of aggression directed towards intruder for GH vs SI; Mann-Whitney test p=0.0466. **f,** Percent of resident males that engaged in aggressive behavior for GH and SI. **g**, Proportion of total time aggressive isolated males engage in face interaction before (pre) and after (post) the first bout of aggression, Wilcoxon test, p=0.1562. **h**, Proportion of total time engaged in body investigation before and after the first aggressive bout, paired t-test, p=0.0364, t=2.683, df=6. **i**, Proportion of total time engaged in anogenital investigation before and after the first aggressive bout, Wilcoxon test, p=0.0156. **j-n**, Females, N = 28 (13 GH, 15 SI). **j**, Total duration (seconds) of face interaction towards intruder in GH and SI females, unpaired t-test, p=0.0010, t=3.711, df=26. **k**, Total duration (seconds) of body investigation towards intruder for GH vs SI, unpaired t-test, p<0.0001, t=6.730, df=26. **l**, Total duration (seconds) of anogenital investigation towards intruder for GH vs SI, unpaired t-test, p=0.0007, t=3.853, df=26. **m**, Pie charts showing percent of total investigation time directed towards body region (face, anogenital, or body). **n,** Total duration (seconds) of aggression towards intruder for GH vs SI, Mann-Whitney test: p = 0.0023. **o**, Percent of resident females that engaged in aggressive behavior for GH and SI. **p**, Proportion of total time aggressive isolated females engaged in face interaction before (pre) and after (post) the first bout of aggression, Wilcoxon test, p=0.2754. **q**, Proportion of total time engaged in body investigation before and after the first aggressive bout, Wilcoxon test, p=0.0195. **r**, Proportion of total time engaged in anogenital investigation before and after the first bout of aggression, paired t-test, p=0.0068, t=3.497, df=9. **s**, Average temporal distribution of face interaction, body investigation, anogenital investigation and aggression occurring over the course of the entire Resident Intruder assay. **t**, Proportion of time group housed (GH) mice spend engaged in face interaction compared to isolated (SI) mice before (pre) and after (post) the first instance of aggression. Kruskal-Wallis test p < 0.0001, Dunn’s multiple comparisons test: GH vs. SI pre-aggression p<0.0001, GH vs. SI post-aggression p=0.7412, SI pre-aggression vs. SI post-aggression p=0.0085. **u**, Proportion of time spent engaged in body investigation, one-way ANOVA p<0.0001, Tukey's multiple comparisons test: GH vs SI pre-aggression p<0.0001, GH vs. SI post-aggression p=0.3460, SI pre- vs SI post-aggression p=0.0011. **v**, Proportion of time spent engaged in anogenital investigation. Kruskal-Wallis test p<0.0001, Dunn's multiple comparisons test: GH vs SI pre-aggression p=0.0002, GH vs SI post-aggression p=0.7638, SI pre- vs SI post-aggression p<0.0001. *Bars represent mean ± SEM, dots within bars represent mean for each animal. ns p > 0.05, * p < 0.05, ** p < 0.01, *** p < 0.001, **** p < 0.0001.*

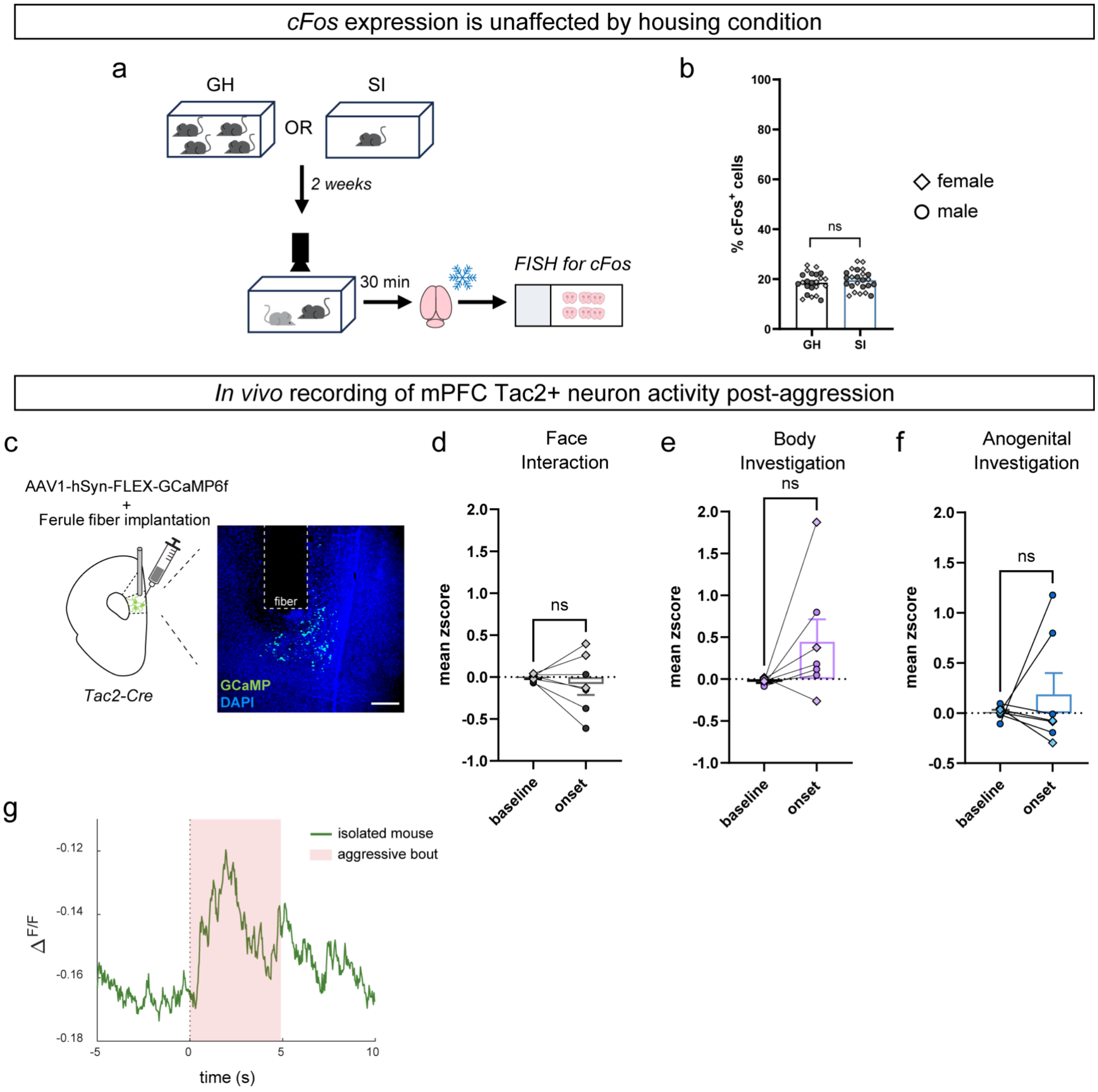


**Extended Data Fig 2. | Activity of mPFC neurons during the Resident Intruder assay.** **a**, Experimental design. Mice were isolated (SI) or group housed (GH) and tested on the Resident Intruder assay. Brains were extracted and processed for FISH to visualize cFos mRNA. **b**, Percent of cFos$^{+}$ cells, N=8 mice (n=4 GH mice, n=4 SI mice), ~3 sections/animal, two-tailed nested t-test, p = 0.6538, t=0.4718, df=6. **c-f**, N = 7 isolated mice (3F, 4M). **c**, Depiction and representative image of viral infusion of Cre-dependent GCaMP (green) and ferule fiber implantation into mPFC of isolated Tac2-IRES-Cre mice. Scale bar represents 50µm. **d**, Mean z-scored dF/F of GCaMP signal from mPFC Tac2$^{+}$ neurons during baseline period (-4 to -2 seconds before behavior onset) and following behavior onset (1 to 3 seconds post onset) for face interaction that occurred post the initial aggressive bout, paired t-test, p=0.5541, t=0.6284, df=6. **e**, Mean z-scored dF/F of GCaMP signal from mPFC Tac2$^{+}$ neurons during baseline period (-4 to -2 seconds before behavior onset) and following behavior onset (1 to 3 seconds post onset) for body investigation that occurred post the initial aggressive bout, paired t test p=0.1263, t=1.775, df=6. **f**, Mean z-scored dF/F of GCaMP signal from mPFC Tac2$^{+}$ neurons during baseline period (-4 to -2 seconds before behavior onset) and following behavior onset (1 to 3 seconds post onset) for anogenital investigation that occurred after the initial aggressive bout, Wilcoxon test, p=0.9375. **g**, Representative dF/F GCaMP trace from mPFC Tac2$^{+}$ neurons during single aggression bout from one mouse. *Bars represent mean ± SEM, individual dots represent mean of individual animals, diamonds=females, circles=males. ns p>0.05.*

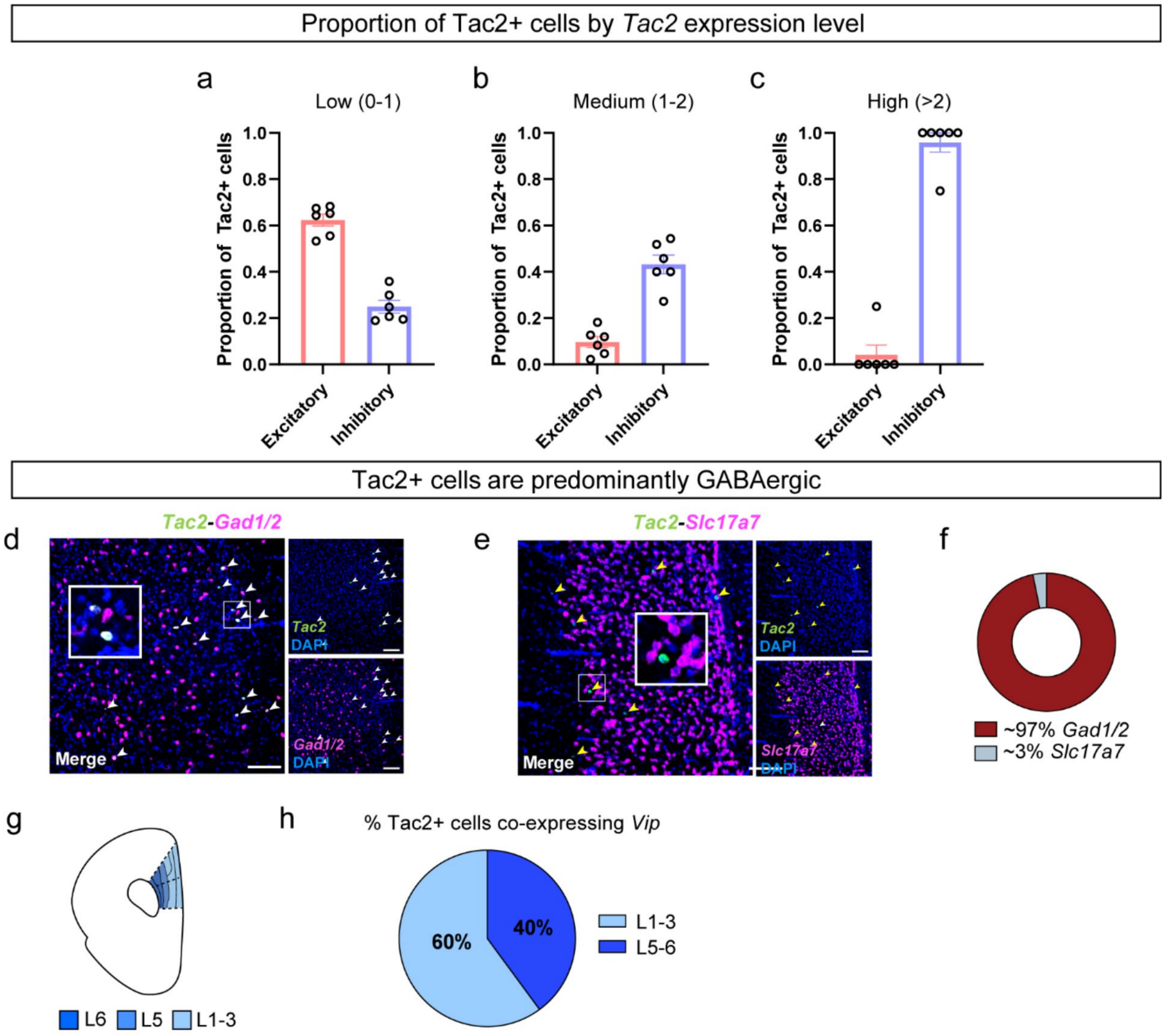


**Extended Data Fig 3. | Cells expressing differing levels of *Tac2* represent a diverse population.** **a**, Proportion of Tac2$^+$ cells expressing low levels of *Tac2* (0-1 normalized copies) belonging to excitatory and inhibitory neuronal cell type clusters. **b**, Proportion of Tac2$^+$ cells expressing medium levels of *Tac2* (1-2 normalized copies) belonging to excitatory and inhibitory neuronal cell type clusters. **c**, Proportion of Tac2$^+$ cells expressing high levels of *Tac2* (>2 normalized copies) belonging to excitatory and inhibitory neuronal cell type clusters. **d**, Representative image of *Tac2* (green) and *Gad1/2* (magenta) co-localization in mPFC using fluorescent in situ hybridization (FISH). White arrows indicate cells with *Tac2* and *Gad1/2* expression, scale bar represents 100µm. **e**, Representative image of *Tac2* (green) and *Slc17a7* (magenta) in mPFC using FISH. Yellow arrows indicate cells with only *Tac2* expression, scale bar represents 100µm. **f**, Proportion of Tac2$^+$ cells that express *Gad1/2* or *Slc17a7*, n = 4 mice, 3 sections/hemisphere/animal. **g**, Diagram of cortical layers in mPFC. **h**, Proportion of Tac2$^+$ cells co-expressing *Vip* broken down by cortical layer: early (L1-3) vs deep (L5-6), n = 4 mice, 3 sections/hemisphere/animal. *Bars represent mean ± SEM, individual dots represent mean of individual animals for sequencing data.*

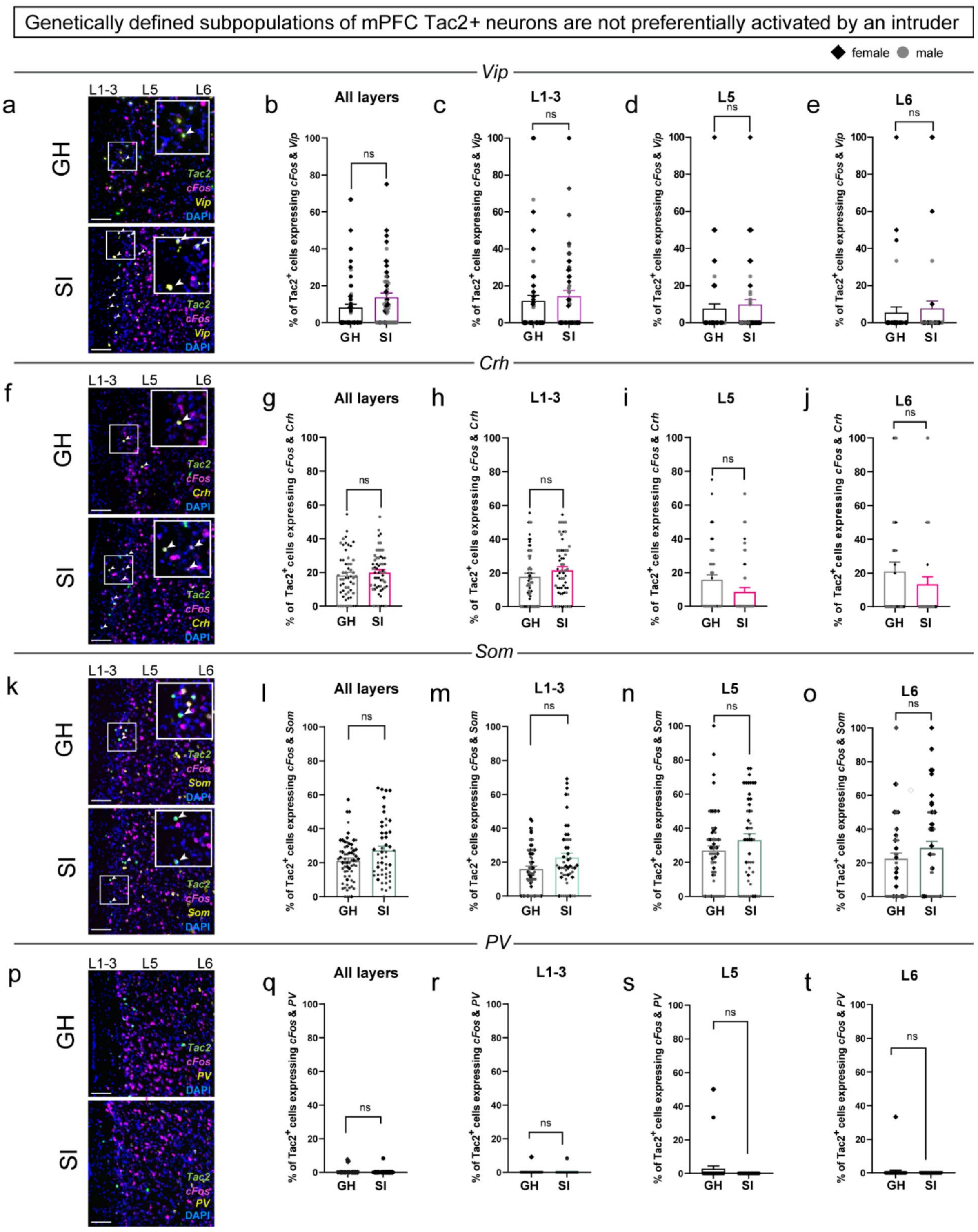


**Extended Data Fig 4. | Fluorescent in situ hybridization labeling of mPFC Tac2⁺ neuron populations activated by the Resident Intruder assay.** **a**, Representative FISH images depicting *Tac2* (green), *cFos* (magenta), and *Vip* (yellow) mRNA expression in group housed (GH) and socially isolated (SI) mice, scale bars represent 100µm. White arrows indicate cells positive for all three markers (*Tac2*, *cFos*, *Vip)*. **b**, Percent of Tac2⁺ neurons expressing both *cFos* and *Vip* in GH vs SI animals, nested t-test p=0.5207, t=0.6819, df=6. **c**, Percent of early layer (L1-3) Tac2⁺ neurons expressing both *cFos* and *Vip*, nested t-test p=0.7647, t=0.3133, df=6. **d**, Percent of layer 5 (L5) Tac2⁺ neurons expressing both *cFos* and *Vip*, nested t-test p=0.7044, t=0.3980, df=6. **e**, Percent of layer 6 (L6) Tac2⁺ neurons expressing both *cFos* and *Vip,* nested t-test p=0.8422, t=0.2079, df=6. **f**, Representative FISH images depicting *Tac2* (green), *cFos* (magenta), and *Crh* (yellow)

mRNA expression in group housed (GH) vs socially isolated (SI) mice, scale bars represent 100µm. White arrows indicate cells positive for all three markers (*Tac2*, *cFos*, *Crh)*. **g**, Percent of $Tac2^+$ neurons expressing both *cFos* and *Crh* in GH vs SI mice, nested t-test p=0.6026, t=0.5493, df=6. **h**, Percent of early layer (L1-3) $Tac2^+$ neurons expressing both *cFos* and *Crh*, nested t-test p=0.3898, t=0.926, df=6. **i**, Percent of layer 5 (L5) $Tac2^+$ neurons expressing both *cFos* and *Crh*, nested t-test p=0.1202, t=1.811, df=6. **j**, Percent of layer 6 (L6) $Tac2^+$ neurons expressing both *cFos* and *Crh,* nested t-test p=0.3077, t=1.114, df=6. **k**, Representative FISH images depicting *Tac2* (green), *cFos* (magenta), and *Som* (yellow) mRNA expression in group housed (GH) vs socially isolated (SI) mice, scale bars represent 100µm. White arrows indicate cells positive for all three markers (*Tac2*, *cFos*, *Som)*. **l**, Percent of $Tac2^+$ neurons expressing both *cFos* and *Som* in GH vs SI mice, nested t-test p=0.5437, t=0.6435, df=6. **m**, Percent of early layer (L1-3) $Tac2^+$ neurons expressing both *cFos* and *Som*, nested t-test p=0.3116. **n**, Percent of layer 5 (L5) $Tac2^+$ neurons expressing both *cFos* and *Som*, nested t-test p=0.6971, t=0.4085, df=6. **o**, Percent of layer 6 (L6) $Tac2^+$ neurons expressing both *cFos* and *Som,* nested t-test p=0.7094, t=0.3908, df=6. **p**, Representative images of FISH for *Tac2* (green), *cFos* (magenta), and *PV* (yellow) mRNA expression in group housed (GH) vs socially isolated (SI) mice, scale bars represent 100µm. **q**, Percent of $Tac2^+$ neurons expressing both *cFos* and *PV* in GH vs SI mice, nested t-test p=0.4175, t=0.8705, df=6. **r**, Percent of early layer (L1-3) $Tac2^+$ neurons expressing both *cFos* and *PV*, nested t-test, p=0.9947, t=0.006889, df=6. **s**, Percent of layer 5 (L5) $Tac2^+$ neurons expressing both *cFos* and *PV*, nested t-test p=0.1839, t=1.502, df=6. **t**, Percent of layer 6 (L6) $Tac2^+$ neurons expressing both *cFos* and *PV,* nested t-test p=0.3559, t=1.00, df=6. N = 8 mice (n = 4 animals/housing condition, ~4 sections/side per animal. *Bars represent mean ± SEM, individual dots represent mean of individual animals for behavioral data and individual mPFC tissue sections/hemisphere for FISH data, diamonds = females, circles = males. ns p > 0.05.*

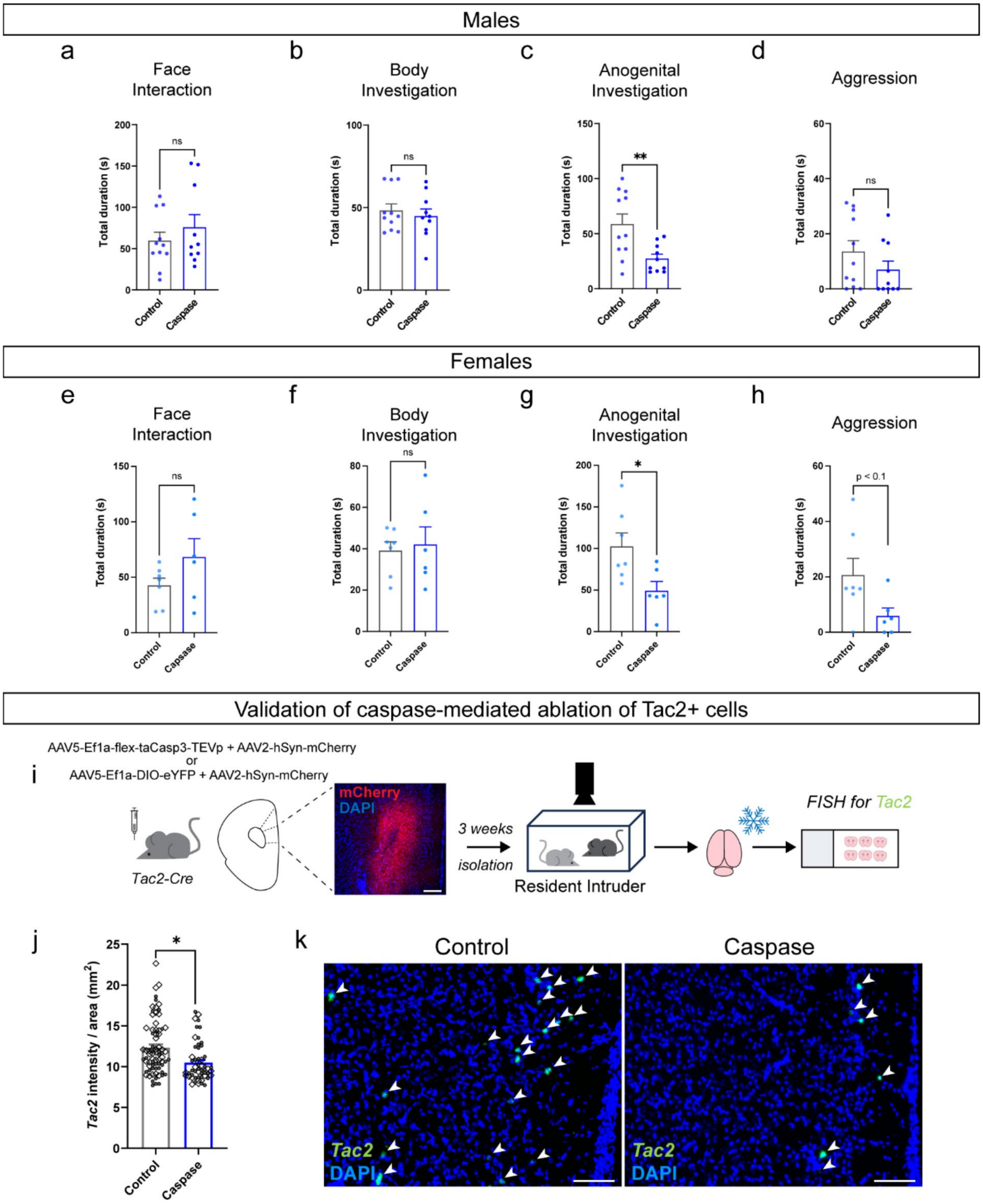

**Extended Data Fig 5. | Caspase-mediated ablation of mPFC Tac2⁺ neurons attenuates isolation-induced investigation and aggression.** **a**-**d**, Males, n = 11 control, n = 10 caspase. **a**, Total duration (seconds) of face interaction in mice expressing control eYFP virus or caspase virus in mPFC Tac2⁺ neurons, Mann-Whitney test, p=0.7045. **b**, Total duration (seconds) of body investigation in mice expressing control eYFP virus or caspase virus in mPFC Tac2⁺ neurons, Mann-Whitney test, p=0.7045. **c**, Total duration (seconds) of anogenital investigation in mice expressing control eYFP virus or caspase virus in mPFC Tac2⁺ neurons, unpaired t-test, p=0.0068, t=3.3038, df=19. **d**, Total duration (seconds) of aggression in mice expressing control eYFP virus or caspase virus in mPFC Tac2⁺ neurons, Mann-Whitney test, p=0.1649. **e**-**h**, Females, n = 7 control, n = 6 caspase. **e**, Total duration (seconds) of face interaction in mice expressing control eYFP virus or caspase virus in mPFC Tac2⁺ neurons, unpaired t-test p=0.1535, t=1.533, df=11. **f**, Total duration (seconds) of body investigation in mice expressing control eYFP virus or caspase virus in mPFC Tac2⁺ neurons, unpaired t-test p=0.7485, t= 0.3287, df=11. **g**, Total duration (seconds) of anogenital investigation in mice expressing control eYFP virus or caspase virus in mPFC Tac2$^+$ neurons, unpaired t-test p=0.0234, t=2.631, df=11. **h**, Total duration (seconds) of aggression in mice expressing control eYFP virus or caspase virus in mPFC Tac2$^+$ neurons, Mann-Whitney test, p=0.0997. **i**, Experimental design for behavior and brain extraction in preparation for fluorescent in situ hybridization (FISH) to validate ablation of mPFC Tac2$^+$ neurons. **j**, Fluorescence intensity of *Tac2* expression normalized by total area imaged, nested t-test p=0.0284, t=2.362, df=20, n = 11 control, n = 11 caspase (mix of sexes per viral condition), ~2 sections/hemisphere per animal. **k**, Representative image of FISH for *Tac2* in control animal (left) and caspase animal (right). Scale bars represent 100µm. White arrows indicate Tac2$^+$ cells. *Bars represent mean ± SEM, individual dots represent mean of individual animals for behavioral data and individual mPFC tissue sections/hemisphere for FISH data, diamonds = females, circles = males. ns p > 0.05, * p < 0.05, ** p<0.01.*

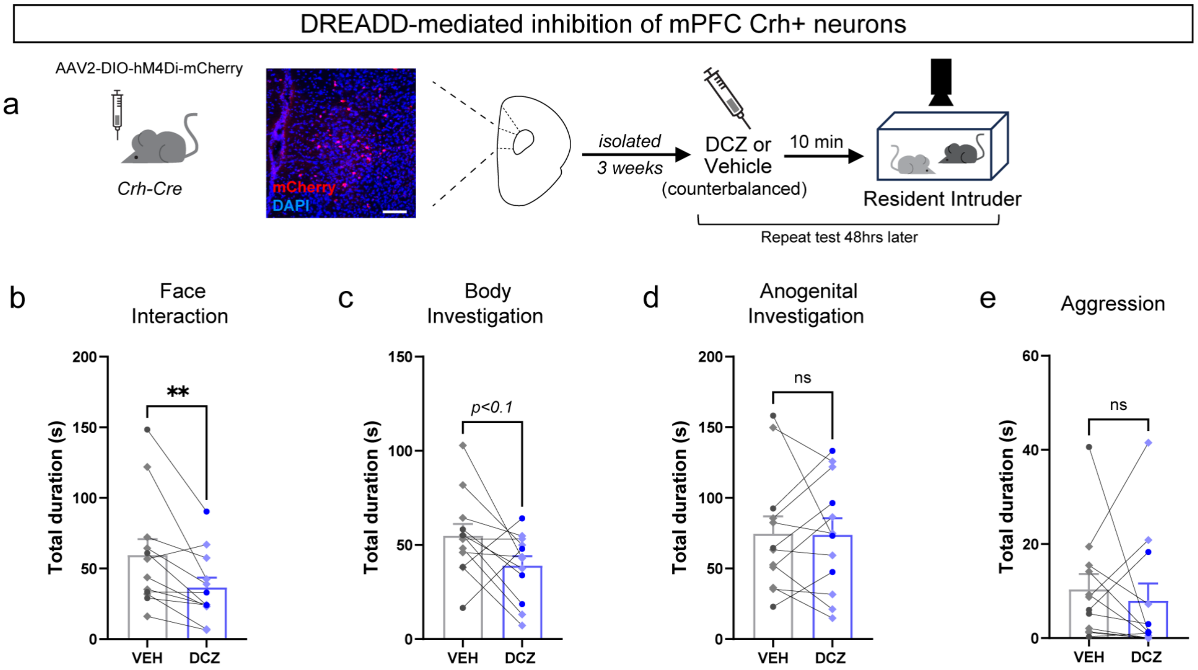


**Extended Data Fig 6. | DREADD-mediated inhibition of mPFC Crh$^+$ cells has no impact on isolation-induced aggression or anogenital investigation. a**, Experimental design depicting Cre-dependent, virally-mediated injection of hM4Di into the mPFC of Crh-IRES-Cre male and female mice. Mice were isolated and tested in the Resident Intruder assay to understand whether mPFC Crh$^+$ neurons contribute to the aggressive state induced by isolation. Prior to testing, mice were injected with vehicle (VEH, controls) or Deschloroclozapine (DCZ), for DREADD-mediated inhibition of mPFC Crh$^+$ neurons. Mice were tested twice using a within-subjects design: once under the influence of DCZ, once under the influence of vehicle (order counterbalanced). Representative image of viral infusion (red) in the mPFC, scale bar represents 100µm **b-e**, N=12 mice. **b**, Total duration (seconds) of face interaction, paired t-test, p=0.0090, t=3.165, df=11. **c**, Total duration (seconds) of body investigation, paired t-test p=0.0620, t=2.077, df=11. **d**, Total duration (seconds) of anogenital investigation, paired t-test p=0.9506, t=0.0634, df=11. **e**, Total duration (seconds) of aggression, Wilcoxon test, p=0.4697. *Bars represent mean ± SEM, individual dots represent mean of individual animals, diamonds = females, circles = males. ns p > 0.05, ** p<0.01.*

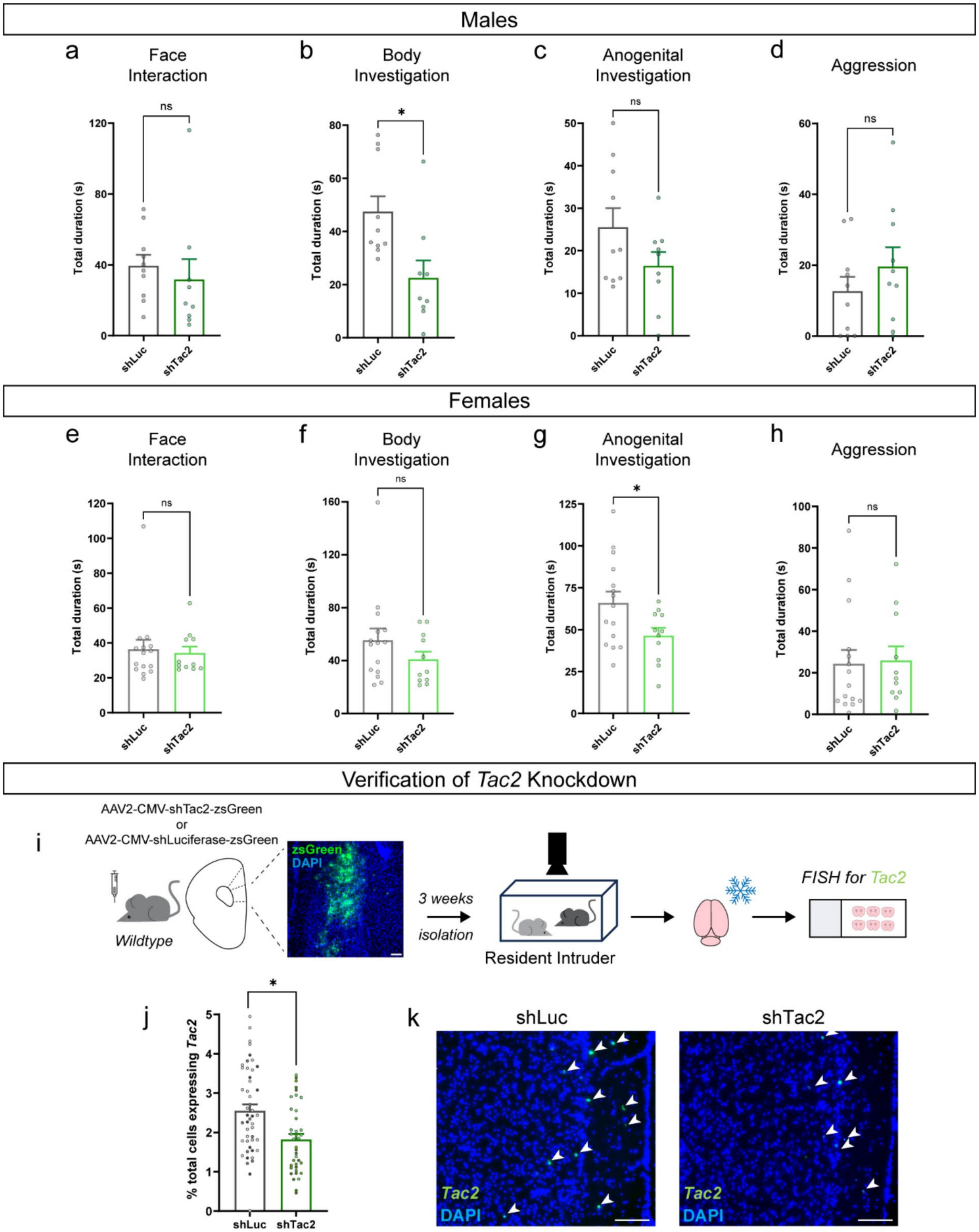


**Extended Data Fig 7. | *Tac2* knockdown in mPFC attenuates isolation-induced increases in investigation. a-d**, Males, n = 10 shLuc, n = 9 shTac2. **a**, Total duration (seconds) of face interaction in animals expressing shRNA targeting Luciferase (shLuc) or *Tac2* (shTac2) in the mPFC, Mann-Whitney test: p = 0.1564. **b**, Total duration (seconds) of body investigation in animals expressing shRNA targeting Luciferase (shLuc) or *Tac2* (shTac2) in the mPFC, unpaired t test: p = 0.0108, t=2.861, df=17. **c**, Total duration (seconds) of anogenital investigation in animals expressing shRNA targeting Luciferase (shLuc) or *Tac2* (shTac2) in the mPFC, unpaired t test: p = 0.1271, t=1.604, df=17. **d**, Total duration (seconds) of aggressive behavior in animals expressing shRNA targeting Luciferase (shLuc) or *Tac2* (shTac2) in the mPFC, unpaired t test: p = 0.3172, t=1.029, df=18. **e-h**, Females, n = 15 shLuc, n = 11 shTac2. **e**, Total duration (seconds) of face interaction in animals expressing shRNA targeting Luciferase (shLuc) or *Tac2* (shTac2) in the mPFC, Mann-Whitney test, $p = 0.8785$. **f**, Total duration (seconds) of body investigation in animals expressing shRNA targeting Luciferase (shLuc) or *Tac2* (shTac2) in the mPFC, unpaired t-test $p = 0.2199$, $t = 1.260$, $df = 24$. **g**, Total duration (seconds) of anogenital investigation in animals expressing shRNA targeting Luciferase (shLuc) or *Tac2* (shTac2) in the mPFC, unpaired t-test $p = 0.0384$, $t = 2.191$, $df = 24$. **h**, Total duration (seconds) of aggression in animals expressing shRNA targeting Luciferase (shLuc) or *Tac2* (shTac2) in the mPFC, unpaired t-test $p = 0.8706$, $t = 0.1646$, $df = 24$. **i**, Representative image of viral infusion site (green) in mPFC, scale bar represents 100μm. Experimental design depicting virally mediated shRNA infusions into the mPFC of isolated wildtype mice, subsequent behavior testing and fluorescent in situ hybridization (FISH) verification of *Tac2* knockdown. **j**, Percent of total cells ($DAPI^+$, blue) expressing *Tac2* (green), verifying *Tac2* knockdown in mPFC of shTac2 mice, nested t-test, $p = 0.0448$, $t = 2.662$, $df = 5$. N = 7 mice for FISH analyses (4 shLuc, 3 shTac2 across sexes), ~ 3 sections/hemisphere per animal. **k,** Representative images of FISH for *Tac2* expression in shLuc (left) or shTac2 (right) mice. Scale bars represent 50μm**.** White arrows indicate $Tac2^+$ cells. *Bars represent mean ± SEM, individual dots represent mean of individual animals for behavioral data and individual sections for FISH data; ns $p > 0.05$, * $p < 0.05$.*

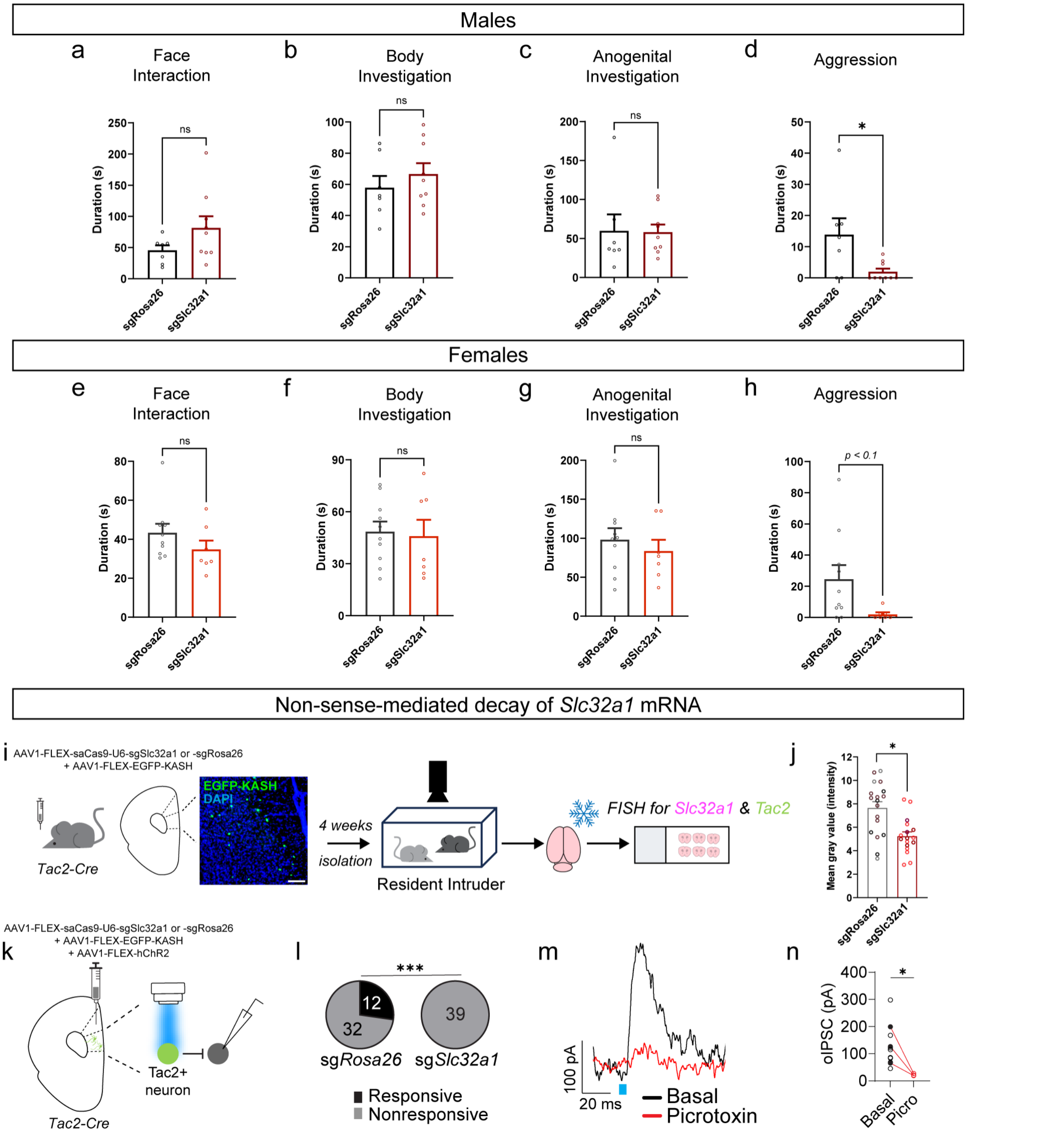


**Extended Data Fig 8. | *Slc32a1* missense mutations in mPFC Tac2⁺ neurons attenuates isolation-induced aggression.** **a**-**d**, Males, n = 7 sgRosa26, n = 9 sgSlc32a1. **a**, Total duration (seconds) of face interaction in mice with CRISPR/SaCas9 virus directed to the Rosa26 locus (sgRosa26, controls) or *Slc32a1* (sgSlc32a1) in the mPFC, unpaired t-test, p = 0.1321, t = 1.599, df = 14. **b**, Total duration (seconds) of body investigation in sgRosa26 vs sgSlc32a1 mice, unpaired t-test, p = 0.4110. **c**, Total duration (seconds) of anogenital investigation in sgRosa26 vs sgSlc32a1 mice, Mann-Whitney test, p = 0.6065. **d**, Total duration (seconds) of aggression in sgRosa26 vs sgSlc32a1 mice, Mann-Whitney test, p = 0.0273. **e**-**h**, Females, n = 10 sgRosa26, n = 7 sgSlc32a1. **e**, Total duration (seconds) of face interaction in mice with CRISPR/SaCas9 virus directed to the Rosa26 locus (sgRosa26, controls) or *Slc32a1* (sgSlc32a1) in the mPFC, unpaired t-test, p = 0.2157, t = 1.292, df = 15. **f**, Total duration (seconds) of body investigation in sgRosa26 vs sgSlc32a1 mice, unpaired t-test p = 0.8129, t = 0.2408, df = 15. **g**, Total duration (seconds) of anogenital investigation in sgRosa26 vs sgSlc32a1 mice, unpaired t-test, p = 0.5041, t = 0.6845, df = 15. **h**, Total duration (seconds) of aggression in sgRosa26 vs sgSlc32a1 mice, unpaired t-test, p = 0.0582, t = 2.505, df = 15. **i**, Experimental design depicting virally-mediated

CRISPR/SaCas9-sgSlc32a1 or CRISPR/SaCas9-sgRosa26 and KASH-eGFP infusions into the mPFC of Tac2-IRES-Cre mice, followed by behavior testing, and fluorescent in situ hybridization (FISH) on mPFC sections from sgRosa26 and sgSlc32a1 mice to confirm nonsense mediated decay of *Slc32a1* in Tac2$^+$ neurons. Representative image of viral infusion site (green) in mPFC, scale bar represents 100µm. **j**, Fluorescence intensity (mean grey value) of *Slc32a1* fluorescently labeled mRNA in a mask of Tac2$^+$ cells from FISH analyses (RNAscope™), N = 6 animals for FISH (3 per viral condition, mix of sexes), ~ 4 sections per animal, nested t-test $p = 0.0215$, $t = 3.665$, $df = 4$. **k**, Validation of disrupted GABA release from mPFC Tac2$^+$ neurons expressing sg*Slc32a1* construct. Tac2-Cre mice were injected with a Cre-dependent ChR2 in combination with the CRISPR/SaCas9 virus directed to *Rosa26* (controls) or directed to *Slc32a*1. Brain slices were prepared for electrophysiological recordings. Recordings were obtained from Tac2$^-$ cells that receive inhibitory synaptic inputs from Tac2$^+$ neurons following optogenetic stimulation via ChR2. **l**, Proportion of responsive cells for sgRosa26- vs sgSlc32a1-injected mice, Fisher's exact test, $p=0.0003$, $n=44$ cells from 4 sg*Rosa26* mice, and $n=39$ from 3 sg*Slc32a1* mice. **m**, Representative traces of optogenetically evoked IPSCs following bath application of picrotoxin vs basal condition. **n**, IPSC amplitudes for basal conditions vs. following picrotoxin application, unpaired t-test, $p=0.0485$. *Bars are mean ± SEM, individual dots for behavioral graphs represent mean duration for an individual animal, individual sections for FISH data, and individual cells for electrophysiological data. ns $p > 0.05$, * $p < 0.05$, *** $p < 0.001$.*

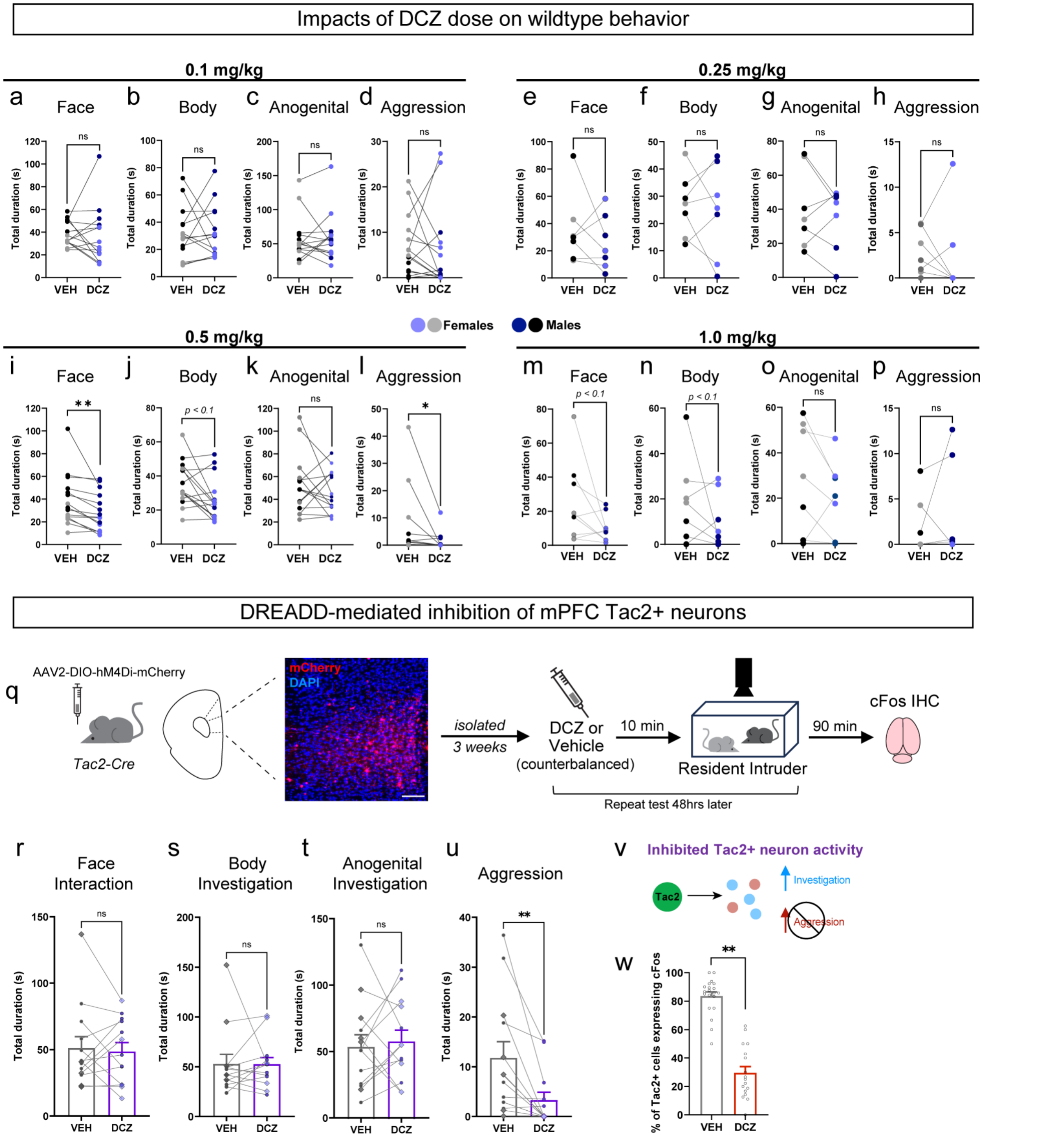


**Extended Data Fig 9. | Acute DREADD-mediated inhibition of mPFC Tac2$^{+}$ neurons attenuates isolation-induced aggression but not investigation.** **a**-**p**, Experiments testing dose-dependent effects of DCZ in wildtype, isolated male and female mice. **a-d**, 0.1 mg/kg DCZ; n = 14 mice. **a**, Total duration (seconds) of face interaction for isolated animals that received VEH or 0.1 mg/kg DCZ; unpaired t-test p=0.7161, t=0.3717, df=13. **b**, Total duration (seconds) of body investigation; paired t test: 0.9307, t = 0.08870, df = 13. **c**, Total duration (seconds) of anogenital investigation; Wilcoxon test: p = >0.9999. **d**, Total duration (seconds) of aggression; Wilcoxon test: p = 0.4263. **e-h**, 0.25 mg/kg DCZ, n = 7. **e**, Total duration (seconds) of face interaction for isolated animals that received VEH or 0.25mg/kg DCZ; paired t-test p = 0.4242, t = 0.8573, df = 6. **f**, Total duration (seconds) of body investigation; paired t-test p = 0.7299, t = 0.3618, df = 6. **g**, Total duration (seconds) of anogenital investigation, paired t-test p = 0.4744, t=0.7629, df=6. **h**, Total duration (seconds) of aggression; Wilcoxon test p = 0.8438. **i-l,** 0.5 mg/kg DCZ, n = 15. **i-l,** 0.5 mg/kg DCZ, n = 15. **i,** Total duration (seconds) of face interaction for isolated animals that received VEH or 0.5 mg/kg DCZ, paired t-test p = 0.0063, t = 3.207, df = 14. **j**, Total duration (seconds) of body investigation, Wilcoxon test p = 0.0833. **k**, Total duration (seconds) of anogenital investigation, paired t-test p = 0.4595, t = 0.7606, df = 14. **l**, Total duration (seconds) of aggression, Wilcoxon test, p =

0.0215. **m-p,** 1.0 mg/kg DCZ; n = 9 mice. **m**, Total duration (seconds) of face interaction for isolated animals that received VEH or 1.0 mg/kg DCZ, paired t-test p = 0.0918, t = 1.915, df = 8. **n**, Total duration (seconds) of body investigation, paired t-test p = 0.3200, t = 1.060, df = 8. **o,** Total duration (seconds) of anogenital investigation, paired t-test p = 0.1896, t = 1.433, df = 8. **p**, Total duration (seconds) of aggression, Wilcoxon test, p = 0.4375. *Individual dots with connecting lines show the mean duration of behavior for each animal on each drug, lighter colors represent females, darker colors males.* **q**, Experimental design depicting Cre-dependent, virally-mediated injection of hM4Di into the mPFC of Tac2-IRES-Cre male and female mice and subsequent behavioral testing following isolation. Prior to testing, mice were injected with vehicle (VEH, controls) or Deschloroclozapine (DCZ), for DREADD-mediated inhibition of mPFC Tac2$^+$ neurons. Mice were tested twice using a within-subjects design: once under the influence of DCZ, once under the influence of vehicle (order counterbalanced). Representative image of viral infusion (red) in the mPFC, scale bar represents 100µm. **r-u**, n = 13 mice. **r**, Total duration (seconds) of face interaction, Wilcoxon test: p = 0.8926. **s**, Total duration (seconds) of body investigation, Wilcoxon test: p = 0.541. **t**, Total duration (seconds) of anogenital investigation, paired t-test: p = 0.6932, t=0.4042, df=12. **u**, Total duration (seconds) of aggression, Wilcoxon test: p = 0.0012. *Individual dots with connecting lines show the mean of individual animals' behavior duration on each drug, diamonds represent females, circles males.* **v**, Model depicting the impact of acute mPFC Tac2$^+$ neuron silencing to reduce aggression but not investigation. **w**, Percent of Tac2$^+$ neurons expressing cFos in hM4Di animals given VEH or DCZ to validate cell silencing, N=8 (4 VEH, 4 DCZ), nested t-test, p=0.0012, t=5.763, df=6. *Bars represent mean ± SEM, individual dots represent mean of individual animals for behavioral data and individual sections for FISH data. ns p > 0.05, * p < 0.05, ** p < 0.01.*

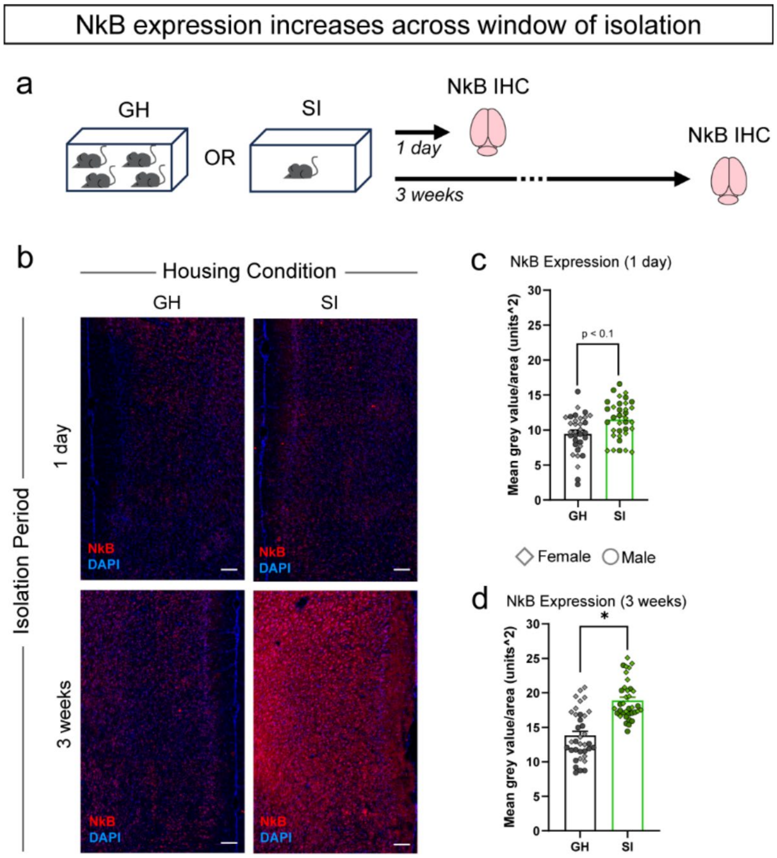


**Extended Data Fig 10. | Neurokinin B expression in the mPFC is increased by social isolation.** **a**, Experimental design for NkB immunohistochemistry (IHC) after 1 day or 3 weeks of isolation. **b**, Representative image of NkB immunostaining (red) in a group housed mouse after 1 day of group-housing (upper left). Representative image of NkB immunostaining (red) after 1 day of social isolation (upper right). Representative image of NkB immunostaining (red) in a group housed mouse after 3 weeks of group-housing (lower left). Representative image of NkB immunostaining (red) in an animal single housed for 3 weeks (lower right). **c**, Mean intensity of NkB expression in group housed (GH) vs socially isolated (SI) mice at 1 day, nested-test p=0.0894, t=1.881, df=10, n=12 mice (6 GH, 6 SI, ~3 sections/animal). **d**, Mean intensity of NkB expression in group housed (GH) vs socially isolated (SI) mice at 3 weeks, nested t-test p=0.0116, t=3.082, df=10, n=12 mice (6 GH, 6 SI, ~3 sections/animal). Scale bars represent 100µm. *Bars represent mean±SEM, dots represent mean intensity/section/hemisphere, diamonds = females, circles = males. ns p>0.05, *p<0.05.*

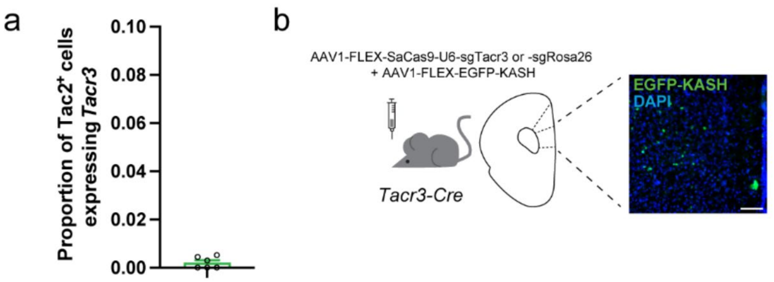


**Extended Data Fig 11. | *Tacr3* expression in mPFC Tac2+ cells and strategy for Cre-dependent Tacr3 mutagenesis.** **a**, Proportion of mPFC Tac2$^{+}$ cells expressing *Tacr3*, N=6 mice. **b**, Depiction of injection cocktail into the mPFC of Tacr3-Cre mice for Cre-dependent CRISPR-mediated nonsense mutations in *Tacr3* (sgTacr3) vs control (sgRosa26) mice. Representative image (right) of viral injection site (green), scale bar represents 100µm. *Bars represent mean ± SEM, individual dots represent mean of individual animals for sequencing data.*

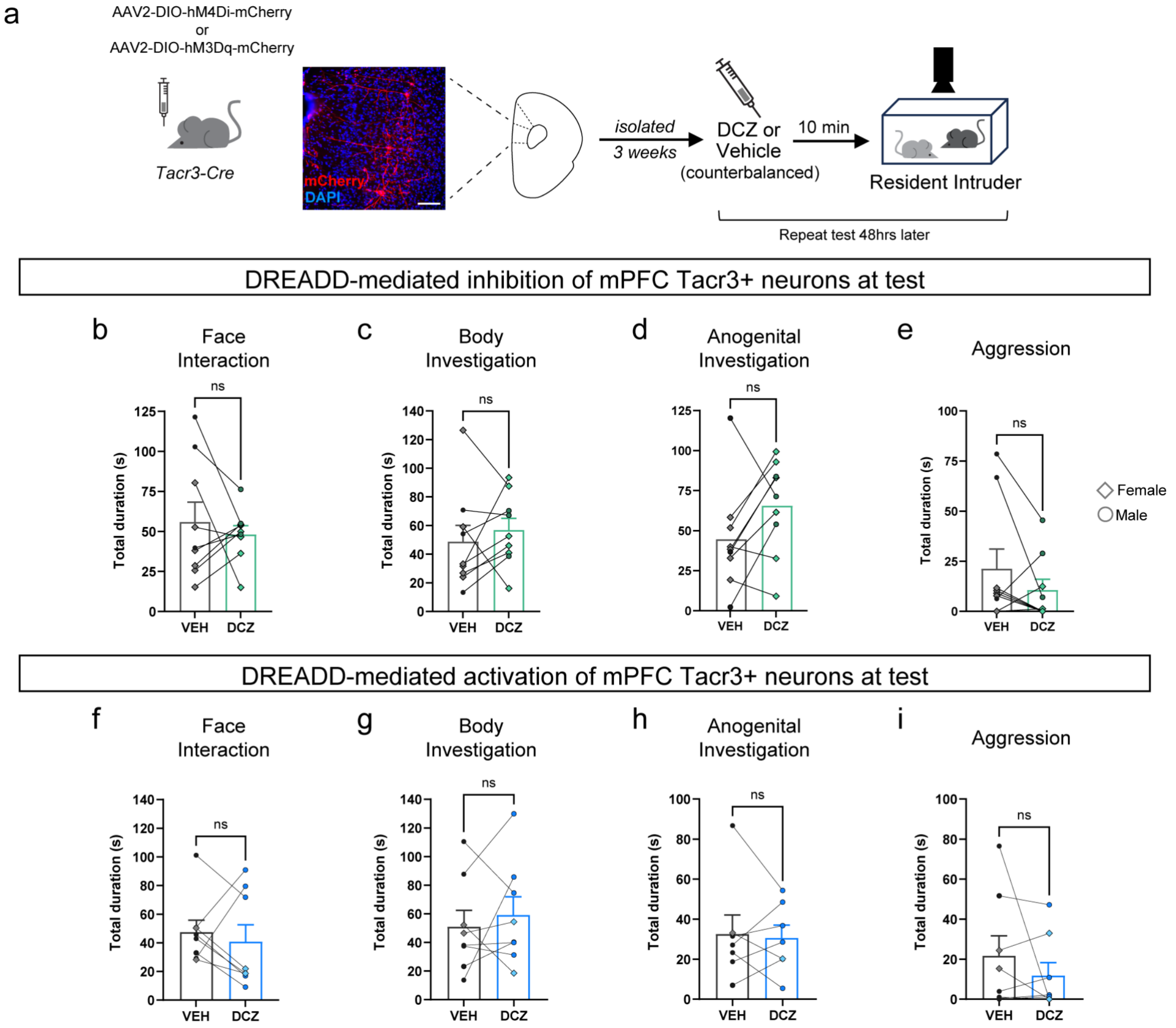


**Extended Data Fig 12. | Acute manipulation of Tacr3$^{+}$ neuron activity is insufficient to reverse isolation-induced behavioral changes.** **a**, Experimental design for DREADD-mediated, acute inhibition or activation of mPFC Tacr3$^{+}$ neurons in isolated male and female mice at test. Tacr3-Cre mice were injected with a virally-mediated hM4Di (inhibition) or hM3Dq (activation), isolated, and administered vehicle (VEH) or deschloroclozapine (DCZ). All mice were tested twice using a within-subjects design: once under the influence of VEH, once under the influence of DCZ (test order counterbalanced). Scale bar represents 100µm. **b-e**, DREADD-mediated inhibition of Tacr3$^{+}$ neurons at test, n = 9 mice. **b**, Total duration (seconds) of face interaction, Wilcoxon test, p=0.3594. **c**, Total duration (seconds) of body investigation, paired t-test p=0.4778, t=0.7446, df=8. **d**, Total duration (seconds) of anogenital investigation, paired t-test p=0.1115, t=1.788, df=8. **e**, Total duration (seconds) of aggression, Wilcoxon test, p=0.3594. **f-i**, DREADD-mediated activation of mPFC Tacr3⁺ neurons at test, n=8 animals. **f**, Total duration (seconds) of face interaction, Wilcoxon test, p=0.7422. **g**, Total duration (seconds) of body investigation, paired t-test, p=0.5434, t=0.6387, df=7. **h**, Total duration (seconds) of anogenital investigation, Wilcoxon test, p=0.9375. **i**, Total duration (seconds) of aggression, Wilcoxon test, p=0.8438. *Bars represent mean ± SEM, individual dots represent mean of individual animals for behavioral data, diamonds = females, circles = male; ns p > 0.05.*

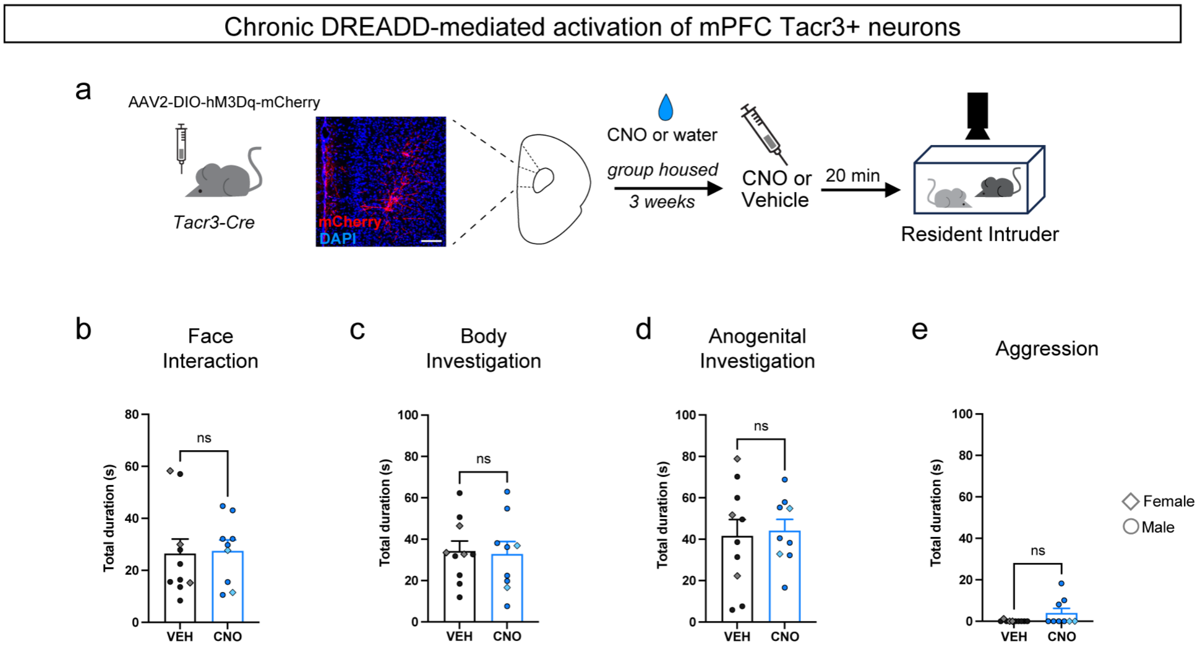


**Extended Data Fig 13. | Chronic activation of mPFC Tacr3⁺ neurons in group-housed mice is insufficient to induce an isolation-like state. a**, Experimental design for DREADD-mediated, chronic activation of mPFC Tacr3⁺ neurons in group-housed male and female mice. Tacr3-Cre mice received mPFC injections of a virally-mediated hM3Dq (activation) and were administered vehicle (VEH) or clozapine-N-oxide (CNO) in their drinking water across a 3-week period to chronically activate mPFC Tacr3⁺ neurons (across a fictive window of social isolation). All mice were given an additional injection of VEH or CNO (concordant with what they received in their drinking water) prior to test, n = 9 vehicle, n = 10 CNO. Scale bar represents 100μm**. b**, Total duration (seconds) of face interaction, unpaired t-test p=0.8931 t=0.1364, df=17. **c**, Total duration (seconds) of body investigation, unpaired t-test p=0.8465, t=0.1965, df=17. **d**, Total duration (seconds) of anogenital investigation, unpaired t test p=0.7991, t=0.2586, df=17. **e**, Total duration (seconds) of aggression, Mann-Whitney test, p=0.1409. *Bars represent mean±SEM, diamonds (females) and circles (males) represent mean behavior duration for each animal. ns p > 0.05.*
